# CELLISA – a cell-cell binding assay for evaluation of nanovesicle targeting proteins

**DOI:** 10.64898/2026.04.09.717595

**Authors:** Taylor F. Gunnels, Jonathan D. Boucher, Yazeed S. Alroogi, Neha P. Kamat, Joshua N. Leonard

## Abstract

Enhancing targeted delivery of biomedicines improves efficacy and can reduce off-target effects by lowering the effective dose, but achieving targeting is challenging. Extracellular vesicles (EVs) are promising biological nanovesicles which can be targeted by displaying binding proteins and are being developed as therapeutics. Currently, discovering EV targeting constructs is limited by low throughput and resource-intensive EV production and isolation. To accelerate discovery, we developed a screening pipeline to identify EV targeting constructs without requiring EV production. This approach is premised on the hypothesis that cell-cell interactions may predict some cell-EV interactions. Our <u>cell</u> binding as<u>say</u> (CELLISA) quantifies binding of a cell surface-displayed targeting protein to its cognate receptor on a target cell, employing a microscopy-based analysis pipeline. After validating the premise using existing T cell-targeting reagents, we develop CELLISA for either adherent or suspension EV producer cells. Finally, we use CELLISA to evaluate new binders and validate that hits mediate targeting and/or delivery of genetic cargo to natural killer cells and T cells. CELLISA increased throughput > 6-fold and decreased time by 40% compared to standard EV screens, and it identified a T-cell binder conferring efficient gene delivery. CELLISA is easily adaptable to other laboratories and can accelerate EV research.

## Introduction

Targeting therapeutic biomolecules to their intended sites of action is a wide-reaching goal in genetic medicine and drug delivery. Bioactive cargos such as siRNA [1, 2] or CRISPR-based prime editors [3] can be packaged into nanoparticles to protect them during delivery and confer targeting to specific cell types, for example, by displaying antibody fragments on the nanoparticle that bind to protein markers on the target cells’ surfaces [4–7]. Extracellular vesicles (EVs) are an appealing class of such nanocarriers—these lipid bilayer vesicles natively transport protein and nucleic acid cargo between cells [8, 9], are well tolerated *in vivo* [10–12], and have been successfully retargeted using surface-displayed antibody fragments [4, 5]. Strategies that enable robust EV targeting can improve delivery of therapeutics to intended cells, lower effective doses required, and by doing so reduce off-target effects.

A key challenge when engineering EV properties such as targeting is that EV production of multiple variants is typically low throughput and time-consuming. This restriction on throughput is particularly limiting for combinatorial design problems such as targeting, in which one must evaluate choices that impact function in pronounced and interacting ways, including: 1) surface marker and epitope, 2) affinity reagent (e.g., antibody-derived fragment, nanobody, de novo designed binder, etc.), 3) transmembrane anchor/display system, and 4) linker or “scaffold” sequence between the affinity reagent and the anchor (**Figure 1A**). Exploring this design space is challenging because production of engineered EVs is limited by low throughput purification processes (e.g., differential ultracentrifugation or sequential tangential flow filtration and size exclusion chromatography [13]) and characterization methodologies often needed for normalization (e.g., particle counting by nanoparticle tracking analysis [14] or other methods) (**Figure 1B).** To provide a sense of scale, for a single researcher, evaluating ∼five binding constructs plus a non-targeted control typically saturates the capacity of differential ultracentrifugation and can take several days of work for EV purification and characterization alone. While pooled analyses facilitate the scale-up of other discovery/engineering methods (e.g., lentiviral libraries to screen protein variants [6] for the function they confer on recipient cells), this is not readily applied to the EV targeting problem; the “phenotype” of the EV is conferred by its surface, necessitating that each cell produce only one targeting construct, and physically linking that EV phenotype to its corresponding genotype (in the EV producer cell) is nontrivial. Thus, unsurprisingly, EV targeting studies typically evaluate only a few constructs (e.g., three [5], four [4], five [15–17], six [18], or seven constructs [19]). The ability to scale up the evaluation of possible designs would enable greater sampling of the large design space and drive better EV targeting.

**Figure 1.**
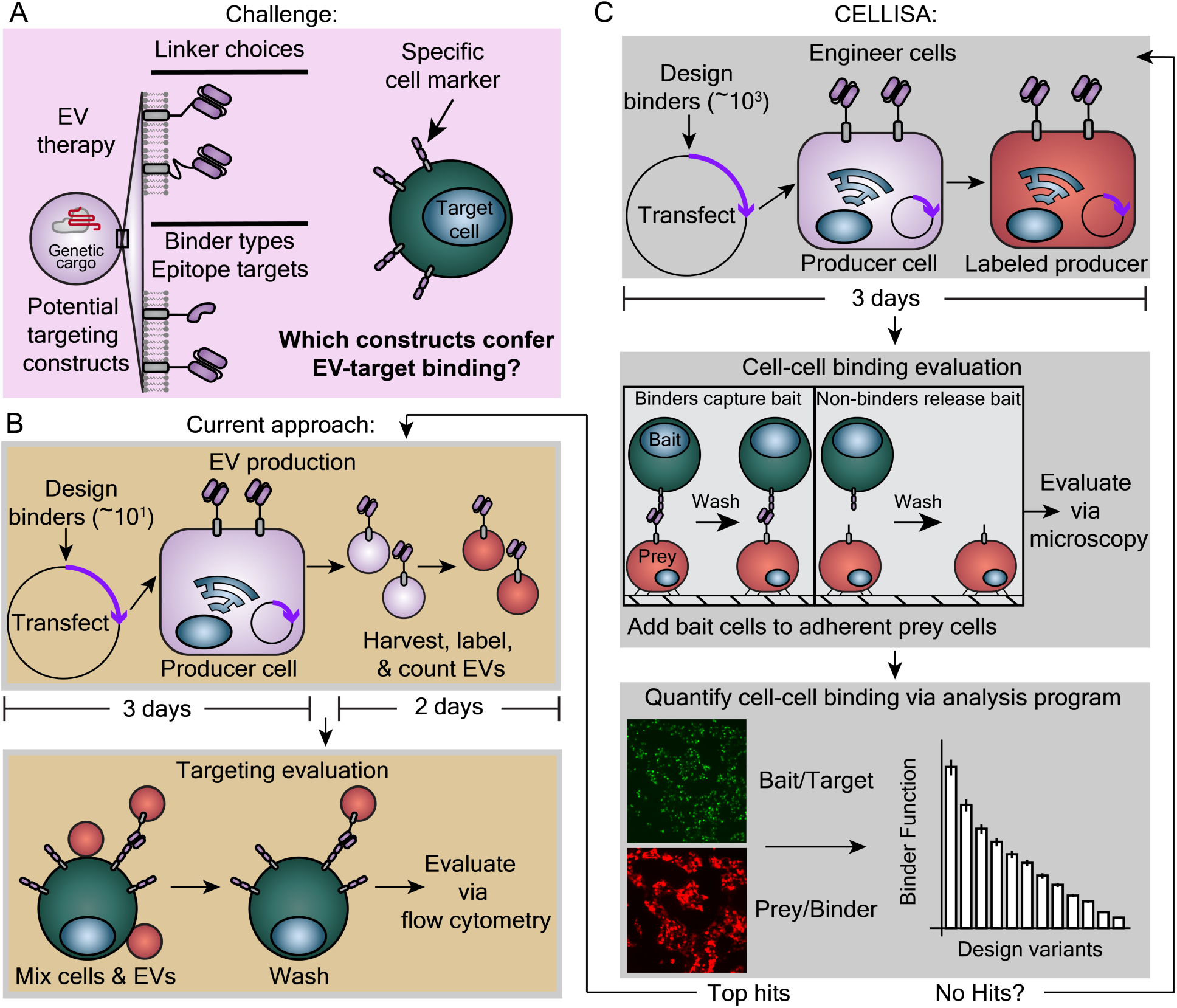
A cell-based binding assay enables higher throughput screening of EV targeting molecules. (**A**) Schematic summarizing a key step in targeting EVs to specific cell types—exploring the design space of engineered targeting proteins. (**B**) Pictorial summary of existing methods for evaluating EV targeting. Producer cells are engineered to express candidate targeting constructs, followed by time consuming and low-throughput processes to isolate EVs and prepare them (top) for EV-cell targeting studies (bottom). (**C**) Schematic overview of the <u>cell</u>-based binding as<u>sa</u>y (CELLISA), which is designed to increase throughput and accelerate screening of candidate targeting proteins by eliminating EV isolation and preparation steps. Shown here is an adherent CELLISA (aCELLISA), where the engineered, binder-displaying cells (prey) are adhered to the plate, and then target cells (bait) are introduced, captured (if the binder is functional), and unbound bait cells are washed away prior to analysis by microscopy.

In this study, we sought to develop a new method for rapidly evaluating targeting constructs, with the specific goal of identifying promising candidates to then be subjected to detailed, standard EV targeting characterization. We investigated whether this first-pass screen might bypass the limiting EV production step by screening targeting constructs in a cell-cell, rather than an EV-cell, binding format. We reasoned that EV producer cell surfaces share many key characteristics with the targeted EVs they generate (e.g., a fluid, lipid bilayer membrane and membrane-tethered targeting constructs), and thus it is possible that cell-cell interactions resemble EV-cell interactions enough to recapitulate binding (**Figure 1B**). This strategy is somewhat analogous to the use of membrane-enclosed virus like particles to display membrane proteins as target epitopes during antibody discovery or vaccination [20]. Supporting our approach, we previously observed that a cell-cell binding assay based on qualitative cellular aggregation could recapitulate EV-virus binding interactions [21]. In this context, we expected that many failure modes in EV targeting would manifest in a cell-based assay; constructs that express poorly, traffic to the cell surface inefficiently, or do not bind to the target epitope should fail in both EV-cell and a cell-cell binding evaluations. We further hypothesized that conserving the relevant membrane context in the down-selection process is needed to predict EV targeting. For example, a membrane-tethered antibody fragment may bind a soluble target in a yeast display study (or vice versa), but that 3D binding geometry may not be physically possible or favorable when both binder and target are displayed on membranes *in trans*. Moreover, binder-target interactions may exhibit avidity when displayed on membranes, which are absent when one or both of the pair is soluble. Even though cell-cell interactions may not predict delivery steps beyond binding (e.g., internalization), separating misses from potential hits (i.e., successful binders) would still be valuable. This study explored these hypotheses.

Here, we report a <u>cell</u> binding as<u>sa</u>y (CELLISA) and analysis pipeline to accelerate the engineering of EV targeting (**Figure 1C)**. The CELLISA methodology may be applied in either adherent (aCELLISA) or suspension (sCELLISA) cell culture formats. In the former approach, drawing inspiration from enzyme-linked immunosorbent assays (ELISAs), fluorescently labelled, “prey” cells are engineered to express a candidate binding construct and adhered to a well plate. Labeled, “bait” cells expressing the receptor being targeted are added such that unbound cells are washed away and successful binders capture the bait cells in the plate. In a sCELLISA, “hits” cause cells to form multicellular aggregates. We built a custom, open-source software package to analyze the resulting fluorescence micrographs and quantify such interactions. We first validated aCELLISA by recapitulating results from a previously published dataset engineering EV targeting using a single chain variable fragment (scFv) binding CD2 (cluster of differentiation 2) on T cells [4]. We then explored how select assay design and protocol choices affected the utility and reproducibility of aCELLISA. We similarly validated and refined sCELLISA for robustness. Finally, we demonstrated the utility of CELLISA through using it in an engineered EV targeting campaign—screening natural killer (NK) and T cell-targeting molecules, identifying candidates, and validating hits in both EV targeting and lentiviral delivery studies. CELLISA vastly improves the throughput and efficiency of EV targeting research and will be of use to researchers developing EV therapeutics.

## Materials and Methods

### General DNA assembly

Generally, new plasmids in this study were ordered from Twist as individual genes cloned into a custom backbone (pPD005 (Addgene plasmid #138749) [22], a pcDNA 3.1 derivative (Thermo V87020). In some cases, plasmids were generated in house by standard restriction enzyme cloning using Twist clonal genes or Twist gene fragments. For in house cloning, restriction enzymes, Phusion DNA polymerase, T4 DNA Ligase, and Antarctic Phosphatase were all from New England Biolabs. Nucleotide sequences for NK-targeting or T cell-targeting constructs were obtained directly from literature or by first reverse translating published amino acid sequences of binders (α-CD16a Nb: D6 (WO 2018/039626 Al) [23]; α-CD16a scFv: LSIV21 [24]; α-NKG2D Nb: ET1F08 (WO2017081190) [25]; α-NKG2D scFv: Tesnatilimab (JNJ-64304500, “MS Light and Heavy Chain”) [26, 27]; α-NKp46 scFv-1: (Seq ID No: 119, US 2020/0131268 A1) [28]; α-NKp46 scFv-09: (Seq ID No: 25-26, WO 2022/216723 A1) [29]; α-CD3 scFv: OKT3 [30]; α-CD3 Nb: T0170060E11 (WO2016180982A1) [31]; α-CD3 DARPin: DARPIN4 (WO2022129428A1); α-CD5 scFv: Seq ID NO 14 (US20200405811A1) [32]; α-CD7 Nb: VHH6 (US10106609B2) [33]; α-CD21 scFv: HB5 [34] and scaffolds (20mer PAS repeats [35]), or using nucleotide sequences from previous work (PDGFR transmembrane domain, helical linker, 40 GS linker [4]). Unless a specific scFv orientation and linker were reported in literature, scFvs were generated in the following orientation: VL-(GGGGS)_5_-VH. Part sequences for NK-targeting and Jurkat-targeting (i.e., binders, transmembrane anchor, linkers) were individually codon optimized for human expression using GeneScript, and then select codons were changed manually to enable synthesis by Twist.

The wild type CD16a allotype (GenBank X52645.1) and the F176V mutant were ordered from Twist as gene fragments and cloned into a second-generation lentiviral transfer plasmid [36]. mKate2 was originally from Shcherbo et al. [37] and miRFP720 was originally from Shcherbakova et al. [38]. The affinity domain of the αCD2 constructs was described previously (although it was displayed in a slightly different construct designed for expression in stable cell lines) [4]; the αCD2 scFv sequence was originally from Connelly et al. [39]. The PDGFR transmembrane domain was originally sourced from pDisplay-pHuji, a gift from Robert Campbell (Addgene plasmid # 61556) [40]. Plasmids used to generate lentivirus for cell line engineering (pMD2.G and psPAX2) were gifts from William Miller at Northwestern University. pREM016 was created by PCR amplification of the mScarlet-I gene from pcDNA3-LumiScarlet, which was a gift from Huiwang Ai (Addgene plasmid # 126623 ; http://n2t.net/addgene:126623 ; RRID:Addgene_126623) [41], and cloned into a pGIPZ lentiviral backbone using NheI and BsrGI. pHIE822, encoding the VSV-G mutant construct, was created by introducing point mutations (K47Q, R354A) in the VSV-G-encoding gene of pMD2.G using site-directed mutagenesis by PCR [6]. The third-generation lentiviral transfer plasmid used as the transfer plasmid in the lentiviral delivery studies, pGGB500, was made with the pCCL backbone, synthesized by Twist Bioscience, and a DsRed-Express2 transgene cloned via restriction enzyme cloning. DsRed-Express2 was obtained by site directed mutagenesis of pDsRed2-N1, which was a gift from David Schaffer (University of California, Berkeley). The third-generation lentiviral packaging plasmids, pRSV-Rev and pMDLg/pRRE, were gifts from David Schaffer (University of California, Berkeley). Plasmids were sequenced before use by either Sanger sequencing, whole plasmid sequencing (via PlasmidSaurus), or by the manufacturer (e.g., Twist for clonal gene synthesis and cloning). Sequence maps for all plasmids are included in **Supplementary Data 1**.

### Plasmid preparation

A glycerol bacterial stock was purchased for each successful Twist clonal gene. Plasmids cloned in house were transformed into TOP10 *E. coli* (Thermo Fisher, chemically competent) and cultured at 37°C. The lentiviral transfer plasmid was grown in NEB Stable competent *E. coli* (New England Biolabs C3040) and grown at 30 °C. Plasmids used in transfections were purified using either a polyethylene glycol precipitation method [22], or more commonly, ZymoPURE Plasmid Miniprep or ZymoPURE II Midiprep kits according to the manufacturer’s instructions. DNA was stored in TE buffer (10 mM Tris, 1 mM EDTA, pH 8.0) at −20°C, and quality was evaluated using a Nanodrop 2000 (Thermo Fisher).

### Cell culture

HEK293FT cells were from Thermo Fisher/Life Technologies (R7007), FreeStyle (HEK293F) cells were from Gibco (R79007), Jurkat T cells were from ATCC (TIB-152), HEK293T Lenti-X cells were from Takara Bio (632180) and NK-92 MI cells [42] were from ATCC (CRL-2408). HEK293F cells were maintained at 37 °C with 8% CO_2_ rotating at 135 rpm on a shaking platform (Infors HT Multitron – I8000A) with orbital diameter 25 mm. All other cells were cultured in a 37°C incubator with 5% CO_2_. HEK293FT cells were cultured in Dulbecco’s Modified Eagle Medium (DMEM, Gibco #31600-091) supplemented with 10% heat inactivated fetal bovine serum (HI-FBS, #16140-071), 4 mM additional L-glutamine (#25030-081), and 1% penicillin/streptomycin (#15140122) from Gibco and 3.7 g/L sodium bicarbonate (Fisher, #S233) and 3.5 g/L glucose (Sigma, #G7021). To support fluorescence imaging, cells were terminally cultured in a phenol-red free DMEM formulation (Sigma #D2902) supplemented with 4 mg/L pyridoxine-HCl (Sigma #P6280), 16 mg/L sodium phosphate (Sigma #S5011), 3.7 g/L sodium bicarbonate, 3.5 g/L glucose, 1% penicillin/streptomycin, 4 mM L-glutamine, and 10% FBS. HEK293F cells were cultured in FreeStyle 293 Expression Medium (Gibco 12338018) supplemented with 1% penicillin–streptomycin. For lentiviral production in HEK293F cells (see ***Lentiviral delivery studies)***, cells were grown in LV-MAX Production Medium (Gibco A3583402) supplemented with 1% penicillin/streptomycin. Lenti-X cells were cultured in phenol-red containing DMEM supplemented with 1 mM additional sodium pyruvate (Gibco, #11360070) Jurkat T cells were cultured in Roswell Park Memorial Institute Medium (RPMI, #31800105) 1640 at pH ≈ 7.2 with 10% FBS, 4 mM additional L-glutamine, 1% penicillin/streptomycin, 3.7 g/L sodium bicarbonate, and 3.5 g/L glucose. NK-92 MI cells were cultured in α minimal essential medium (Gibco #12000022) supplemented with 12.5% non-HI FBS (Gibco, #16000044), 12.5% horse serum (Gibco, #16050122), 1.5 g/L sodium bicarbonate, 1% penicillin-streptomycin, 0.2 mM Myo-inositol (Sigma, #I-7508), 0.1 mM β-mercaptoethanol (Gibco, #21985023), and 0.02 mM folic acid (Sigma, #F-8758).

### aCELLISA wet-lab workflow

To transfect cells for aCELLISA using the calcium phosphate method, tissue culture (TC)-treated, 24-well plates (Corning #3524) were coated with Poly-L-Lysine (PLL) (Sigma P6282 resuspended in sterile, nuclease-free water at 0.1 mg/mL) by incubating PLL solution in the wells for 5 min at room temperature (∼20-22°C), removing excess solution, and allowing the wells to dry for several hours before use. The details of calcium phosphate transfection have been described elsewhere [22]. Briefly, 1.5 – 2 × 10^5^ HEK293FTs were plated in DMEM and allowed to attach for 5-8 h at 37°C in a 5% incubator. Plasmid DNA encoding binding constructs and fluorescent proteins were diluted in nuclease-free, sterile water, mixed with 3 M CaCl_2_ (such that the final CaCl_2_ concentration is 0.3 M). This mixture was added dropwise to 2X HEPES-buffered saline (280 mM NaCl, 0.5 M HEPES, 1.5 mM Na_2_HPO_4_) in equal volumes and mixed via pipetting. After 3 – 4 min, the mixture was pipetted aggressively 8-10x and added dropwise to cells. The plates were rocked briefly by hand and put into an incubator overnight. The next morning, fresh medium was added (typically phenol-red free, serum-containing DMEM). Cells were used in studies the day after the media change. For a 24-well plate, cells were plated in 0.5 mL media and transfected with 500 ng total DNA in 100 µL transfection reagent per well. Where necessary, empty plasmid DNA was added to the mixture to bring the total DNA amount to 500 ng.

aCELLISAs were performed two days after transfection. Bait cells expressing the target of interest (e.g., Jurkat) were suspended in fresh, serum-free versions of their growth medium at 1 × 10^6^ cells/mL, mixed with 5 uL of Vybrant DiO dye (V22886) per mL of cell suspension, and incubated for approximately 2 min at 37°C. Cells were pelleted at 1,500 rpm for 5 min at 37°C (Sorvall Legend X1R centrifuge, TX-400 rotor) and resuspended in fresh medium. Cells were washed once more in serum-free growth medium, and then finally resuspended in serum-free, phenol-red free DMEM at 6 × 10^6^ cells/mL. The medium was aspirated from the transfected, prey HEK293FTs, and 500 µL of the bait cell suspension was added. Samples were incubated on a rocker (VWR mini blot mixer from Avantor #95057-436) for 15 min at room temperature and were manually shaken every 5 min of this 15 min process. Finally, the medium was aspirated, cells were carefully washed once in fresh serum-free medium, and then 500 µL fresh serum-free medium was added for imaging. A BZ-x800 microscope running BZ Series Application (v01.01.00.17, Keyence) with a PlanApo4x (NA 0.2) objective was used. Fluorescence channels were captured using GFP (DiO), Texas Red (mKate2), and Cy5 (DiD) filter cubes from Keyence (OP87763, OP-87765 and OP-87766, respectively) or DsRed (mKate2) from Chroma (490005-UF1). Specific exceptions to these protocols are identified in the relevant figure captions.

### sCELLISA wet-lab workflow

For most sCELLISAs, cells were transfected using polyethylenimine (PEI Max from Polysciences #24765, dissolved at 1 mg/mL in DI H_2_O, pH adjusted to 7.0 with hydrochloric acid, and 0.22 μm-filtered) at a 4:1 by mass PEI Max to DNA ratio. HEK293F cells were diluted in fresh medium to 5 × 10^5^ cells / mL, 2 mL of this suspension was added to each well of a non TC-treated 6-well plate, and all plates were returned to an Infors HT Mulitron – I8000A shaking incubator (37°C, 8% CO2, passive humidity, at 135 rpm on a 25 mm orbit shaking platform). Plasmid DNA encoding the binding constructs and fluorescent proteins (typically 2 μg total per well to be transfected (1200 ng binder and 800 ng fluorescent protein), or 20 μg / mL) were diluted in Freestyle Medium in microfuge tubes to bring the volume to 100 μL per transfection. PEI Max was diluted in Freestyle medium to 80 μg/mL, and 100 μL was added dropwise to the 100 uL of diluted DNA. The entire mixture was mixed 4X via pipetting and incubated for 30 min at room temperature. Afterwards, each transfection mixture was gently pipetted 2X to mix and the 200 uL transfection mixture was added dropwise to HEK293Fs in 6-well plates. The plates were swirled to encourage mixing, and sealed with a breathable, Rayon-based membrane (either Thermo #12-567-05 or BrandTech # 701365). No media change was performed.

Cells were used in sCELLISA two days after the transfection. Bait cells expressing the target of interest (Jurkat or NK-92 MI) were pelleted at 125 x g for 5 min at 4°C and resuspended in fresh, serum-free RPMI-1640 at 1 × 10^6^ cells/mL, mixed with 5 uL of Vybrant DiO dye (V22886) per mL of cell suspension, and incubated for approximately 2 min or 10 min at 37°C for Jurkat cells or NK-92 MI cells, respectively. Cells were pelleted at 1,500 rpm for 5 min at 37°C (Sorvall Legend X1R centrifuge, TX-400 rotor) and resuspended in fresh, serum-free RPMI. Cells were washed once more in serum-free RPMI, and then finally resuspended in Freestyle medium at approximately 1 × 10^6^ cells/mL. In non-TC-treated 24-well plates, 200 uL of freestyle medium was added, followed by 200 uL of transfected HEK293F prey cells, and then 100 uL of bait cells. Plates were incubated at 37°C, 5% CO2 on a rocker (Corning LSE Nutating Mixer #6720) for 15 min before imaging as in an aCELLISA (see ***aCELLISA wet-lab workflow)***. Specific exceptions to these protocols are identified in the relevant figure captions.

### CELLISA analysis workflow

Image analysis was performed using a custom Python script using Visual Studio Code that is posted to GitHub (**Appendix A. Supplementary Data)**. The typical CELLISA analysis workflow is depicted visually in **Figure S1** and described in detail in **Supplementary Note 1**; a software user guide is included in the GitHub repository.

### Generation of CD16a-expressing NK92-MI cells via lentiviral transduction

The day before transfection to produce recombinant lentivirus, 5 × 10^6^ HEK293T Lenti-X cells were plated in 8 mL DMEM supplemented with 1 mM sodium pyruvate in 10 cm TC-treated dishes. Two plates were generated for each lentivirus construct. The following day, cells were transfected via the calcium phosphate method (see *aCELLISA wet-lab workflow*) using 2 mL of transfection reagent and the following plasmid masses per 10 cm plate of cells transfected: 10 µg of transfer plasmid encoding CD16a and a Blasticidin-resistance gene, 8 µg of psPAX2 lentivirus packaging plasmid, and 3 µg of pMD2.G plasmid encoding vesicular stomatitis virus G protein (VSV-G). The following day, the medium was replaced, and the cells were cultured for approximately 32 h. Conditioned media was harvested, centrifuged at 500 g for 2 min at 4°C, and the supernatant was passed through a 0.45 µm filter (VWR 28143-505) and stored on ice until use. NK-92 MI cells were resuspended in fresh medium at approximately 2 × 10^6^ cells / mL, and 1 mL of cell suspension was mixed with 12 mL of lentiviral supernatant and 4 µg / mL final polybrene (EMD Millipore TR-1003-G in 0.9% saline). Cells were spinoculated at 2500 g for 90 min at 37°C, resuspended in the same medium, and incubated overnight. The next day, cells were resuspended in fresh medium and allowed to recover. Five days after transduction, cells were drug selected with Blasticidin at 10 µg / mL for at least one week.

### Generation of cytosolic mScarlet EV producer cells

HEK293FT cells were used to produce recombinant lentivirus for generating a stable cell line expressing mScarlet for EV targeting studies. HEK293FT cells were plated in 10 cm dishes at a density of 5 × 10^6^ cells/dish. Cells were transfected 7 h later with 10 μg of viral transfer vector encoding mScarlet and a puromycin-resistance gene (pREM016), 8 μg psPAX2, and 3 μg pMD2.G via calcium phosphate transfection. Cell culture medium was changed 12-16 h later. Lentivirus was harvested from the conditioned medium 28 h post media change and centrifuged at 500 g for 2 min to clear cells. The supernatant was filtered through a 0.45 μm pore filter (VWR, 76479-020). Lentivirus was stored on ice until use or at −80°C for long term storage. HEK293F cells were resuspended in fresh media, and 0.5-5 mL of lentivirus was added, resulting in a final concentration of 2 × 10^5^ cells/mL in 20 mL total volume. Cells were returned to shaking culture. 24 h later, cells were resuspended in fresh media and gently vortexed to break cell clumps. Drug selection began 2 d post-transduction with 1 μg/mL puromycin (Invitrogen, ant-pr-1). Cells were cultured in antibiotic for at least two weeks with subculturing every one to two days before further characterization.

Cells were prepared for fluorescence-activated cell sorting (FACS) by resuspending in fresh Freestyle media supplemented with 25 mM HEPES and 100 μg/mL gentamycin (Amresco, 0304) at a concentration of 1 × 10^7^ cells/mL. Cells were sorted on a BD FacsAria Ilu using a 552 nm laser (582/15 filter). The top 68-88% brightest mScarlet cells were collected in Freestyle media supplemented with 25 mM HEPES and 100 μg/mL gentamycin. Cells were spun down and resuspended in normal growth medium with 100 μg/mL gentamycin for recovery for approximately 1 week prior to freezing stocks. This cell line (referred to as REM025) was used as the parental cell line for EV targeting experiments in this study.

### EV production and isolation

A total of 20 × 10^6^ REM025 cells in 20mL in a 125 mL PETG Erlenmeyer flask of fresh media were transfected per condition. A total of 20 µg of plasmid encoding a given binder was added to 500 µL OptiMEM (Gibco 31985-070). The diluted DNA was then mixed with 80 µg of PEI in 500 µL of OptiMEM. The mixture was briefly vortexed and incubated at room temperature for 30 min before being added to each flask. The day after transfection, cells were centrifuged at 150 x g for 5 min, the supernatant was removed, and the cells were resuspended in fresh Freestyle Medium. 2 d after transfection, EVs were collected from the conditioned medium by differential centrifugation [43]. Briefly, conditioned medium was cleared of debris by centrifugation at 300 x g for 10 min to remove cells followed by centrifugation at 2,000 x g for 20 min to remove dead cells and apoptotic bodies in a Sorvall Legend X1R Centrifuge (ThermoFisher Scientific 75004261) with a TX-400 4 x 400 mL Swinging Bucket Rotor. The supernatant was centrifuged at 15,000 *g* for 30 min in a Beckman Coulter Avanti J-26XP centrifuge with a J-LITE JLA 16.25 rotor to pellet 15k EVs. The supernatant was collected and 120k EVs were pelleted by ultracentrifugation at 120,416 *g* for 135 min in a Beckman Coulter Optima L-80 XP ultracentrifuge with an SW41 Ti rotor using polypropylene ultracentrifuge tubes (Beckman Coulter 331372). All centrifugation steps were performed at 4 °C. EV pellets were left in ∼100–200 µl of conditioned medium and incubated on ice for at least 30 min after supernatant removal before resuspension via pipetting.

### Nanoparticle tracking analysis for EV quantification

Vesicle concentration in harvested samples was determined with a Malvern Nanosight NS300 and a 652 nm laser (software v3.4). Vesicles were diluted in PBS to between 2 to 10 × 10^8^ particles / mL, injected at setting 30, recorded with camera level 14, and analyzed at a threshold of 7. For each sample, concentration was determined from the average of three, 30 s acquisitions.

### NanoFCM for EV fluorescence measurement

EV fluorescence was determined via NanoFCM Flow Nanoanalyzer vesicle flow cytometer using a 640 nm laser and a 580/40 bandpass filter. For each sample, 1 × 10^9^ EVs were diluted to a total volume of 100 µL using 0.1 µm filtered PBS and loaded into microtubes (Axygen, MCT-060-C). After instrument start up, Quality Control (QC) Beads (250 nm Fluorescent Silica Microspheres, NanoFCM Co., QS3003) were run at a 100× dilution in 0.2 µm filtered Milli-Q water with large particle thresholding for alignment, focusing, and calibration. Following QC checks, a Silica Nanosphere Cocktail (NanoFCM Co., S16M-Exo) was run at a 100× dilution in 0.2 µm filtered Milli-Q water with small particle thresholding for use as a size standard to create a calibration curve of particle size and side scatter intensity. After setup and calibration, samples were run at the NanoFCM software standard sample flow rate, as determined by the sampling pressure of 1.0 kPa, and recorded for 2 min. After data collection, samples were pre-analyzed using the NanoFCM software. Briefly, each sample was thresholded on small particles, and samples were background subtracted using the “Set Blank” software feature, where the blanks were PBS only. Data were analyzed using FlowJo v10 (FlowJo, LLC) as described in detail in the **Supplementary Information** (**Figure S2B**). Mean fluorescence intensity (MFI) of PC5-H of EV samples was exported and used to normalize cellular fluorescence measurements acquired in EV targeting studies.

### Transmission Electron Microscopy

For transmission electron microscopy (TEM), samples were diluted in PBS sterilized with 0.1 μm bottle top filters (Corning 431475) to concentrations on the order of 10^8^ particles per μL as determined by Nanoparticle Tracking Analysis. Grids were plasma treated, and 3 µL of diluted sample were placed on the carbon side of the grid for 30 s. The grids were then briefly washed on 3 droplets of distilled water and placed on a droplet of stain solution (2% ammonium molybdate) for a few seconds. The staining solution was blotted off and the grid was placed on a second droplet of staining solution for 30 s. Finally, the excess staining solution was removed, and the grid was air-dried. Samples were observed with a JEOL 3200FS TEM at an acceleration voltage of 300 kV. A K2 camera was used for recording images.

### Western Blotting of Cell Lysates and EVs

For western blots comparing protein in cell lysates and protein in vesicles, cell lysates were made with RIPA buffer (150 mM NaCl, 0.01% Triton X-100, 0.1% sodium dodecyl sulfate, 0.5% sodium deoxycholate, 50 mM TRIS-HCl pH 8.0, and Milli-Q water to final volume). A Pierce Protease Inhibitor Mini Tablet (Thermo Scientific, A32953) was dissolved in 10 mL of RIPA buffer by rocking at 4°C. Cell lysates were made from approximately 1 × 10^6^ EV producer cells (REM025). Cells were pelleted at 150 x g for 5 min at 4°C in a Sorvall Legend X1R Centrifuge (ThermoFisher Scientific 75004261) with a TX-400 4 x 400 mL Swinging Bucket Rotor, the supernatant was removed, and cells were washed in PBS. Cells were pelleted again at 150 g x 5 min at 4°C and resuspended in 1 mL of RIPA buffer supplemented with protease inhibitor and placed in a cold microcentrifuge tube on ice. The mixture was incubated on ice for 30 min. The mixture was centrifuged at 12,200 x g for 20 min at 4°C. The supernatant was transferred to a new microcentrifuge tube on ice. A Pierce BCA Protein Assay Kit (Thermo Scientific, 23227) was used to the manufacturer’s specifications to determine the total protein concentration of the cell lysate. Briefly, BCA albumin standards were thawed on ice. After thawing, 25 μL of each standard was added to an individual well in a 96-well plate. 25 μL of RIPA diluted 1:1 in nuclease-free water (12.5 μL RIPA, 12.5 μL nuclease-free water) was added to a well to measure background. A 1:1 dilution of the cell lysate and nuclease-free water was added to three wells (12.5 μL of the cell lysate with 12.5 μL of nuclease-free water). BCA reagents A and B were mixed at a 50:1 ratio and then 250 μL was added to each well. The plate was covered and incubated in a 55°C incubator for 10 minutes. A BioTek Synergy H1 Microplate Reader was used to analyze the BCA assay at 562nm absorbance. The concentration of protein in the cell lysate was determined by using the standards to make a background-subtracted calibration curve. A linear equation for this calibration curve was used to calculate the amount of protein in each cell lysate well. The cell lysate protein concentration was determined by averaging the concentration in the three cell lysate wells and multiplying by 2 to account for the 1:1 dilution.

For western blots of EVs, 4.5 × 10^8^ EVs were loaded per lane. 2 µg of total protein for cell lysates was added per lane. An established western blot protocol [22] was followed with the following modifications. In most cases, a reducing Laemmli composition was used to boil samples (60 mM Tris-HCl pH 6.8, 10% glycerol, 2% sodium dodecylsulphate, 100 mM dithiothreitol (DTT) and 0.01% bromophenol blue). In some cases, a nonreducing Laemmli composition (without DTT) was used, as previously reported [21]. After transfer, membranes were blocked while rocking for 1 h at room temperature in 5% milk in Tris-buffered saline with Tween (TBST) (pH: 7.6, 50 mm Tris, 150 mm NaCl, HCl to pH 7.6, 0.1% Tween 20). Primary antibody was diluted in 5% milk in TBST and placed on membranes, rocking for 1 h at room temperature. The membranes were then washed three times with TBST for 5 min each. Secondary antibody diluted in 5% milk in TBST was added and rocked overnight at 4°C. Membranes were then washed three times with TBST for 5 min each. The membrane was incubated with Clarity Western ECL substrate (Bio-Rad, #1705061) and imaged on an Azure c280 running Azure cSeries Acquisition software v1.9.5.0606. Specific antibodies, antibody dilution, heating temperature, heating time and Laemmli composition for each antibody was used as previously reported [21] and noted in **Table S1**.

### Surface staining of NK-92 MI, Jurkat, and HEK293F cells

For surface staining NK-92 MI and Jurkat cells, 2.5 × 10^5^ cells were pipetted into flow tubes with 2 mL fluorescence-activated cell sorting (FACS) buffer (PBS pH 7.4 with 2 mM EDTA and 0.05% bovine serum albumin). Cells were pelleted at 125 x g for 5 min at 4°C and the supernatant was removed. 50 μL of blocking buffer was added and the tubes were flicked to mix. Blocking buffer comprised 40 μL of FACS buffer and 10 μL of 1 mg / mL human IgG (Thermo Fisher #02-7102 diluted 5x in PBS pH 7.4) such that human IgG blocking was performed at 0.2 mg / mL. Cells were blocked for 10 min at 4°C. 50 μL of diluted antibody (**Table S2**) was added to each tube (48 μL FACS buffer and 2 μL antibody). The tubes were gently shaken to mix and stained for approximately 15 min at 4°C. The cells were washed twice with 3 mL of FACS buffer and pelleted at 125 x g for 5 min at 4°C. Cells were resuspended in a drop of FACS buffer and analyzed as detailed in the “Analytical flow cytometry and analysis” section.

For surface staining of HEK293Fs, cells were transfected per the “sCELLISA wet-lab workflow” section. 200 μL of transfected cells were moved to FACS tubes with 1 mL of FACS buffer and pelleted at 150 g x 5 min at 4°C. Cells were blocked with 50 uL of blocking buffer for 10 min at 4°C, and then 50 μL of diluted antibody (**Table S2**) was added to each tube (48 μL FACS buffer and 2 μL antibody). Cells were stained for 30 min at 4°C and then washed successively with 3 mL, 2 mL, and 1 mL FACS buffer with pelleting at 150 g x 5 min at 4°C. Cells were resuspended in a drop of FACS buffer and analyzed as detailed in the “Analytical flow cytometry and analysis” section.

### EV targeting studies

Jurkat T cells were incubated with EVs at an EV-to-cell ratio of 100,000:1 (1 × 10^9^ EVs per 1 × 10^4^ cells). Jurkats were plated in a 48-well plate with 300 µl total volume. Cells were plated at the time of EV addition, and wells were brought to equal volumes with additional RPMI as necessary. Cells were incubated for 2 h at 37 °C, then washed three times in FACS buffer (PBS pH 7.4, 2–5 mM EDTA, 0.1% BSA) supplemented with DAPI (to identify live cells) at a final concentration of 3 μM. A single FACS tube of PBS-treated cells was added to a tube containing FACS buffer without DAPI to identify live cells (DAPI negative). Washing occurred by centrifugation at 150 x *g* for 5 min at 4°C, followed by removing the supernatant and resuspending the cell pellet in FACS buffer supplemented with DAPI (or without DAPI for the DAPI negative tube). After washing, cells were resuspended in one drop of FACS buffer supplemented with DAPI (or without DAPI for the DAPI negative tube) before flow cytometry.

### Lentiviral delivery studies

HEK293F cells were spun at 150 x g for 5 minutes and resuspended in LV-MAX Production Medium (Gibco A3583401). 10mL of cells were seeded in flasks at a concentration of 1 × 10^6^ cells per mL. A total of 10 µg of DNA was mixed in a mass ratio of 0.42 : 0.16 : 0.29 : 0.05 : 0.08 (Transfer : pRSV : pMDLg : VSV-Gmut : Binder). This mixture was brought up to 250 µL with OptiMEM (Gibco 31985-070). The diluted DNA was mixed with 40 µg of PEI in 250 µL of OptiMEM. The mixture was incubated at room temperature for 30 minutes before being added to each flask. The day after transfection, cells were centrifuged at 150 x g for 5 min, supernatant removed, and the cells resuspended in fresh LV-MAX Production Medium. Approximately 28 h after media change, lentivirus was collected from the conditioned medium. The medium was centrifuged at 500 *x* g for 2 min to clear cells, and the supernatant was filtered through a 0.45 µm pore filter (VWR 76479-020). To generate concentrated lentivirus, 3 parts of purified supernatant were incubated with 1 part Lenti-X™ Concentrator (Takara 631231) overnight at 4°C. The next day the mixture was spun at 1,500 x g for 45 min at 4°C. The supernatant was carefully removed. The pellet was resuspended in 200 µL of PBS.

For delivery analysis of concentrated lentivirus, virus was added at volumes of 0 µL, 1 µL, 2 µL, 5 µL, 10 µL, 20 µL or 50 µL to a 24-well plate in duplicate. PBS was added to each well to equivalent volume of the highest lentivirus dose. RPMI with 1 × 10^5^ Jurkat T cells were then added to the wells to bring the final volume of each well to 500 µL. For delivery analysis of unconcentrated lentivirus, unconcentrated lentivirus was added at volumes of 0 µL, 1 µL, 5 µL, 10 µL, 50 µL, 100 µL, or 200 µL to a 24-well plate in triplicate. LV-MAX Production Medium was added to each well to equivalent volume of the highest lentivirus dose. RPMI with 1 × 10^5^ Jurkat T cells were then added to the wells to bring the final volume of each well to 500 µL. 3 d post-transduction, Jurkats were harvested and placed in FACS tubes with 1 mL of FACS buffer (PBS pH 7.4, 2–5 mM EDTA, 0.1% BSA) supplemented with DAPI (to identify live cells) at a final concentration of 3 μM. A single FACs tube of vehicle only (LV-MAX Production Medium or PBS) treated cells was added to a tube containing FACS buffer without DAPI to identify live cells (DAPI negative). Cells were centrifuged (150 × g for 5 min), supernatant was decanted, a drop of fresh FACS buffer, with or without DAPI, was added, and cells were resuspended by briefly vortexing before flow cytometry. Titers were calculated via linear regression-based functional titer calculation. Briefly, the percentage of transduced cells was calculated for each well. Transduction was determined as any cell brighter than the brightest 0.1% of cells with no virus/vehicle only (**Figure S2C**). The Multiplicity of Infection (MOI) was calculated for each well:

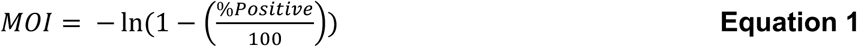

For a single virus type, a linear regression constrained to the origin was fit to the graph of MOI versus volume of virus added to the culture. Each point on the graph was from a single well of cells transduced with that virus. To calculate the functional titer of a virus, the slope of the linear regression was multiplied by the number of cells transduced.

### Analytical flow cytometry and analysis

Flow cytometry was performed on a BD LSR Fortessa Special Order Research Product using the 488 nm laser for SSC and FSC, the 488 nm laser for AF-488 (530/30 bandpass and 505 long pass filter), and the 552 nm laser for mScarlet and dsRedExpress2 (610/20 bandpass filter and 600 long pass filter) or PE (582/15 filter). Typically, 5,000 - 10,000 live cells were collected per sample for analysis. Data were analyzed using FlowJo v10 (FlowJo, LLC). Briefly, cells were identified using a forward scatter area (FSC-A) versus side scatter area (SSC-A) plot and gated for single cells using an FSC-A versus forward scatter height (FSC-H) plot (**Figure S2A, C,S3, S4**). For EV targeting and lentiviral transduction studies, live single cells were identified using a DAPI-A versus SSC-A plot. For EV targeting studies, the mean cellular fluorescence was quantified by the mean of the PE-Texas Red (mScarlet) fluorescence of live single cells and averaged across three biological replicates. The mean cellular fluorescence that was attributable to EVs was determined by subtracting the mean mScarlet fluorescence of HEK293F-containing, PBS-treated negative control samples. This background subtracted mean fluorescence was normalized across samples by dividing by the mean EV fluorescence as determined by NanoFCM analysis. Error was propagated across the analysis. For lentiviral delivery studies, transduction was determined by the percentage of live single cells that were brighter than approximately the 0.1% of the brightest cells in the untransduced (PBS treated) cell samples (**Figure S2C**).

### Statistical analyses

Data were analyzed in Graph Pad Prism 9 or Microsoft Excel. Statistical analyses performed are available in the relevant figure captions; in most cases, a one-way ANOVA was performed with Tukey’s multiple comparisons test.

## Results

### Adherent CELLISA (aCELLISA) recapitulates prior EV targeting data

We first set out to evaluate if a cell-cell binding assay could recapitulate prior data that explored targeting EVs to T cells via the T cell marker CD2 (**Figure 2A**) [4]. Targeted cargo delivery to T cells has therapeutic applications in cancer immunotherapy [44], infectious disease [45], and tissue regeneration [46] and represents a useful test case. In prior work [4], an αCD2 single-chain variable fragment (scFv) was displayed on EVs by fusion to a transmembrane anchor (the transmembrane domain from platelet-derived growth factor receptor, PDGFR) via one of three linkers: a flexible 40 glycine-serine (GS) sequence, a structured helical linker [39], or the hinge motif from immunoglobulin G4 (IgG4) [47]. To recapitulate the delivery evaluated in that study using aCELLISA, we transfected “prey” HEK293FTs to express a red fluorescent protein (mKate2) and the same αCD2 scFv with one of the three linkers. “Bait” Jurkat cells that natively express CD2 were labeled with a green DiO dye, incubated with the scFv-expressing cells, washed, and imaged with a fluorescence microscope (**Figure 2B,C**). These data were analyzed by the CELLISA analysis program, which performed a background subtraction step before counting the number of pixels corresponding to bait and prey cells per image (**Figures 2D, S1, S5A,B**). We built the program to quantify pixel counts rather than cell counts to simplify the analysis step (i.e., to avoid biases and challenges associated with generating masks to identify overlapping cells), and we chose to report the data as a ratio of bait pixel counts to prey pixel counts to normalize for differences in the number of prey cells (i.e., prey pixels) between images. In the aCELLISA study, all three linker designs captured CD2-expressing Jurkats better than a no scFv control, and the helical and 40 GS linker designs significantly outperformed the hinge linker (**Figures 2D, S5B**). We compared these results to targeting data for the two populations of EVs that we evaluated previously: “15k EVs” (pelleted at 15,000 g) and “120k EVs” (the fraction pelleted at 120,000 g) (**Figures 2E, S5C**) [4]. While we expected to detect all hits found in the initial EV-based study, we were surprised that the two data sets were so highly correlated (R^2^ > 0.89 in one experiment and R^2^ > 0.78 in a second experiment) and produced identical rank order of the three linker designs. The aCELLISA study was performed in three days compared to five or more days for an EV targeting study and required substantially less hands-on time to complete (∼3 hours for aCELLISA versus ∼10 hours for an EV targeting study). Therefore, the aCELLISA method successfully recapitulated key conclusions vis-à-vis construct choice for EV targeting using a cell-cell assay in a shorter amount of time, validating the aCELLISA as a useful method to screen binders for EV targeting.

**Figure 2.**
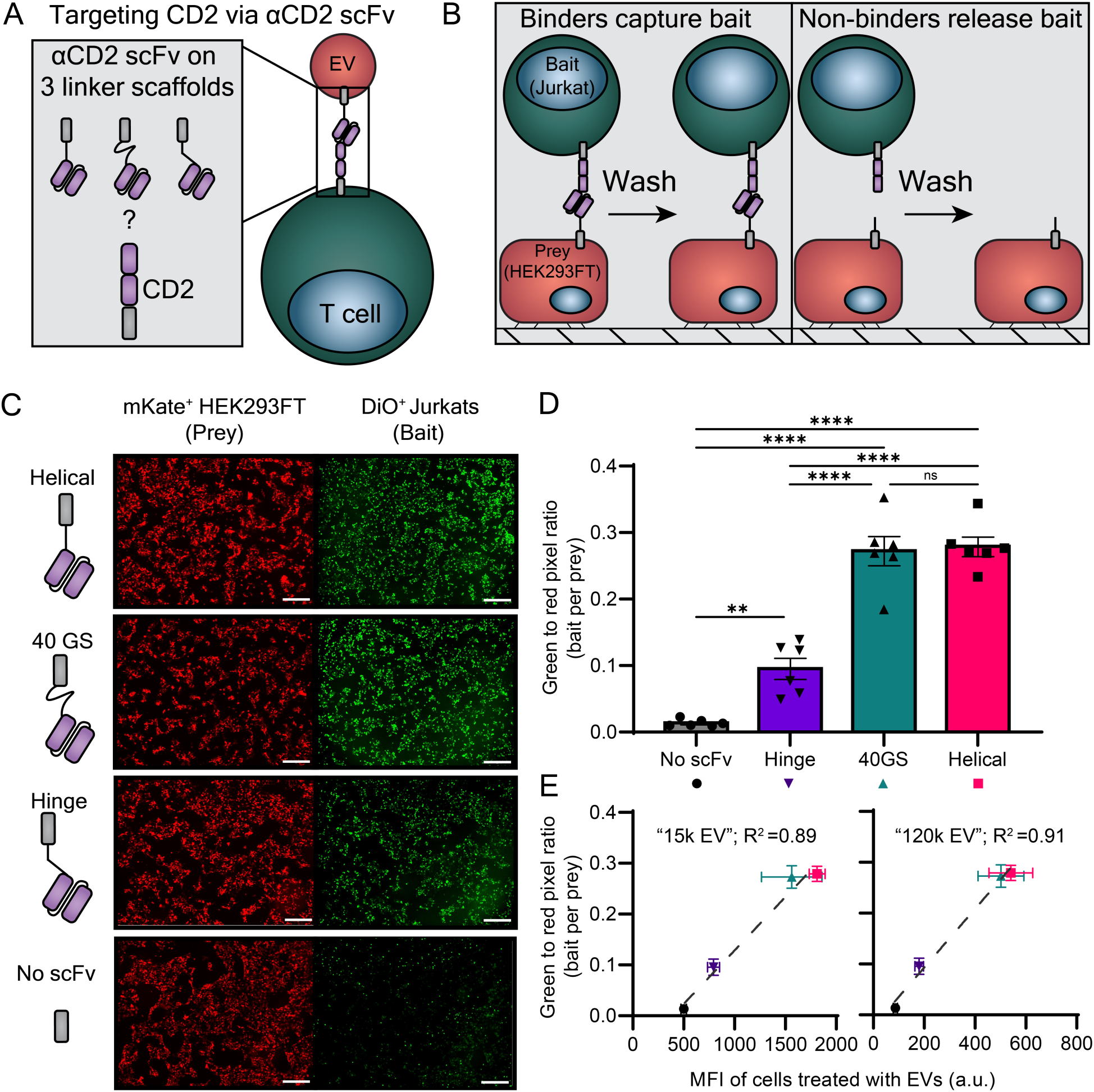
aCELLISA recapitulates rank ordering of EV targeting candidates from an EV-based analysis without the requirement for EV isolation. **(A)** Cartoon describing the goal of this investigation. **(B)** Schematic illustrating the proposed use of the adherent CELLISA method (aCELLISA) to identify scaffolds that enable binding of an αCD2 scFv transiently expressed on the surface of HEK293FTs to CD2 on the surface of Jurkat T cells. **(C)** Sample fluorescence micrographs of prey HEK293FTs expressing one of four binding constructs (left) and bait Jurkat cells expressing CD2 (right). Scale bar represents 500 µm. Images shown here have been brightness and contrast modified for illustrative purposes; the analysis was performed on unmodified images. **(D)** Quantification of data from an aCELLISA experiment, where the green-to-red pixel ratio represents the degree of binding between the two cell types. Each symbol represents a biological replicate (n = 6), the bar represents the mean, and the error bars represent standard error of the mean (SEM). A one-way ANOVA was performed, and the Tukey’s multiple comparisons test are shown (****, p < 0.0001; **, p < 0.01; ns, p > 0.05). Data shown are representative of two independent experiments (**Figure S5**). **(E)** Comparison of aCELLISA data shown in **(D)** with EV targeting data for these constructs for EVs pelleted at 15,000 g (left, “15k EVs”, referred to as microvesicles in the original study) and EVs pelleted at 120,000 g (right, “120k EVs”, referred to as exosomes in the original study) from prior work [4]. Symbols represent the mean of 6 biological replicates for aCELLISA data (y-axis) and 3 biological replicates for EV targeting data (x-axis); error bars represent the SEM. The grey dotted line represents a linear regression drawn through the four points. aCELLISA results highly correlate with EV targeting data (R^2^ > 0.89 for both vesicle populations) and correctly recapitulate the rank order of binders.

### aCELLISA is robust to certain assay design choices

Next, we used the αCD2 system to investigate the robustness of the aCELLISA methodology and quantitation to select experimental design choices. First, we evaluated how the background selection made by the user affects the quantitative metrics that are output by the CELLISA analysis program (**Figure S6**). We found that intentional, systematic, improper background selection of the bait cells as representative of background by the user impaired the ability of aCELLISA to correctly identify the three binding constructs from **Figure 2D**. These data suggest that aCELLISA—as currently implemented—is not robust to poor background selection, and we recommend that users mitigate this limitation by carefully comparing the micrograph being analyzed with the background subtraction preview shown by the analysis program. We found that ensuring the image preview captures individual fluorescent cells without smearing (**Figure S6**, smearing present in “low background”) led to consistent results; this process can sometimes be aided by comparing the fluorescence micrographs to corresponding brightfield images. Next, we explored if aCELLISA is compatible with a protocol wherein the HEK293FT prey cells are dye-labeled instead of labeled via a co-transfected plasmid encoding a fluorescent protein (**Figure S7A**). Interestingly, we found that aCELLISA is only moderately robust to this design choice: correlation between aCELLISA data from DiD-labeled prey cells and EV targeting data was weaker in both independent experiments (R^2^ ≈ 0.21-0.24 and 0.65-0.69, **Figure S7B,C**) than for cells labeled by a co-transfected fluorescent protein-encoding plasmid (R^2^ ≈ 0.78-0.79 and 0.89-0.91, **Figure 2E, S5**). Further, studies using dye-labeled prey cells did not consistently identify all three binding constructs (**Figure S7B,C**), and we therefore recommend engineering prey cells to express fluorescent protein markers (e.g., via transfection) instead of using dye labeling. We speculate that this effect is partially driven by the dissociation between color (i.e., DiD dye) and binding construct expression in the transfected HEK293FT prey cells when they are dye-labeled (i.e., not all dye-labeled cells express a binder). Co-transfection of two plasmids generates correlated expression phenotypes: cells that express one plasmid (e.g., a binder) are likely to express the other (e.g., a fluorescent protein) [48]; in these cases, normalizing the bait pixel number to the prey pixel number helps to account for different transfection efficiencies between wells—this effect is lost when both transfected and non-transfected cells are labeled with dye. We recommend CELLISA designs in which fluorescent labelling (e.g., via dye or fluorescent protein) is employed such that only cells that express the binding protein or target receptor (e.g., αCD2 on HEK293FT cells or CD2 on Jurkat cells) are fluorescent. Finally, we evaluated whether alternate culture plate coating materials such as collagen or fibronectin were compatible with aCELLISA (**Figure S8A**). Indeed, both alternate materials showed similar performance in aCELLISA when compared to our standard poly-L-lysine treatment (**Figure S8B**). These robustness analyses identified CELLISA design considerations that should facilitate its use.

### Suspension CELLISA (sCELLISA) recapitulates prior EV targeting data

Toward expanding the utility of our screening approach, we next explored whether the CELLISA methodology could be adapted to quantitate cell-cell binding between two cell types that are each cultured in suspension. Non-adherent EV producing cell lines such as HEK293F have become leading choices for industrial and academic applications because they are scalable and relevant from an EV-manufacturing perspective [49]. Inspired by qualitative observations made in prior work [21], we pursued a methodology where the two cell types of interest are mixed, and the extent to which the two cell types form multicellular aggregates is quantitated by our analysis program (**Figure 3A**).

**Figure 3.**
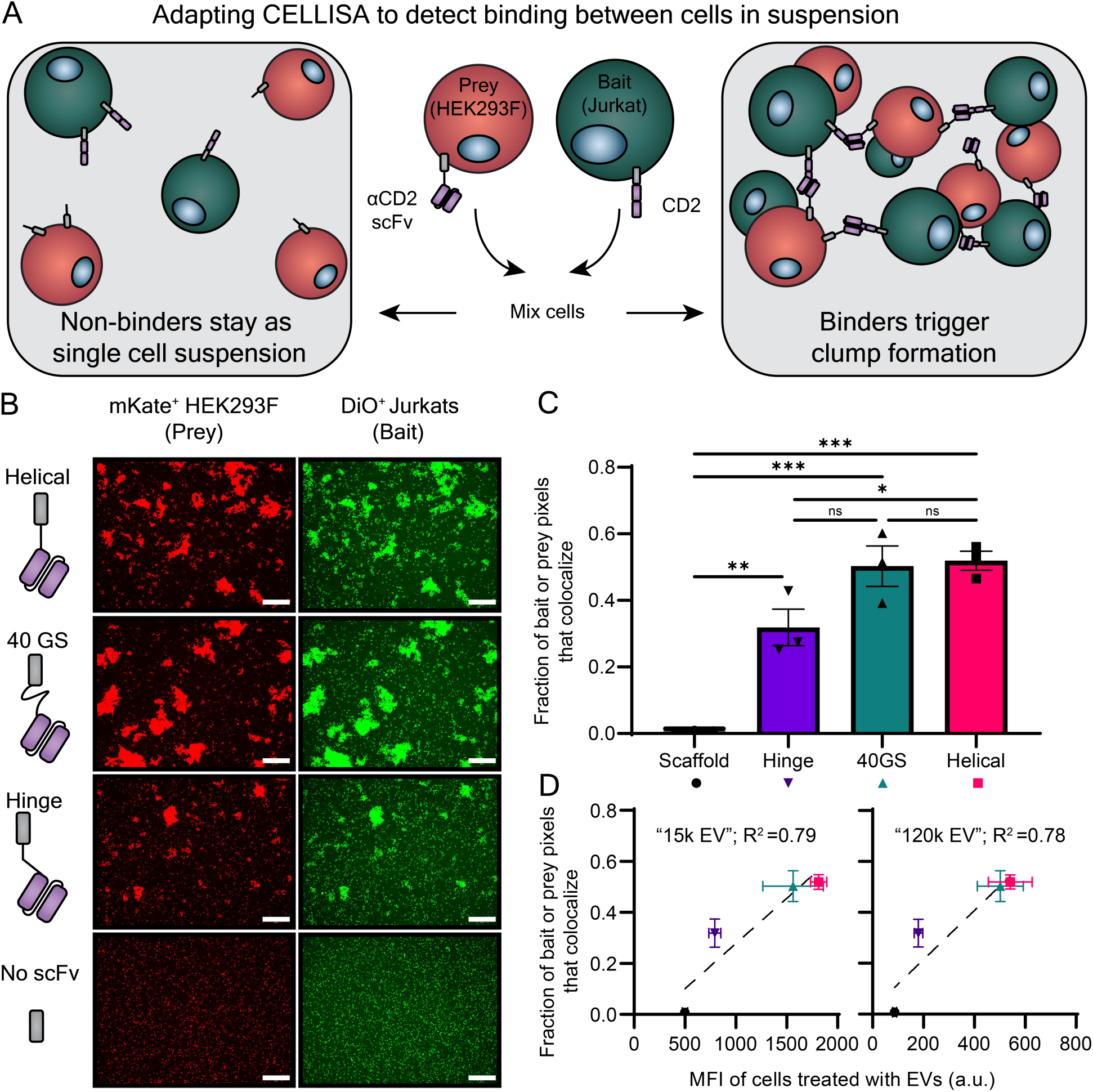
sCELLISA recapitulates rank ordering of EV targeting candidates from EV targeting data without the requirement for EV isolation. **(A)** Schematic illustrating the proposed use of the suspension CELLISA method (sCELLISA) to identify scaffolds that enable binding of an αCD2 scFv transiently expressed on the surface of HEK293Fs to CD2 on the surface of Jurkat T cells. **(B)** Sample fluorescence micrographs of prey HEK293Fs expressing one of four binding constructs (left) and bait Jurkat cells expressing CD2 (right). Scale bar represents 500 µm. Images have been edited for illustrative purposes; the analysis was performed on unmodified images. **(C)** Quantification of data from an sCELLISA experiment, where the fraction of possible pixels that are positive for red and green signals (i.e., that colocalize) indicates interactions between the two cell types. Each symbol represents the average of two fields of view from a biological replicate (n = 3), the bar represents the mean, and the error bars represent SEM. A one-way ANOVA was performed, and the Tukey’s multiple comparisons test are shown (***, p < 0.001; **, p < 0.01; *, p < 0.05; ns, p > 0.05). Data shown are representative of two independent experiments (**Figure S11**). **(D)** Comparison of sCELLISA data shown in **(C)** with EV targeting data for these constructs for EVs pelleted at 15,000 g (left, “15k EVs”, referred to as microvesicles in the original study) and EVs pelleted at 120,000 g (right, “120k EVs”, referred to as exosomes in the original study) from prior work [4]. Symbols represent the mean of 3 biological replicates for sCELLISA data (y-axis) and 3 biological replicates for EV targeting data (x-axis); error bars represent SEM. The grey dotted line represents a linear regression drawn through the four points. sCELLISA results highly correlate with EV targeting data (R^2^ > 0.78 for both vesicle populations) and correctly recapitulate the rank order of binders.

We set out to identify assay design choices that enabled repeatable quantitation of multicellular aggregates using the αCD2/Jurkat system. We transfected HEK293F cells with plasmids encoding a binder and mKate2 using polyethyleneimine (PEI), added these cells to Jurkat cells in a 24-well plate, and incubated them at 37°C with intermittent shaking by hand to encourage mixing, in line with our prior work [21]. This produced detectable, albeit small aggregates (**Figure S9**). We hypothesized that larger aggregates would support more repeatable quantitation and that greater agitation during the incubation could provide more opportunities for cells to interact, bind, and form clumps. Indeed, we found that constant, fast agitation produced large aggregates that enabled higher dynamic range after quantitation than when cells were mixed intermittently by hand (**Figure S9**). Likewise, slow agitation in a larger well format (e.g., 12-well plate instead of a 24-well plate) produced large clumps. We also investigated several metrics for quantitating aggregation formation based on either 1) fluorescent pixels from prey and bait cells colocalizing or 2) calculating the number of bait pixels near each prey pixel using a nearest neighbor algorithm (**Figure S10**). We found that normalizing the number of colocalizing pixels by the number of bait or prey pixels in an image (whichever is smallest) produced a high dynamic range at lower computational cost, and we used this metric going forward (note: both methods are available in our analysis program, **Supplementary Data**). We performed sCELLISAs for the complete binder set from the αCD2/Jurkat system and found that our method correctly determined the rank order of binders repeatedly (**Figure 3B-D**, **Figure S11**) with a good degree of correlation with EV targeting data (R^2^ =0.67-0.79). The method is moderately robust to changes in cell number between experiments, as aggregates were detectable even with 10-fold excess of prey cells (**Figure S12**). Taken together, our results show that the CELLISA methodology, both in adherent and suspension formats, reliably recapitulates EV targeting data without the laborious requirement of harvesting and purifying EVs.

### sCELLISA screening of 36 binders for targeting EVs to NK cells

We next sought to evaluate the utility of the sCELLISA approach by designing a set of binder constructs to target EVs to NK cells—another therapeutically-relevant immune cell type with promising safety and manufacturing advantages [50]. Although engineered NK cells can perform many of the therapeutic functions as engineered T cells (e.g., cancer cell killing), NKs have a better safety profile and may be less expensive to manufacture [51, 52]. We speculated that improving the targeting of biological cargo to NK cells could confer benefits in cell therapy potency, functionality, and manufacturing. We chose to target three well-characterized NK cell markers, CD16a, NKG2D, and NKp46 [53]. CD16a, or FcγRIIIa, recognizes the fragment crystallizable (Fc) region of antibodies bound to molecular targets and is important in directing NK cytotoxicity. NKG2D and NKp46 recognize stress markers on diseased cells and drive NK cells toward an activated, cytotoxic, effector phenotype. For each of the three targets, we selected two antibody-based binders from literature—either scFvs or nanobodies (**Figure S13**). Binders were tethered to the PDGFR transmembrane anchor via one of five linkers: a flexible 40 GS linker, a structured 21 amino acid helical linker, or a semi-flexible, proline-alanine-serine (PAS) linker [35, 54] of 20, 40, or 80 amino acids in length. We evaluated the PAS designs alongside the GS and helical linkers used previously to better understand how linker flexibility and length contribute to targeting capability. These design handles are known to affect liposomal targeting, but they have yet to be systematically explored in the EV field [55]. Thirty-six constructs in total, including matched non-binding controls, were generated to evaluate NK-targeting.

Toward validating our screening methodology, we then performed a sCELLISA to identify NK-targeting constructs using an immortalized NK-92 cell line as bait (**Figure S13**) [56]. We chose to use NK-92s, which are clinically relevant [51, 57], and we selected a clone termed NK-92 MI that grows without requiring a supply of exogenous interleukin-2 (IL-2) [42]. Because NK-92s do not express CD16a natively [58], we generated a CD16a-expressing, NK-92 MI cell line via lentiviral transduction to use in the sCELLISA. When we performed the sCELLISA, we detected no hits from our binder panel (**Figure S13**). We hypothesized that the most likely failure modes could be poor target expression on our NK-92 MI cell line, an incompatibility of the NK-92 MI cell line to produce multicellular aggregates (e.g., perhaps due to different size, mixing propensity, or native adhesion molecule expression relative to the Jurkat studies from **Figure 3**), or poor binder performance (e.g., from either poor surface expression or poor binding capacity in the sCELLISA context). The first explanation is unlikely because surface staining confirmed that our molecular targets were present on the NK cell surface (**Figure S3**). To probe whether the sCELLISA failure was caused by poor binder performance or by some inherent NK-92 MI property, we performed a smaller screen of select NK-binding designs and included the αCD2 helical scFv design from **Figure 3**. NK cells express CD2 (**Figure S3**), and we previously validated this binder design against Jurkat T cells (**Figure 3**), suggesting that its surface expression and binding geometry in HEK293Fs is compatible with sCELLISA. In this study, the αCD2 scFv design generated substantial aggregates between HEK293Fs and NK-92 MIs, while the new NK designs did not, suggesting the original failure mode is not a property of the NK cells but rather a result of poor binder performance (**Figure S14**). None of the NK-targeting designs caused aggregation when HEK293Fs expressing these constructs were mixed with Jurkat T cells, consistent with the lack of target expression on Jurkats (**Figure S3**). Interestingly, most binders in this study showed similar surface expression, which suggests that some other binder property (e.g., binder-target geometry, binding affinity or avidity, etc.) is likely the mechanism behind the failure to discover functional binders (**Figure S4**). Although our NK-targeting library failed to produce a novel hit, this study supports the utility of sCELLISA method: we successfully screened out dozens of poor designs in substantially less time (both hands-on and hands-off time) than traditional methods, reducing the design-build-test-learn cycle. Furthermore, this study demonstrated that the sCELLISA method is compatible with alternate cell types, which supports the idea that this method is generalizable and adaptable to other use cases.

### sCELLISA screening of alternate binders targeting EVs and lentivirus to T cells

Finally, we employed sCELLISA to screen a small library of newly designed T cells binders with various linker types, target molecules, and binder types to improve targeting over our αCD2-helical design (**Figure 4A**). Of the sixteen designs evaluated, we found four hits: an αCD5 scFv and an αCD7 nanobody both tethered to the membrane with a helical or 20-PAS linker (**Figure 4B**). To evaluate whether these CELLISA hits are true positives, we then harvested EVs from HEK293Fs transiently expressing our putative hits, a scaffold control, or the baseline αCD2 scFv with a helical linker. Analysis of our 15k and 120k EVs showed expected protein markers (**Figures S15A,B**), appropriate size distributions (**Figure S15C**), and morphology by TEM (**Figure S15D**) in compliance with the MISEV2023 EV characterization standards [14]. We performed two independent EV harvests and targeting experiments (**Figure 4C)** and found that all sCELLISA hits conferred 15k EVs with significantly more Jurkat T cell targeting compared to the scaffold control (**Figure 4D**). In one of the two independent experiments (**Figure 4D**, top, Experiment #1), 120k EVs expressing the αCD7 nanobody with the helical linker and 120k EVs expressing the αCD7 nanobody with the 20PAS linker demonstrated increased Jurkat targeting compared to scaffold control. These EV targeting results generally correlate with the sCELLISA data (15k EVs: R^2^ = 0.41 - 0.59, 120k EVs: R^2^ = 0.56 across both experiments) (**Figure S16A,B**). Overall, these results demonstrate that our sCELLISA assay can successfully screen for candidate binders that enable increased EV targeting.

**Figure 4.**
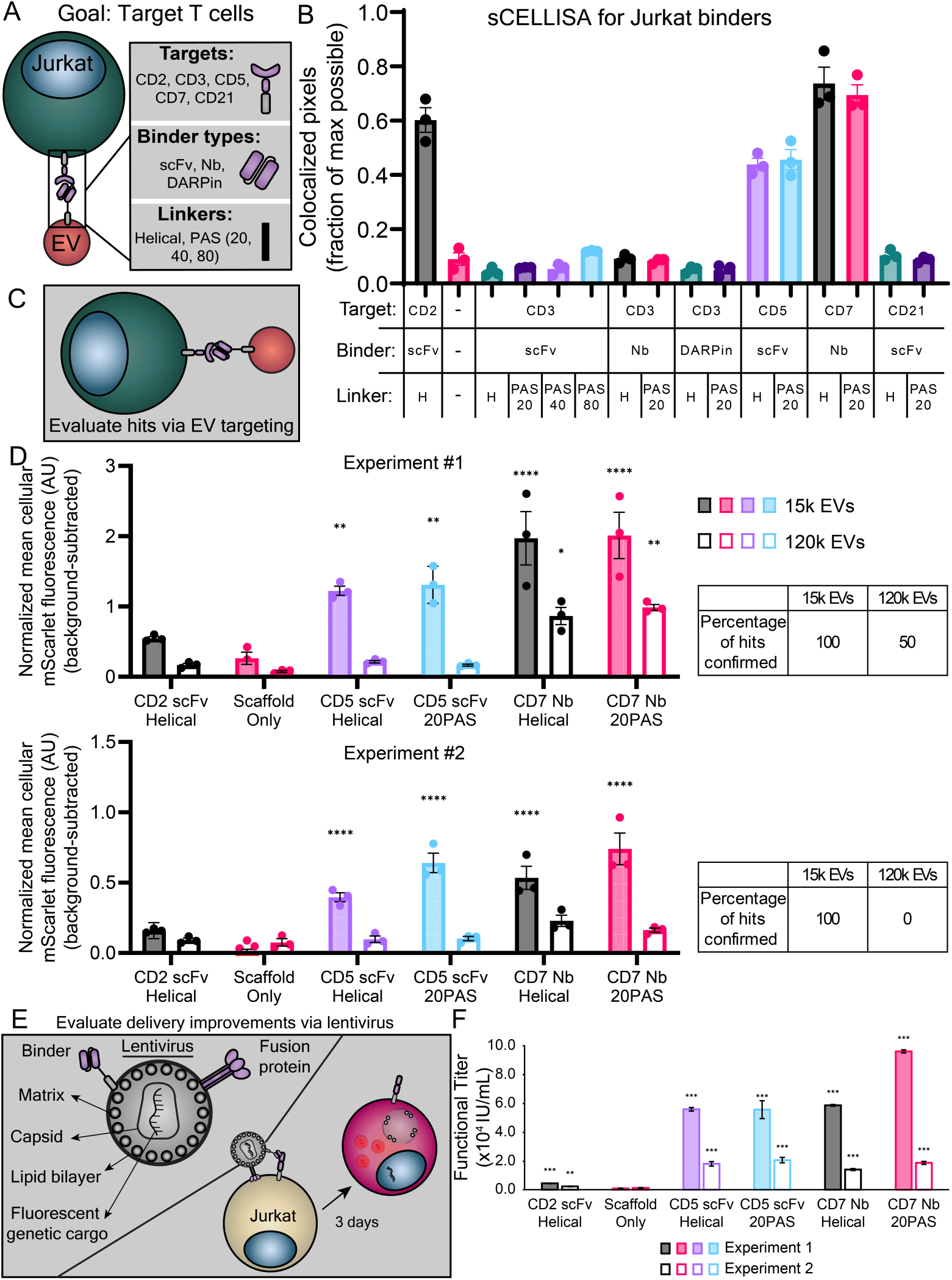
sCELLISA enables identification of targeting peptides for improved EV targeting and lentiviral delivery. **(A)** Cartoon describing the targets, binder types, and linkers used in this investigation. **(B)** sCELLISA data identifying candidate binders to Jurkat T cells. Each symbol represents the average of two fields of view from a biological replicate (n = 3), the bar represents the mean, and the error bars represent standard error of the mean. **(C)** Cartoon depicting the goal of evaluating sCELLISA hits via EV targeting. **(D)** Quantification of 15k (filled bars) and 120k (outlined bars) EVs targeted to Jurkat T cells with binders identified from the sCELLISA assay. The cellular fluorescence of each condition was background fluorescence subtracted (fluorescence attributable to cells only) and normalized by EV fluorescence determined by NanoFCM. Each bar graph represents an independent experiment, and each bar represents the mean of three biological replicates (n = 3). The error bars represent the standard error of the mean. To evaluate whether binders resulted in increased EV binding compared to the scaffold negative control, a two-way ANOVA was performed, and the Dunnett’s multiple comparison test is shown (****, p < 0.0001; ***, p < 0.001; **, p < 0.01; *, p < 0.05; ns, p > 0.05). **(E)** Cartoon depicting evaluation of delivery improvement by candidate binders via lentivirus. **(F)** Quantification of functional titers of unconcentrated lentiviruses expressing mutant VSV-G and the binder indicated on the x-axis. Filled and outlined bars each represent an independent experiment. Each bar represents the mean functional titer as determined via linear regression by calculating the multiplicity of infection using the percent of Jurkat cells expressing dsRedExpress2 (also see Methods: Lentiviral Delivery Studies). Error bars represent the standard error of the slope of the linear regression. To identify if a binder improved delivery compared to the scaffold negative control, a pairwise comparison of each slope from each linear regression to the slope of the scaffold using a one-sample t-test was performed (***, p < 0.001; **, p < 0.01; *, p < 0.05; ns, p > 0.05).

Having established that sCELLISA can identify T cell binders, we next explored whether sCELLISA hits could also confer nanovesicle-mediated delivery of functional cargo. Here, we chose to employ lentiviruses targeting Jurkat T cells as a model system (**Figure 4**). Recombinant lentiviruses are lipid bilayer enclosed nanovesicles, like EVs, that are secreted by cells after being transiently transfected with plasmids encoding: 1) a viral fusion protein, often vesicular stomatitis virus glycoprotein (VSV-G), 2) the lentiviral structural and enzymatic proteins, and 3) the lentiviral genomic RNA including transgenic cargo (here, a dsRedExpress2-encoding reporter gene [59]) that becomes genomically integrated and expressed in target cells [60]. Lentiviruses share many features of EVs (e.g., a fluid lipid bilayer structure), and like EVs, they can be retargeted to specific cell types by loading affinity reagents on their surface [6]. Recombinant, replication-incompetent lentiviruses are commonly used in research [21] and clinical settings where stable expression of transgenic cargo is required [60]; for example, in the engineering of T cell-based chimeric antigen receptor (CAR) cancer therapies [61]. Lentiviruses therefore represent a useful model for understanding how the Jurkat-targeting constructs identified in **Figure 4B** modulate delivery of lipid bilayer-encapsulated genetic cargo. We generated recombinant lentiviruses with a mutant VSV-G that cannot bind its cognate receptor (low density lipoprotein receptor) but retains its membrane-fusion capability [6], along with tested targeting constructs (**Figure 4C**), and we then transduced Jurkat T cells. Excitingly, we found that unconcentrated lentiviruses functionalized with each sCELLISA hit exhibited significantly increased functional titer compared to a control lentivirus (**Figure 4F**). Moreover, compared to our internal reference αCD2 scFv binder, designs identified in the CELLISA increased functional titer up to 22-fold. While overall lentiviral yields varied somewhat across independent experiments, which is typical, the overall trends were consistent. We also confirmed that after concentrating lentiviral preps, each sCELLISA hit also showed significantly increased functional titer compared to scaffold lentivirus (**Figure S17A**). Unconcentrated and concentrated lentivirus functional titers correlated moderately well to our sCELLISA data (R^2^ = 0.19 - 0.37, R^2^ = 0.37-0.47, respectively) (**Figure S17B,C**), suggesting that CELLISA captures some, but perhaps not all, of the determinants of relative functional delivery efficiency. Overall, these data demonstrate that CELLISA can successfully identify binders that confer targeted nanovesicle mediated delivery of functional genetic material.

## Discussion

In this study, we showed that a cell-based method, CELLISA, can be used to efficiently screen surface-binders for their ability to confer nanovesicle targeting and functional delivery to cells of interest. Specifically, we developed a microscopy-based approach and analysis pipeline to evaluate binding of bait (e.g., Jurkat or NK cells) to prey (e.g., transfected HEK293 cells) in both adherent and suspension formats. Our method recapitulated the results of prior EV targeting studies (**Figures 2, 3**) and was robust to several design choices (**Figures S7, S8, S12**, e.g., labeling method, plate treatment, cell count) but sensitive to others (**Figures S6, S9**, e.g., analysis parameters and agitation). We then employed CELLISA to screen NK and T cell binding constructs (on the order of 10^1^–10^2^) and validated the majority of hits in EV targeting and nanovesicle delivery studies (**Figure 4, S16**). Compared to traditional EV targeting studies, CELLISA enabled faster binder evaluation (3 days vs 5+ days) at larger scales (10^1^–10^2^ vs. <10^1^ binders) without specialized laboratory equipment. Altogether, we found that CELLISA is a useful, rapid tool to screen nanovesicle-targeting constructs.

CELLISA’s successful proof of concept will enable future efforts towards increasing throughput. The CELLISA approach is currently well suited to evaluate EV targeting for ∼10^2^ binders with traditional manual, analog pipetting. While this is a substantial improvement over existing techniques, it is small by binder discovery library standards (frequently in the range of 10^7-15^ candidates [62]). An important future direction is increasing the size of CELLISA studies to enable more candidates to be evaluated in parallel. Scaling down the cell culture sizes from 24-well plates to 96- or 384-well plates is an attractive avenue towards reaching a library size of ∼10^3^-10^4^ constructs. This will require technical de-risking to understand how scaling down the well size affects cell mixing, binding interactions, and washing steps (if applicable) (**Figure S9**). In conjunction with scaling down the volume, the use of automated liquid-handling would enable efficient transfection and cell-cell mixing without substantially increasing the labor required to perform these studies, potentially into the 10^4^ to 10^5^ binder range. In certain cases, adherent cell-based CELLISAs might be amenable to plate reader outputs to improve the speed of data collection and analysis, at perhaps a cost in signal or noise. Each of these avenues is a possible strategy for improving the throughput of EV targeting evaluations.

We were motivated to develop CELLISA to better understand how various design choices, such as linker length and composition, affect targeting and nanovesicle delivery, and toward that goal, we evaluated several new linker designs in our T cell and NK cell studies (**Figure 4, Figure S13**). We were surprised to find that our linker designs had negligible impact on CELLISA screening (**Figure 4B, Figure S13),** EV targeting (**Figure 4D**), or lentiviral delivery (**Figure 4F**) outcomes for our three CELLISA hits, which contrasts with prior observations (**Figure 2D**, **Figure 3C**, and prior reports [4]) that linker design can impact performance. To reconcile these observations, we simply propose that linker choice *can* be a useful handle in binder design, depending on the specific binder, target, and cellular context under investigation. We speculate that screening different linker designs is a worthwhile effort to understand the sensitivity of a given binder-target choice to linker composition. Such a screen could be done as part of an initial CELLISA, or as part of a focused, delivery study that evaluates only the most promising binder “hits.”

We were surprised that the αCD2 scFv helical binder did not confer substantial EV targeting for the 15k or 120k EV populations despite success using this design previously [4]. We suspect that several differences in experimental setup could explain this discrepancy. The αCD2 EV targeting data used to benchmark CELLISA (**Figure 2E, 3D**) were acquired from EVs harvested from adherent HEK293FT cells grown in serum-containing medium that stably expressed the binder of interest and a fluorescent protein marker. In contrast, the EV targeting data from the sCELLISA in **Figure 4D** were generated from transient transfection of binder-encoding plasmids into suspension Freestyle 293F cells grown in serum-free medium. These differences may affect the expression, loading, or trafficking of specific binders into EVs (e.g., for our αCD2 scFv) or the specific molecular composition and functional activity of our subpopulations of EVs (i.e., for all binder types). Supporting this hypothesis, we observed some difference in protein markers and relative sizes of EV subpopulations between **Figure S15** and our prior work [4]. Nonetheless, the αCD2 design conferred functional delivery (**Figure 4F**) in our study, as previously observed [4], validating the αCD2 scFv helical design for delivery to Jurkat cells. These subtleties may also be most impactful for binders exhibiting modest performance (e.g., the αCD2 scFv), and they suggest that it could be wise to align CELLISA design with anticipated nanovesicle production format.

This study also highlights an important subtlety for binder discovery campaigns—targeting a nanovesicle to a cell is not necessarily sufficient to enable therapeutic cargo delivery. The therapeutic cargo’s biological mechanism, the degree to which the targeted receptor is internalized after binding, and the method of endosomal escape for a given delivery system, among others, are key considerations that influence delivery efficiency [63, 64]. For example, VSV-G-mediated delivery is dependent on nanovesicle-endosome membrane fusion that is driven by low pH in early to late endosomes [65, 66]. For a lentivirus decorated with VSV-G, a targeting construct that causes receptor internalization to the endosome [24] may drive greater delivery than does a construct that does not trigger receptor internalization [29]. We chose to evaluate CELLISA hits with EV targeting studies and lentiviral delivery assays toward better understanding this phenomenon. In our limited study (**Figure 4**), the rank order from CELLISA was largely, but not entirely, preserved in the 15k EV targeting contexts. Likewise, constructs conferring strong 15k EV targeting also generated higher functional titer lentivirus. We had anticipated that CELLISA might be more predictive of EV targeting than functional delivery *per se*, and indeed we found that correlations with CELLISA hits were stronger for EV targeting studies (R^2^ 0.41-0.59 for 15k EVs, or 0.56 for 120k EVs) than for functional delivery studies (R^2^ of 0.19-0.37) (**Figure S16, S17**). Ultimately, these observations underscore complexities of designing nanovesicle-targeting molecules: 1) targeting does not necessarily confer functional delivery, 2) choices of target, binder, and scaffold (among others features) contribute to performance, and 3) function ultimately needs to be validated in the most relevant cellular and molecular context possible. Although the CELLISA method cannot entirely predict functional delivery, it appears to be a powerful tool to efficiently screen out poor designs such that hits can be evaluated in more relevant contexts.

Looking forward, we envision CELLISA can be further developed to expand its utility. For example, although we focused our efforts on identifying single binder proteins using CELLISA, the method should be amenable to screening multiplex binders or small pools of binders. Binder specificity (i.e., on- vs off-target binding for a given design) could also be evaluated using a mixed population of labeled bait cells and evaluating pixel colocalization for multiple cell types. Finally, CELLISA could be developed to screen for membrane fusion proteins that could enhance nanovesicle-mediated delivery. VSV-G (**Figure 4F**) is a commonly used fusogen [3, 4], but it may not be appropriate for all delivery applications (e.g., due to its wide tropism [67]). With a CELLISA-like method, one could potentially screen viral or synthetic membrane fusion constructs for their ability to fuse bait and prey plasma membranes (with or without a targeting molecule). One could imagine quantifying fusion with aCELLISA or sCELLISA by monitoring colocalization of bait and prey colors as cells fuse to form large, multicolor syncytia or by observing reconstitution of a split fluorescent protein expressed in the bait and prey cells [68, 69]. In sum, we hope that CELLISA will be a useful, open-source tool to help researchers screen targeting constructs for a variety of therapeutic nanovesicles towards the ultimate goal of safer, more effective medicines.

## Acknowledgements

We thank the Kamat and Leonard labs for providing feedback and proofreading the manuscript; Devin Stranford for useful discussions and help performing early pilot studies on cell-cell binding assays; Danielle Tullman-Ercek for help brainstorming the CELLISA acronym during a thesis committee meeting; Tim Vu for collaborating on the CD16a-expressing NK-92 MI cell line; and Roxi Mitrut for use of the cytosolic mScarlet Freestyle 293F cell line for EV production. This work was supported in part by the National Institutes of Health National Institute of Biomedical Imaging and Bioengineering Award 2R01EB026510 (J.N.L.), a gift from Kairos Ventures (J.N.L.), a sponsored research project funded by Syenex, Inc. (J.N.L.), and a McCormick Catalyst grant from Northwestern University (J.N.L.). Y.S.A. was supported by the Jaharis Family Foundation through the Michael Jaharis Undergraduate Research Fellowship. T.F.G., and J.B. were supported by NSF Graduate Research Fellowships (DGE-1842165) N.P.K. was supported by NSF (DMR-2145050 and DMR-2308691). Any opinion, findings, and conclusions or recommendations expressed in this material are those of the authors(s) and do not necessarily reflect the views of the National Science Foundation. T.F.G. and J.B. were supported in part by the Northwestern University Graduate School Cluster in Biotechnology, Systems, and Synthetic Biology, which is affiliated with the Biotechnology Training Program. We thank the Robert H. Lurie Comprehensive Cancer Center of Northwestern University in Chicago, IL, for the use of the Flow Cytometry Core Facility, which provided analytical flow cytometry service. The Lurie Cancer Center is supported in part by an NCI Cancer Center Support Grant #P30 CA060553. Biological analysis was performed in the Analytical bioNanoTechnology Equipment Core Facility of the Simpson Querrey Institute for BioNanotechnology at Northwestern University. ANTEC receives partial support from the Soft and Hybrid Nanotechnology Experimental (ShyNE) Resource (NSF ECCS-2025633) and Feinberg School of Medicine, Northwestern University. This work was supported by the Northwestern University Sanger Sequencing Facility. This work made use of the BioCryo facility of Northwestern University’s NUANCE Center, which has received support from the SHyNE Resource (NSF ECCS-2025633), the IIN and Northwestern’s MRSEC program (NSF DMR-2308691). This work made use of the Center for Synthetic Biology SynBio Foundry facility at Northwestern University, which has received support from the Army Contracting Command (W52P1J-21-9-3023).

## Appendix A. Supplementary data

**Supplementary Information:** This file contains Supplementary Figures, a Supplementary Note, and a Supplementary Table.

**Supplementary Data 1**: This file contains an archive of maps for all plasmids generated in this study.

**Supplementary Data 2**: This file contains source data for all figures in this paper.

**Software**: Software created for this study is published as a public repository on GitHub https://github.com/leonardlab/cellisa.

## Data availability

In addition to the source and supplementary data provided in **Appendix A**, all raw image files used in this study are published in a public repository [70].

## Author contributions

J.B., T.F.G., and J.N.L. conceived the project. Y.S.A., J.B., and T.F.G. performed the experiments. Y.S.A. wrote the analysis program. Y.S.A., J.B., T.F.G., N.P.K., and J.N.L. planned and analyzed experiments. Y.S.A., J.B., T.F.G., N.P.K., and J.N.L. wrote the manuscript. N.P.K. and J.N.L. supervised the work.

