## Supplementary Information for "CELLISA – a cell-cell binding assay for evaluation of nanovesicle targeting proteins"

**Contents:**

Supplementary Figures 1–21

Supplementary Table 1-2

Supplementary Note 1

References cited in this document

### SUPPLEMENTARY FIGURES

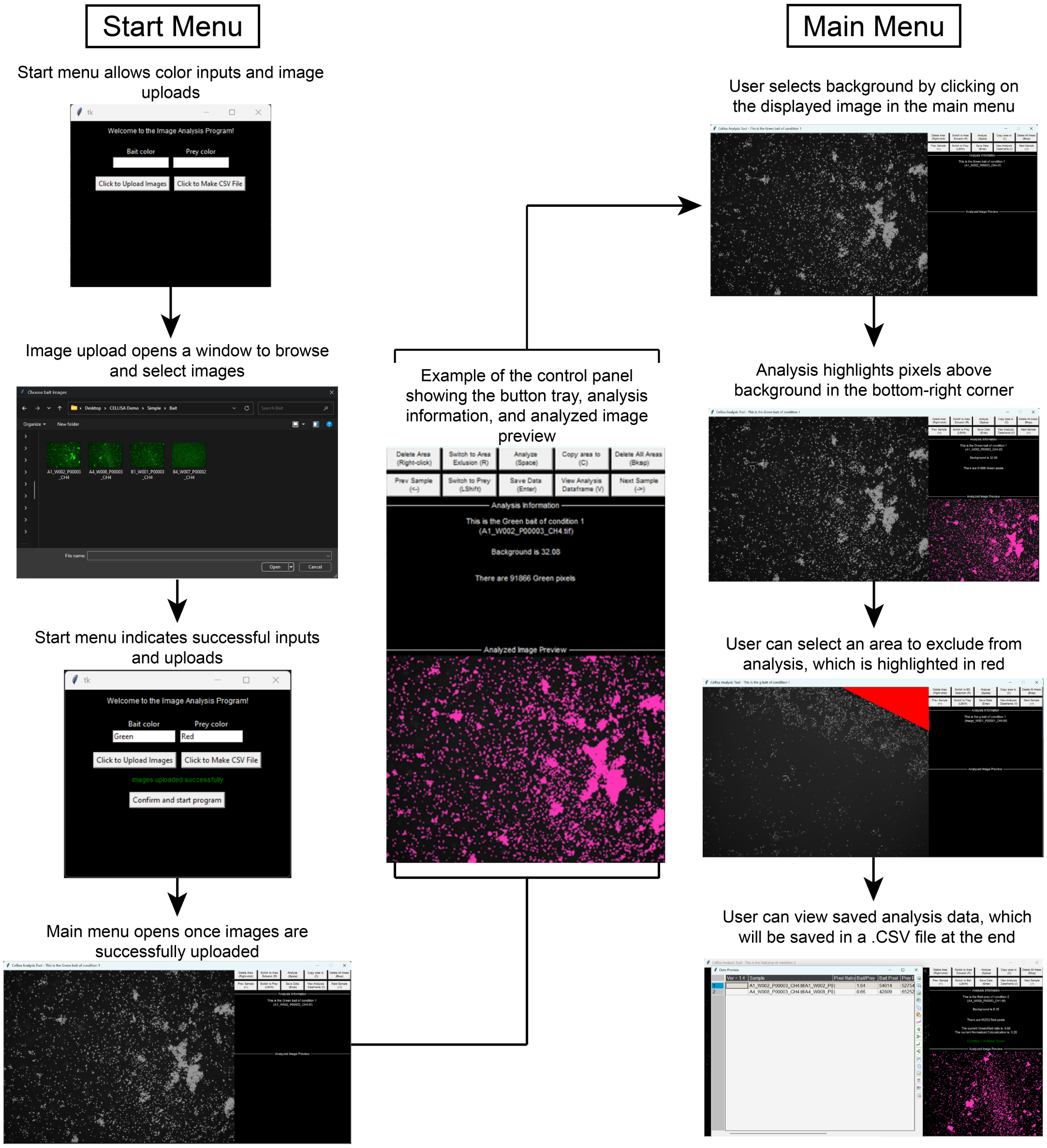


**Figure S1. The key features and data processing steps in the CELLISA analysis script.** Diagram of the analysis program's workflow. This figure illustrates the key steps and features available in the CELLISA analysis program. After opening the micrographs being analyzed (left), the user is presented with a control panel (middle). The program prompts the user to select a polygon encompassing the background for each fluorescence channel (top right), and the pixels that are above the background are displayed in pink in the bottom right of the analysis window to provide the user feedback on the background selection choice. The user can also exclude an area (e.g., the edge of a well) from analysis. After all images are analyzed, the data are appended to an exported CSV file.

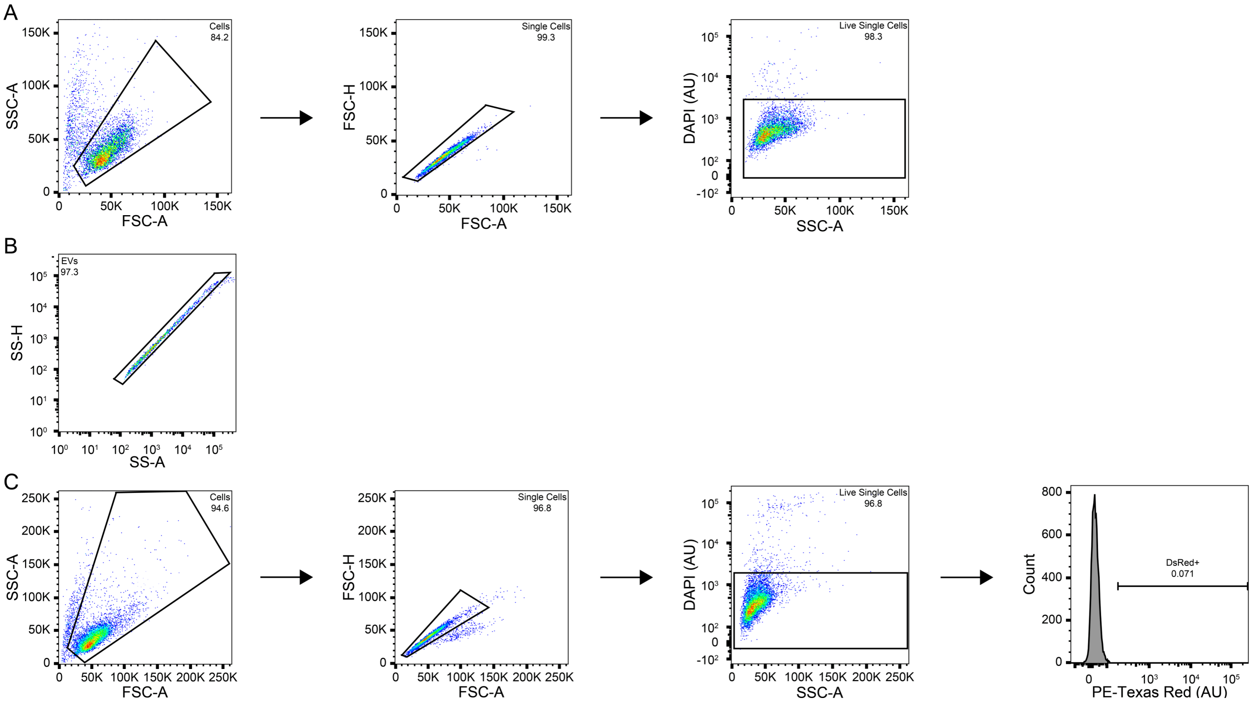


**Figure S2. Flow cytometry and NanoFCM gating. (A)** Conventional flow cytometry gating scheme for EV targeting of Jurkat T cells in **Figure 4D**. The plots show a sample of Jurkats that have not been incubated with EVs (control for background cellular fluorescence). In the gating procedure, cells were identified based on the FSC-A vs SSC-A profile. From this population, single cells were identified based on the FSC-A vs FSC-H profile. Cells were stained with 3 µM DAPI, and live cells were identified on the SSC-A vs DAPI-A profile. **(B)** NanoFCM gating of EVs. The plot shows a scaffold 15k EV sample. In the gating procedure, EVs were identified based on the SS-A vs SS-H profile. **(C)** Conventional flow cytometry gating scheme for lentiviral delivery to Jurkat T cells. The plots show a sample of cells incubated with no vector (negative control for vector transduction). In the gating procedure, cells were identified based on the FSC-A vs SSC-A profile. From this population, single cells were identified based on the FSC-A vs FSC-H profile. Cells were stained with 3 µM DAPI, and live cells were identified on the SSC-A vs DAPI-A profile. Lentiviral gene delivery was defined as all single live cells with a greater PE-Texas Red signal than the top 0.1% of the sample of cells only (no vector addition). These % positive gates were drawn such that they did not encompass more than 0.1% of this non-fluorescent population of cells averaged across three replicates.

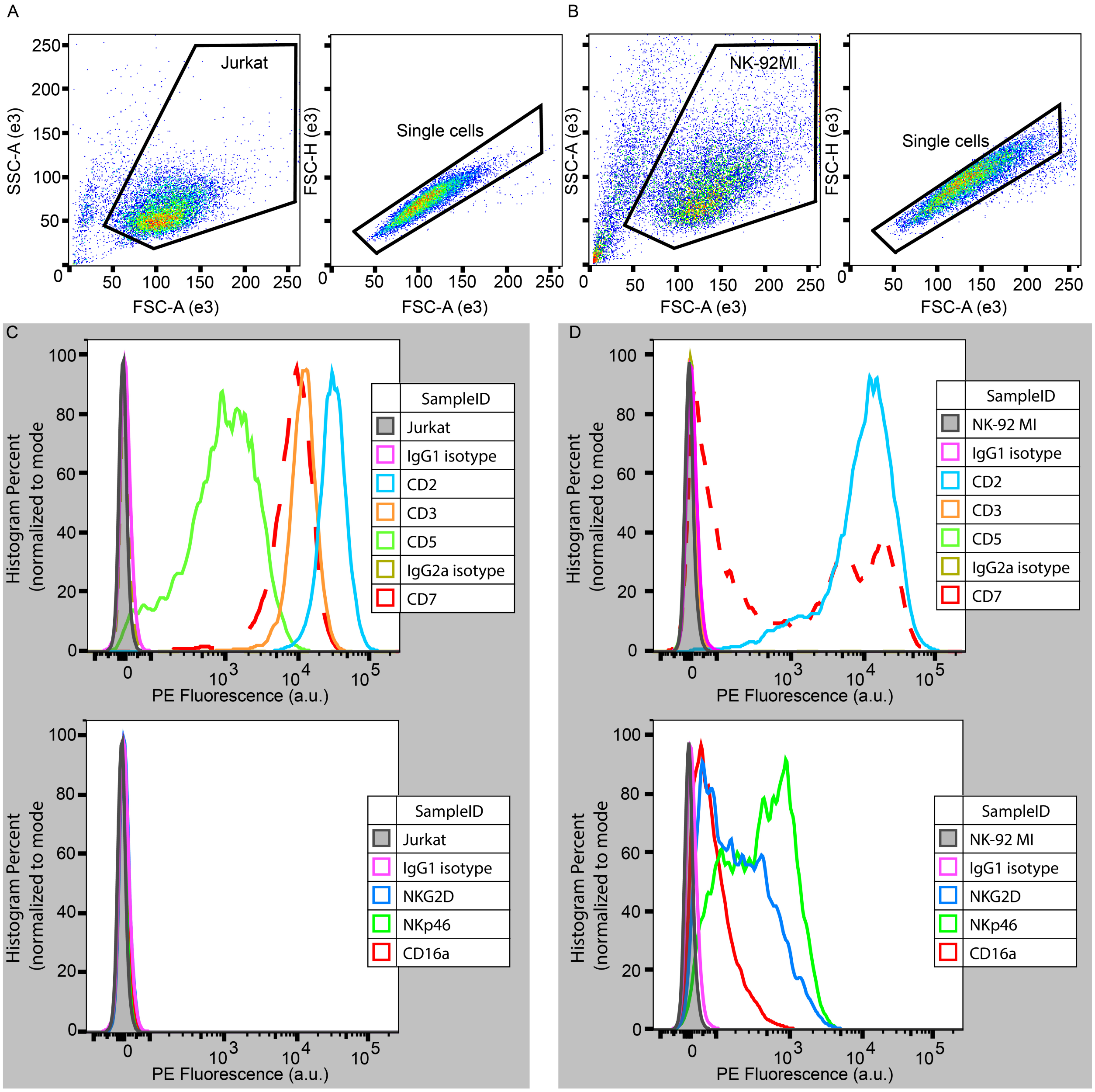


**Figure S3. NK & T cell surface expression of key markers in this study. (A, B)** Flow cytometry dot plots indicating gating strategy. Jurkat and engineered NK-92 MI cells, respectively, were gated first on side scatter area (SSC-A) vs forward scatter area (FSC-A) to identify cells and then forward scatter height (FSC-H) vs FSC-A to identify single cells. **(C, D)** Surface staining data. Surface expression for **(C)** Jurkats and **(D)** engineered NK-92 MI cells for known surface receptors on T cells (top) and NK cells (bottom) via surface staining. Histograms are representative of two independently stained cultures from the same experiment. Here, the Jurkat cells show surface expression of known T cell markers and the NK-92 MI cells show surface expression of known NK markers. The engineered NK-92 MI cells used in this study also surface express CD2 and CD5 which are also markers of T cells.

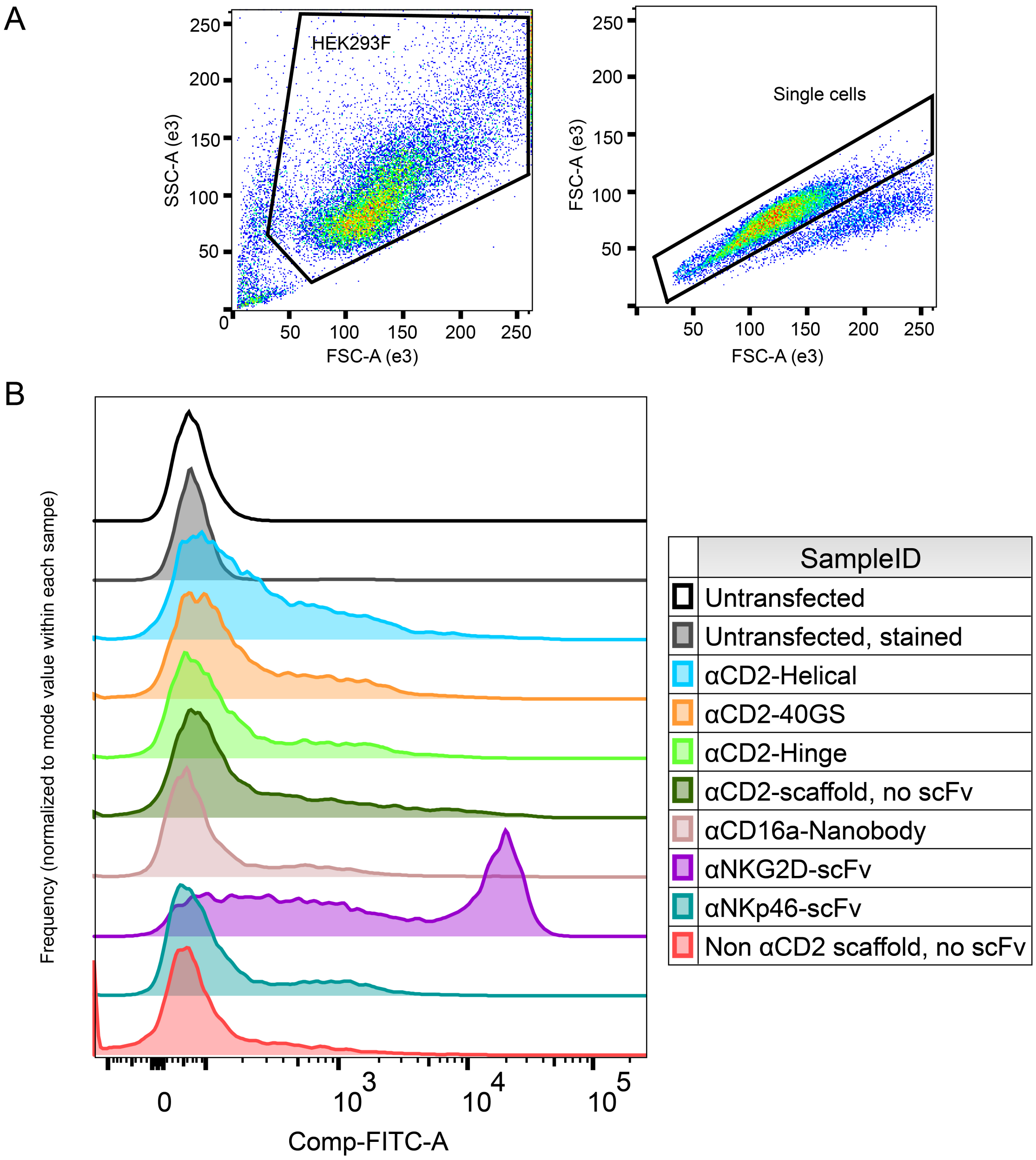


**Figure S4. Transfected HEK293Fs surface express binding proteins. (A)** Gating strategy for HEK293Fs transiently transfected with plasmids encoding putative targeting constructs. **(B)** Surface staining of binding proteins from transfected HEK293Fs. An AF-488 labeled, anti-FLAG antibody was used to stain HEK293Fs to detect surface expression of binders (via the N-terminal 3x FLAG tag). Samples were compensated with single color control samples to account for spectral bleed through between signal from the transfected mKate2-encoding plasmid and the AF-488 antibody stain. Histograms are representative of two biological replicates for all samples except for the untransfected but stained sample (n = 1). The "non-αCD2 scaffold, no scFv" condition is the scaffold used in the αCD16a, αNKG2D, and αNKp46 designs; this condition does not encode an scFv.

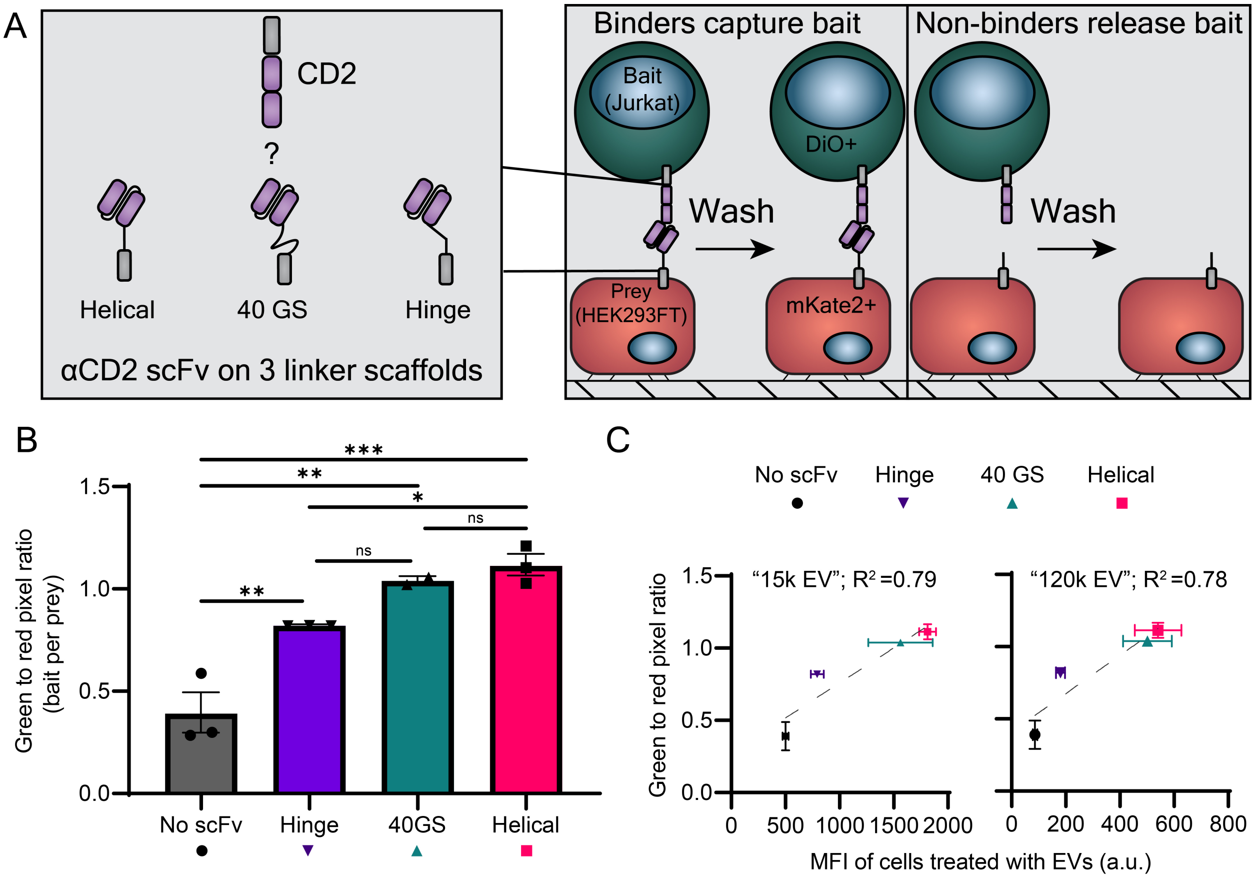


**Figure S5. aCELLISA repeatably recapitulates rank ordering of EV targeting candidates from EV targeting data. (A)** Schematic illustrating the use of the adherent CELLISA method (aCELLISA) to identify scaffolds that enable binding of an αCD2 scFv transiently expressed on the surface of HEK293FTs to CD2 on the surface of Jurkat T cells. In this experiment, Jurkats were labeled with DiO (green) and HEK293FTs were labeled with a co-transfected plasmid encoding mKate2 (red). **(B)** Quantification of a single aCELLISA study, where the green-to-red pixel ratio represents the degree of binding between the two cell types. Each symbol represents the average of three fields of view from a biological replicate (n = 2 or 3), the bar represents the mean, and the error bars represent standard error of the mean (SEM). A one-way ANOVA was performed, and the Tukey's multiple comparisons test are shown (***, $p < 0.001$; **, $p = 0.01$; *, $p < 0.05$; ns, $p > 0.05$). This experiment is a repeat of the study shown in **Figure 2. (C)** Comparison of aCELLISA data shown in **(B)** with EV targeting data from these constructs for EVs pelleted at 15,000g (left, "15k EVs", referred to as microvesicles in the original study) and EVs pelleted at 120,000 g (right, "120k EVs", referred to as exosomes in the original study) from the prior study highlighted in **Figure 2.** Symbols represent the mean of 2 or 3 biological replicates for aCELLISA data (y-axis) and 3 biological replicates for EV targeting data (x-axis); error bars represent the SEM. The grey dotted line represents a linear regression drawn through the four points. As in **Figure 2**, the results indicate a high degree of correlation between aCELLISA results and EV targeting studies ($R^2 > 0.78$).

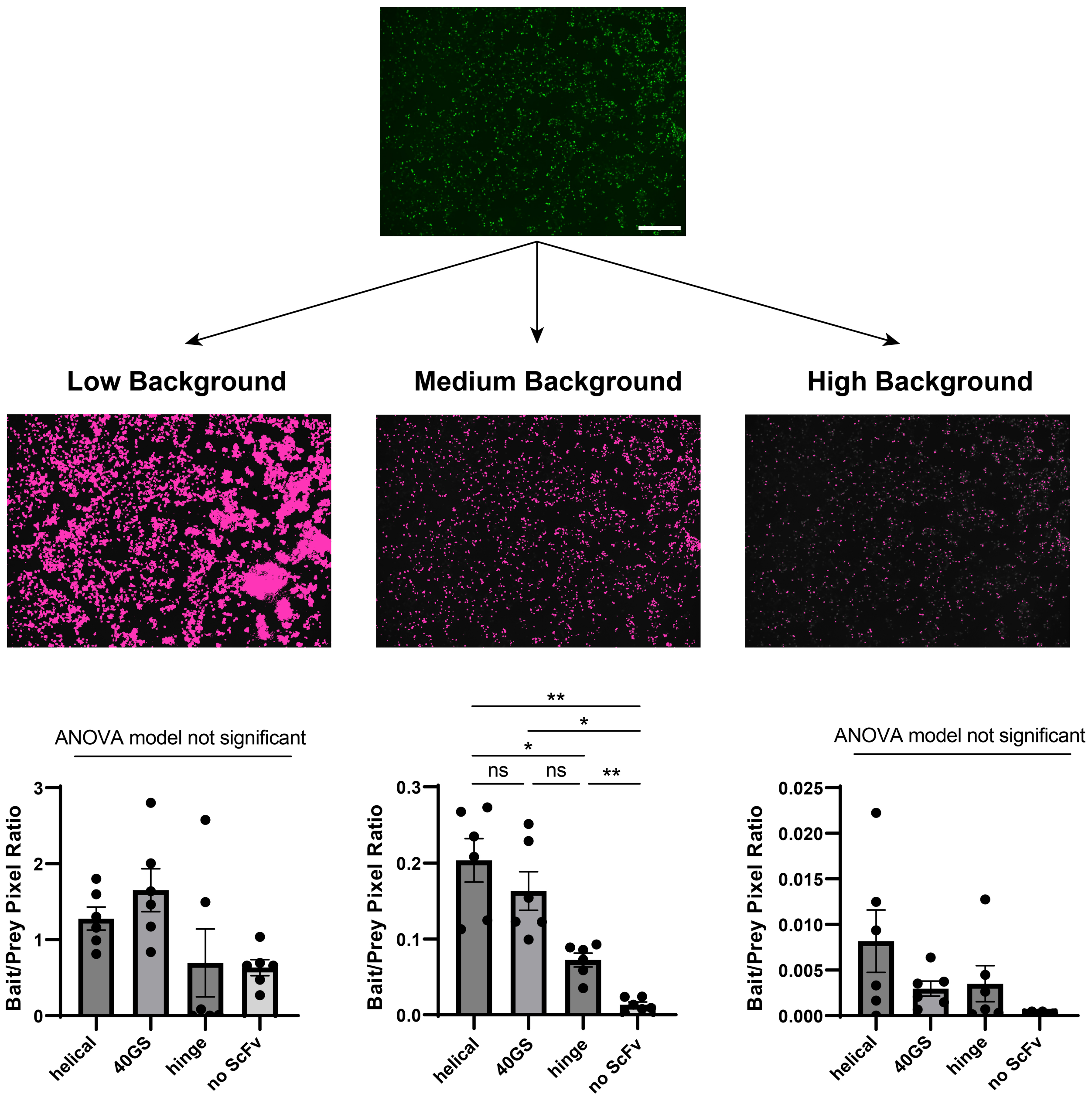


**Figure S6. Improper background selection during image processing impairs the ability of aCELLISA to quantify targeting.** Micrographs illustrating proper background selection ("Medium") and improper background selection ("Low" or "High") of DiO-labeled Jurkat cells during analysis (as described in **Figure S1, Supplementary Note 1**). Note the y-axes are different. An αCD2 aCELLISA dataset (as in **Figure 2**) was analyzed where the HEK293FT prey cells were analyzed with an appropriate "Medium" background, and the Jurkat bait cells were systemically analyzed with too low of a background selection (left), a proper background selection (middle), or too high of a background selection (right) for all conditions. Note that "low background" shows signal (pink pixels) where cells are clearly absent in the original image, while "high background" shows grey pixels (i.e., where fluorescent cells are) that are clearly not pseudocolored pink by the analysis software. The aCELLISA methodology failed to identify any of the three binding constructs with poor background selection; we concluded that proper attention to background selection is critical for aCELLISA. Each symbol represents a biological replicate (n = 6), the bar represents the mean, and the error bars represent standard error of the mean (SEM). A Brown-Forsythe ANOVA test was performed, and Dunnett's T3 multiple comparisons test are shown (**, $p < 0.01$; *, $p < 0.05$; ns, $p > 0.05$). Scale bar represents 500 μm.

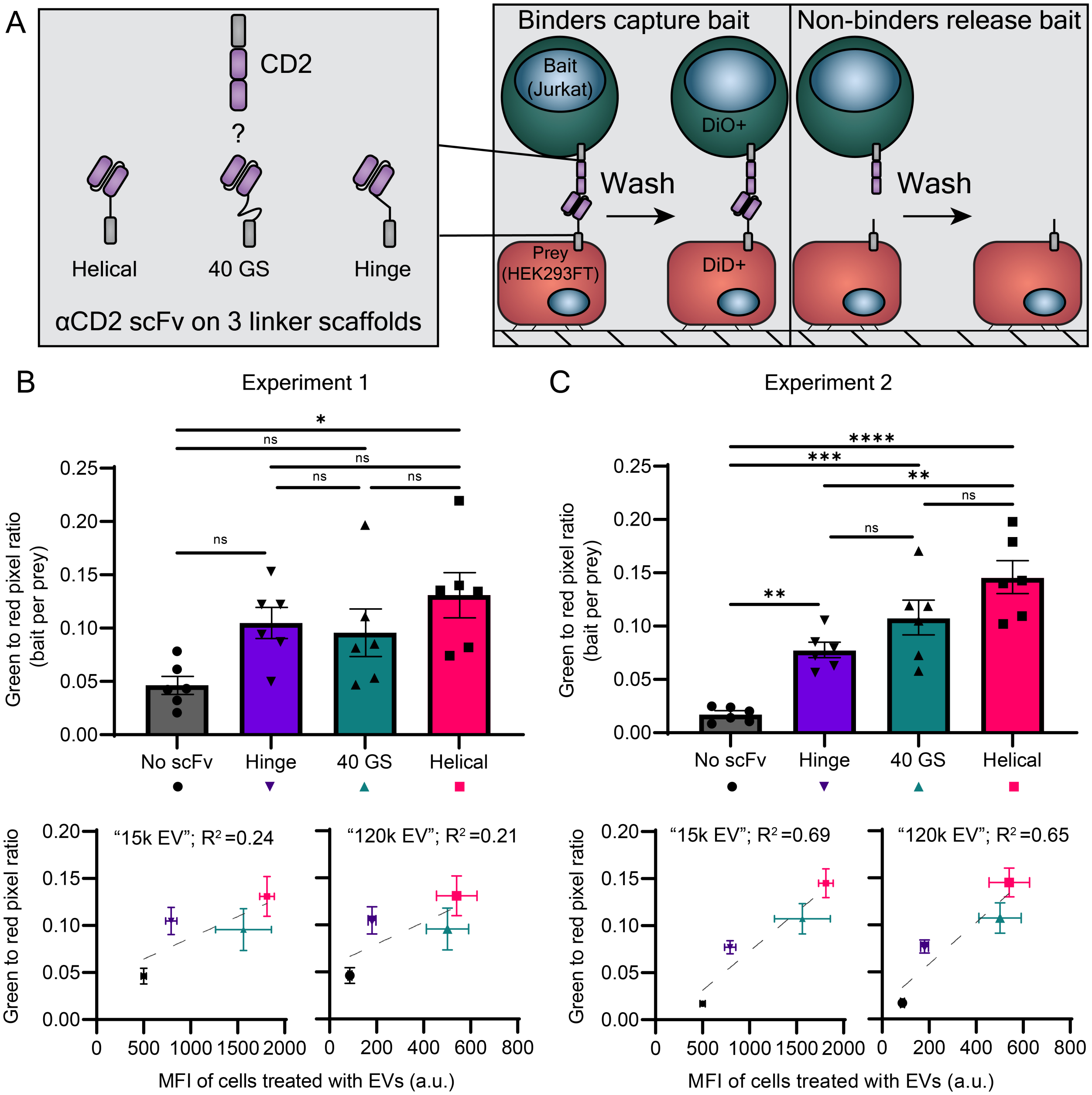


**Figure S7. Studies demonstrate that dye-label transfected HEK293FT prey cells are compatible with aCELLISA quantitation. (A)** Schematic illustrating the use of the adherent CELLISA method (aCELLISA) to identify scaffolds that enable binding of an αCD2 scFv transiently expressed on the surface of HEK293FTs to CD2 on the surface of Jurkat T cells. Jurkats were labeled with DiO (green) and HEK293FTs were labeled with DiD (red). **(B-C top)** Quantification of data from two independent aCELLISA experiments, where the green-to-red pixel ratio represents the degree of binding between the two cell types. Each symbol represents a biological replicate ($n = 6$), the bar represents the mean, and the error bars represent standard error of the mean (SEM). A one-way ANOVA was performed, and the Tukey's multiple comparisons test are shown (****, $p < 0.0001$; ***, $p < 0.001$; **, $p = 0.01$; *, $p < 0.05$; ns, $p > 0.05$). **(B-C bottom)** Comparison of aCELLISA data shown in **top** with EV targeting data from these constructs for EVs pelleted at 15,000g (left, “15k EV”, referred to as microvesicles in the original study [1]) and EVs pelleted at 120,000 g (right, “120k EV”, referred to as exosomes in the original study) from the prior study highlighted in **Figure 2**. Symbols represent the mean of 6 biological replicates for aCELLISA data (y-axis) and 3 biological replicates for EV targeting data (x-axis); error bars represent the SEM. The grey dotted line represents a linear regression drawn through the four points. Dye-labeled HEK293FT aCELLISA results correlate with EV targeting data ($R^2 > 0.2$), but to a lesser extent than when HEK293FTs are labeled with a co-transfected plasmid encoding mKate2 (**Figures 2, S5**). **Methods note**: in the experiment shown in this figure, HEK293FTs were not co-transfected with a plasmid encoding mKate2 for fluorescent labeling. Instead, before plating for transfection, these cells were resuspended in serum-free DMEM at 1 x $10^6$ cells/mL, labeled with 5 µL Vybrant DiD dye (V22887) per mL of cell suspension, incubated for 10 min at 37°C, and then pelleted at 1,500 rpm for 5 min at 37°C. Cells were resuspended in serum-containing DMEM, pelleted, washed in fresh medium, pelleted, and resuspended again in serum-containing DMEM. Cells were then plated for transfection.

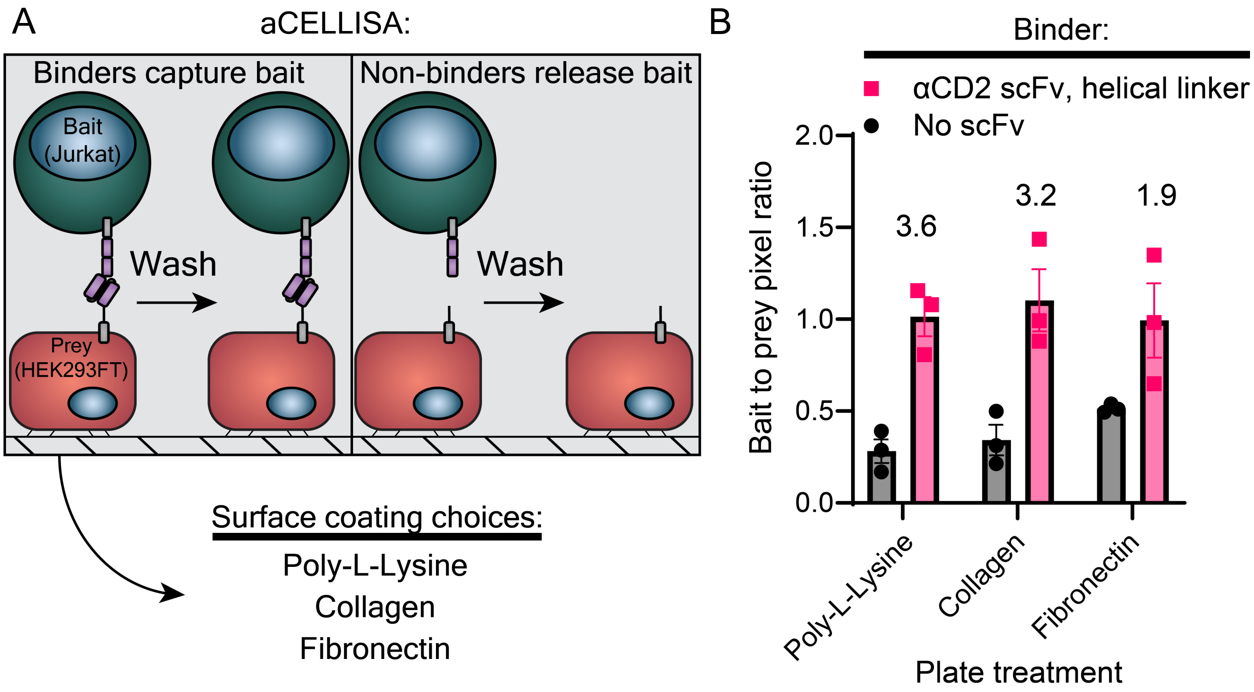


**Figure S8. aCELLISA is compatible with several tissue culture plate treatments. (A)** Schematic illustrating the use of the adherent CELLISA method (aCELLISA). Three surface treatments were evaluated: poly-L-Lysine (used throughout this work), collagen, and fibronectin. In this experiment, HEK293Ts were transfected with plasmids encoding either an αCD2 scFv-helical linker design (**Figure 2**) or a control construct encoding no scFv. Jurkats were labeled with DiO (green) and HEK293FTs were labeled with a cotransfected plasmid encoding mKate2 (red). **(B)** Quantification of data from one aCELLISA experiment, where the green-to-red pixel ratio represents the degree of binding between the two cell types. Each symbol represents a biological replicate (n = 3), the bar represents the mean, the error bars represent standard error of the mean (SEM), and the numbers above the bars represent the average of the αCD2 scFv-helical linker design over the no scFv condition for each plate treatment. All plate treatments enabled detection of binding between the two cell types. **Methods Note:** In this experiment, a modified pre-treatment and aCELLISA method was used. Rat collagen (Gibco A10483-01) was prepared by diluting the stock in 20 mM acetic acid (Sigma 695092 diluted in nuclease free $H_2O$) from 3 mg/mL to 50 µg/mL. Human plasma fibronectin (Gibco 33016015) was prepared by resuspending the sample in 5 mL of nuclease free $H_2O$ for 1 h at 37°C; a working stock was generated by diluting the resuspended fibronectin to 50 µg/mL in PBS pH 7.4. The day before transfection, 200 µL of PLL (prepared as in the Methods section “aCELLISA wet-lab workflow”), collagen, or fibronectin was added to tissue-cultured treated 24-well plates. After 5 min at room temperature, the PLL solution was aspirated. After 60 min at room temperature, the collagen and fibronectin solutions were aspirated. Wells treated with collagen were rinsed with 1 mL PBS pH 7.4 three times to remove residual acid. Immediately after, Lenti-X cells were seeded at 1.5 x $10^5$ cells / mL in 0.5 mL of DMEM supplemented with sodium pyruvate and cultured overnight. Approximately 24 h after plating, cells were transfected via the calcium phosphate method with plasmids encoding mKate2 and either the αCD2-scFv on a helical linker or a scaffold only control. The following day, the media was replaced; the day after the media change, the aCELLISA was performed. Jurkat cells were prepared as per the Methods section “aCELLISA wet-lab workflow” except that after staining with DiO and washing, they were resuspended in phenol red-free, serum-containing DMEM.

During the cell mixing step, the cells were rocked (VWR mini blot mixer from Avantor #95057-436) in a 37°C incubator and shaken by hand once during the incubation. Cells were washed twice in phenol red-free, serum-containing media before imaging.

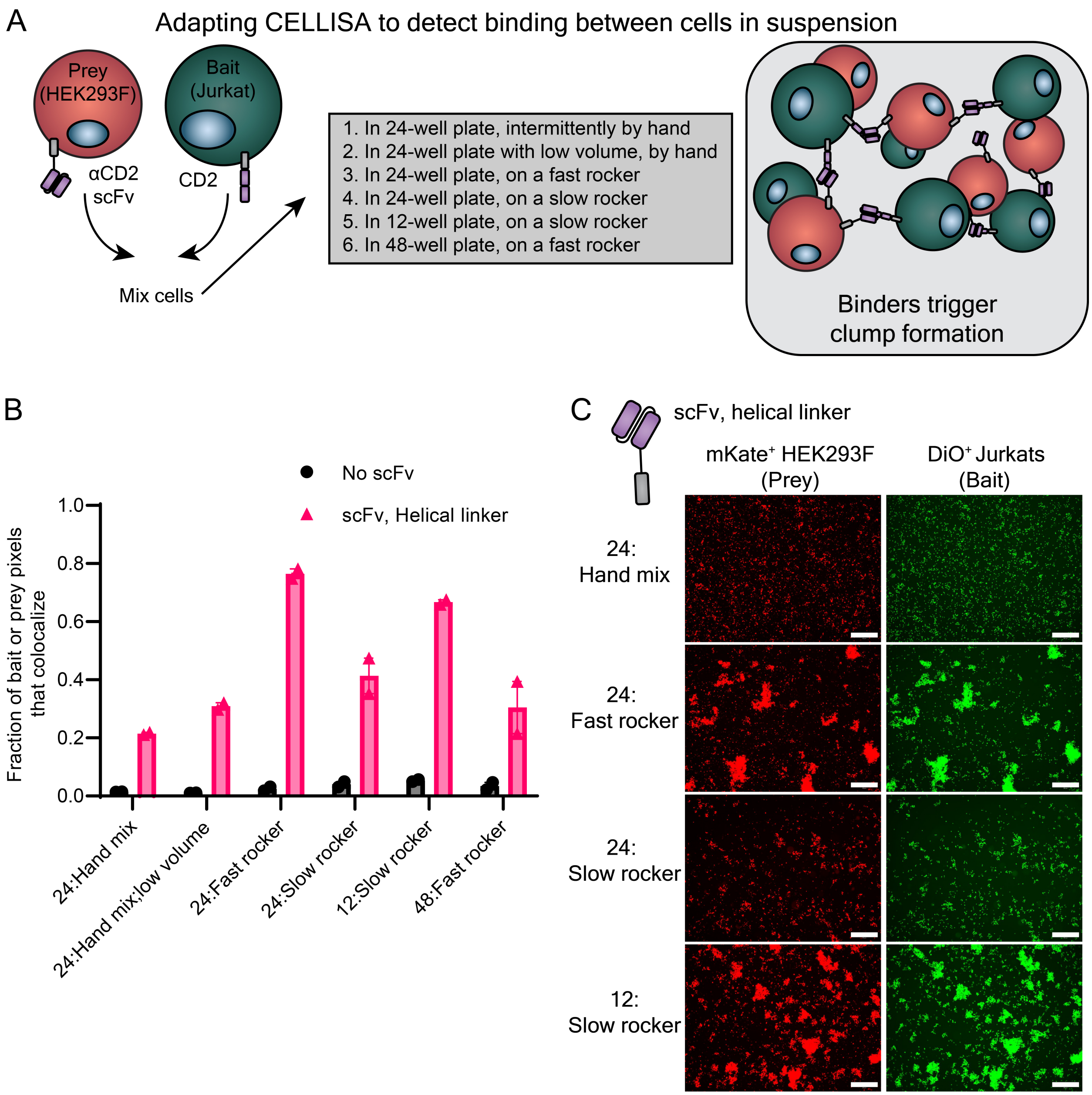


**Figure S9. Mixing conditions govern the extent to which cell-cell aggregates form during sCELLISA. (A)** Diagram of study design. This study evaluated how mixing during the 15-minute sCELLISA incubation step drove cellular aggregation. Jurkat bait and transfected HEK293F prey cells were mixed in 24-, 12-, or 48-well plates by hand or by being placed on a fast or slow laboratory rocker prior to microscopy. We also evaluated if reducing the volume of incubation (from 500 µL to 300 µL) in a 24-well, hand-mix format would change aggregation. HEK293Fs were transfected with plasmids encoding either a scaffold (no scFV) or an αCD2 scFv with a helical linker (i.e., the top performer in **Figure 2**). **(B)** Quantified data from micrographs show that mixing in a 24-well plate with a fast rocker or in a 12-well plate with a slow rocker drove high aggregation in conditions where a plasmid encoding an scFv was transfected. Each symbol is the average of two distinct fields of view within the same well, and the two symbols per bar graph are biological replicates in separate wells. The bars represent the mean and the error bars represent

SEM. This experiment was performed once. **(C)** Sample micrographs from **(B)**. Shown are conditions with the HEK293Fs expressing the scFv-helical linker design to illustrate how different mixing regimes drive different degrees of visual aggregation. The scale bar represents 500 µm. Images have been edited for illustrative purposes; the analysis was performed on unmodified images. **Methods Note**: a modified sCELLISA was performed as part of this method's development. HEK293Fs were seeded in 6-well plates as described in the Methods section "sCELLISA wet-lab workflow", and the cells were transfected with Lipofectamine LTX with PLUS reagent (Thermo #15338100). DNA (2 µg total per well to be transfected) was diluted in Freestyle medium to bring the total volume to 98 µL per transfection, and 2 µL PLUS reagent was added and mixed via pipetting. LTX was diluted in Freestyle medium (7.6 µL LTX into 92.4 µL per well to be transfected), mixed well, and added dropwise to the DNA mixture. The transfection reaction was mixed 4X, incubated for 5 min at room temperature, mixed 4X, and the 200 µL mixture was added dropwise to cells. The plates were swirled, sealed, and returned to the shaking incubator as described earlier. Two days later, the sCELLISA was performed. Bait Jurkat cells were labeled with DiO as described in the Methods section "sCELLISA wet-lab workflow" and mixed with transfected HEK293Fs at 37°C for 15 min before imaging in non-TC treated plates. For sCELLISAs performed in 24-well plates, 200 µL Freestyle media, 200 µL transfected HEK293Fs, and 100 µL Jurkat cells were added. Conditions in 12-well plates had all volumes doubled, and conditions in 48-well plates had all volumes halved. For the sCELLISA performed in a 24-well plate with low volume, the 200 µL medium was omitted (i.e., 300 µL total volume). Hand mixed conditions were mixed by gentle swirling for ~15 s every 5 min. Conditions mixed on a "slow" rocker (~18 rpm with 5° of pitch) were placed on a VWR mini blot mixer (Avantor #95057-436) during the incubation. Conditions mixed on a "fast" rocker (~24 rpm with 20° of pitch) were placed on a Corning LSE Nutating Mixer (Corning #6720) during the incubation.

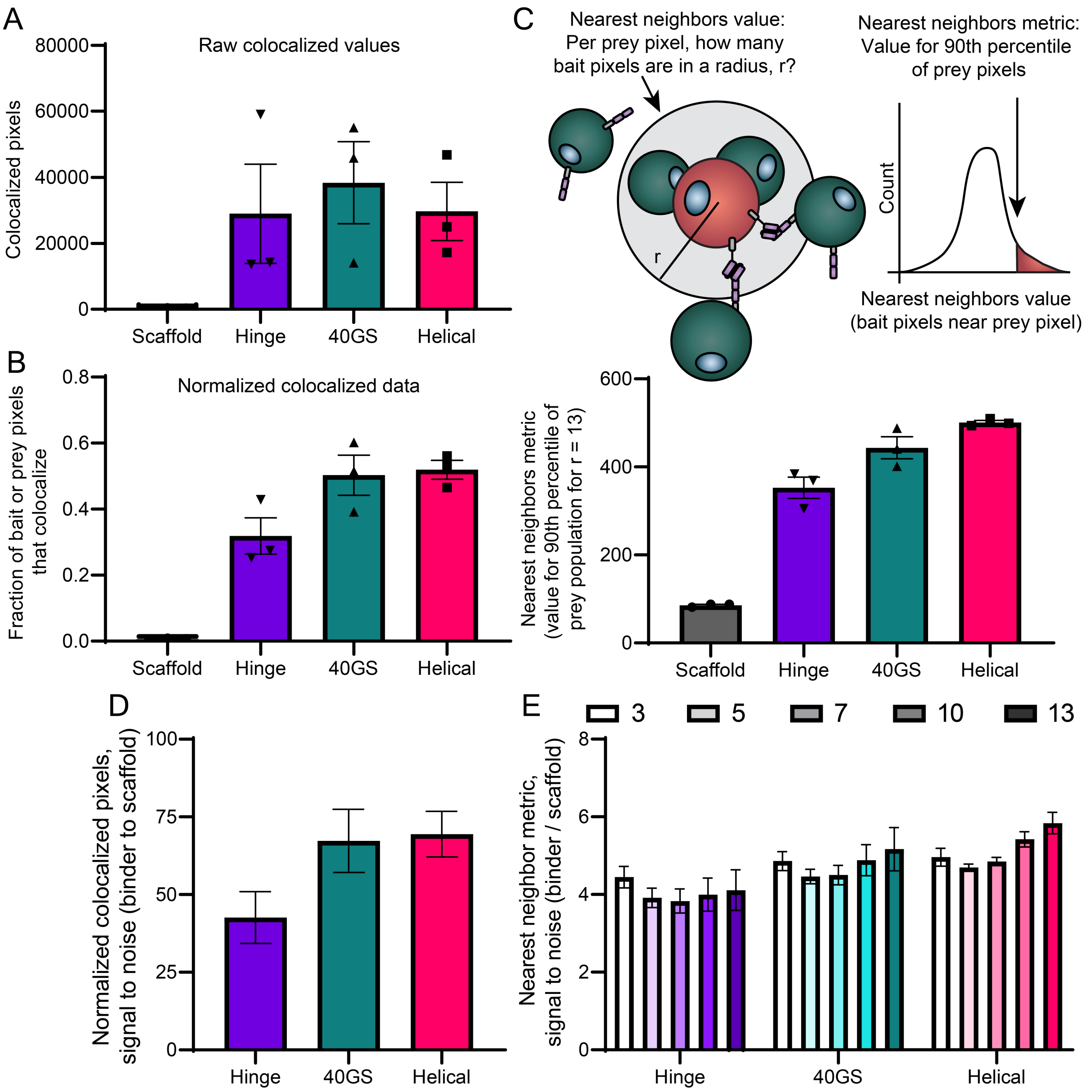


**Figure S10. Normalized pixel colocalization is a useful sCELLISA method that provides high signal-to-noise. (A)** Raw colocalized values from the sCELLISA study presented in **Figure 3C**. (**B)** *Normalized colocalized* values (see **Supplementary Note 1**) presented in **Figure 3C**. **(C)** Definition of a *Nearest neighbors* metric and calculation of this metric for data presented in **Figure 3C.** We defined a *Nearest neighbors* metric that is explained visually in the summary cartoon (top) and elaborated in **Supplementary Note 1**. This population-level metric captures the number of bait pixels that are "near" each prey pixel, is a function of the value of r selected to defined a threshold of proximity, and focuses on the most-clustered regions of an image (see **Supplementary Note 1**). **(D,E)** Signal-to-noise ratios (defined as the metric values for the validated binders divided by the same metric value calculated for the scaffold only condition) for **(D)** normalized colocalization and **(E)** nearest neighbors values calculated for multiple choices of r used to define "near" (in units of pixels; inset legend). Multiple values of r were evaluated to determine whether the signal-to-noise varies by changing the threshold for defining proximity (i.e., in the nearest neighbors definition). The effect of changing this threshold value of r was quite modest under the conditions evaluated. Signal to noise was higher in our system when using normalized colocalization compared to the nearest neighbors method. Thus, normalized colocalization was selected for subsequent analysis. Symbols represent the average of two images per well, bar graphs represent the mean, and error bars represent SEM. Error is propagated for panels **D** and **E**.

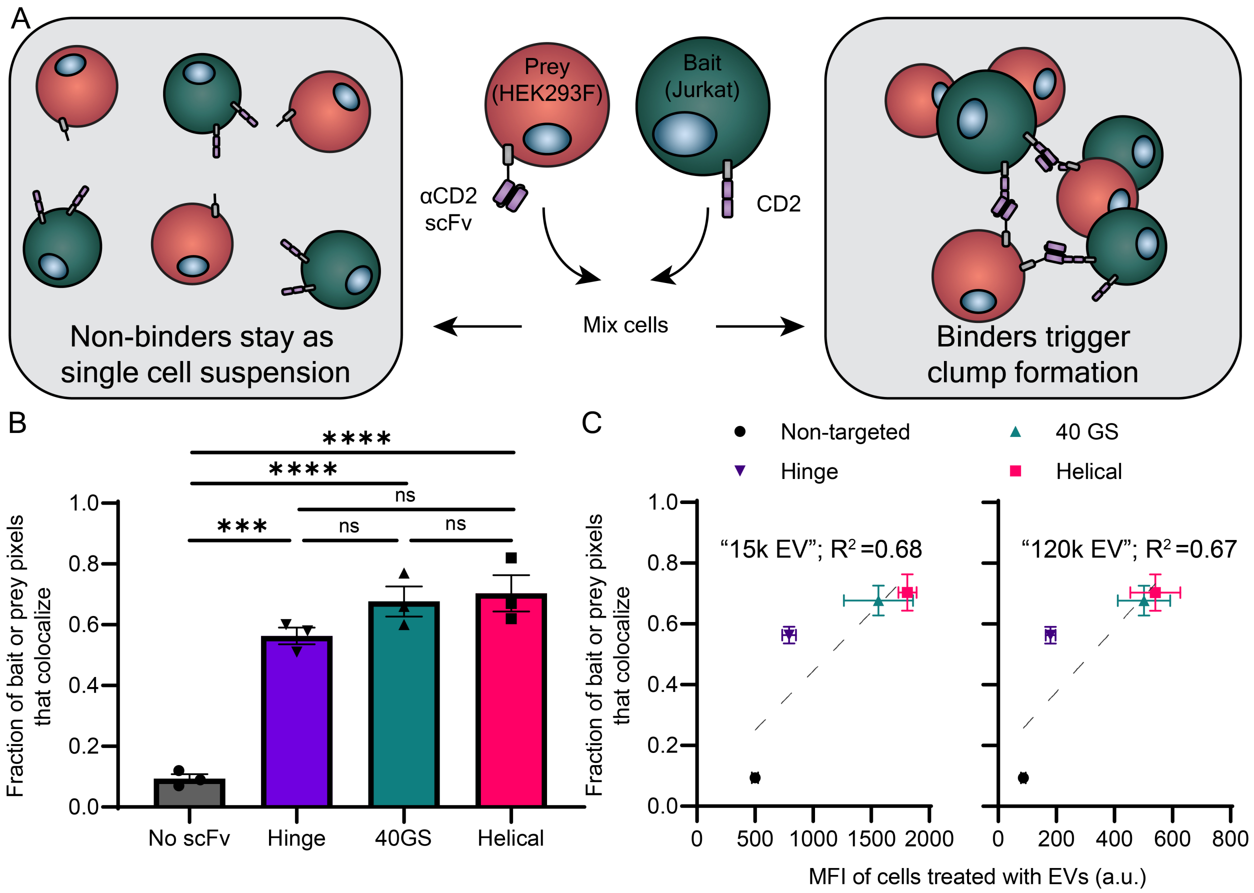


**Figure S11. sCELLISA repeatably recapitulates rank ordering of EV targeting candidates from EV targeting data. (A)** Repeat of the sCELLISA study found in **Figure 3**. **(B)** Quantified sCELLISA data. The fraction of possible pixels that are positive for red and green signals (i.e., that colocalize) indicates interactions between the two cell types. Each symbol represents the average of three fields of view from a biological replicate (n = 3), the bar represents the mean, and the error bars represent SEM. A one-way ANOVA was performed, and the Tukey's multiple comparisons test are shown (****, $p < 0.0001$; ***, $p < 0.001$; ns, $p > 0.05$). **(C)** Comparison of sCELLISA data shown in **(B)** with EV targeting data for these constructs for EVs pelleted at 15,000 g (left, "15k EV", referred to as microvesicles in the original study) and EVs pelleted at 120,000 g (right, "120k EV", referred to as exosomes in the original study) from prior work [1]. Symbols represent the mean of 3 biological replicates for sCELLISA data (y-axis) and 3 biological replicates for EV targeting data (x-axis); error bars represent the SEM. The grey dotted line represents a linear regression drawn through the four points. sCELLISA results highly correlate with EV targeting data ($R^2 > 0.67$ for both vesicle populations) and also correctly predict the rank order of binders.

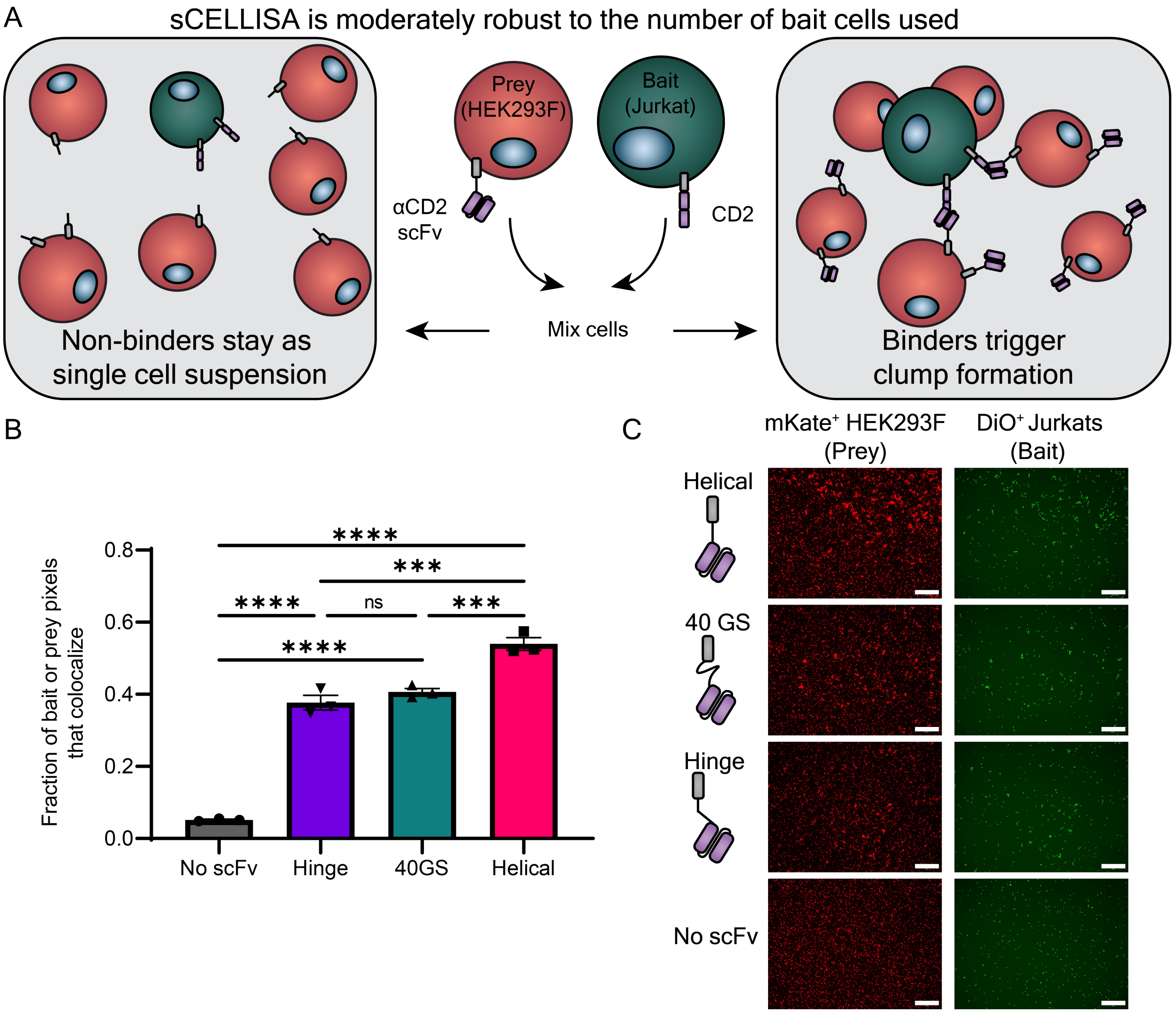


**Figure S12. sCELLISA is modestly robust to the number of prey cells in a given assay. (A)** Schematic of the sCELLISA. An sCELLISA was performed with 10-fold fewer Jurkat bait cells ($10^4$ Jurkats per well) than in other sCELLISA studies (e.g., **Figure 3**) to understand if the sCELLISA methodology was sensitive to the number of bait cells used. **(B)** Quantified sCELLISA data. Under these conditions, the sCELLISA accurately recapitulated the rank order of binders found in **Figure 3**. The y-axis is the fraction of possible pixels that are positive for red and green signals (i.e., that colocalize) and indicates interactions between the two cell types. Each symbol represents the average of two fields of view from a biological replicate (n = 3), the bar represents the mean, and the error bars represent SEM. A one-way ANOVA was performed, and the Tukey's multiple comparisons test are shown (****, $p < 0.0001$; ***, $p < 0.001$; ns, $p > 0.05$). **(C)** Samples micrographs from this study. This experiment was performed once. Scale bar represents 500 μm. Images have been edited for illustrative purposes; the analysis was performed on unmodified images.

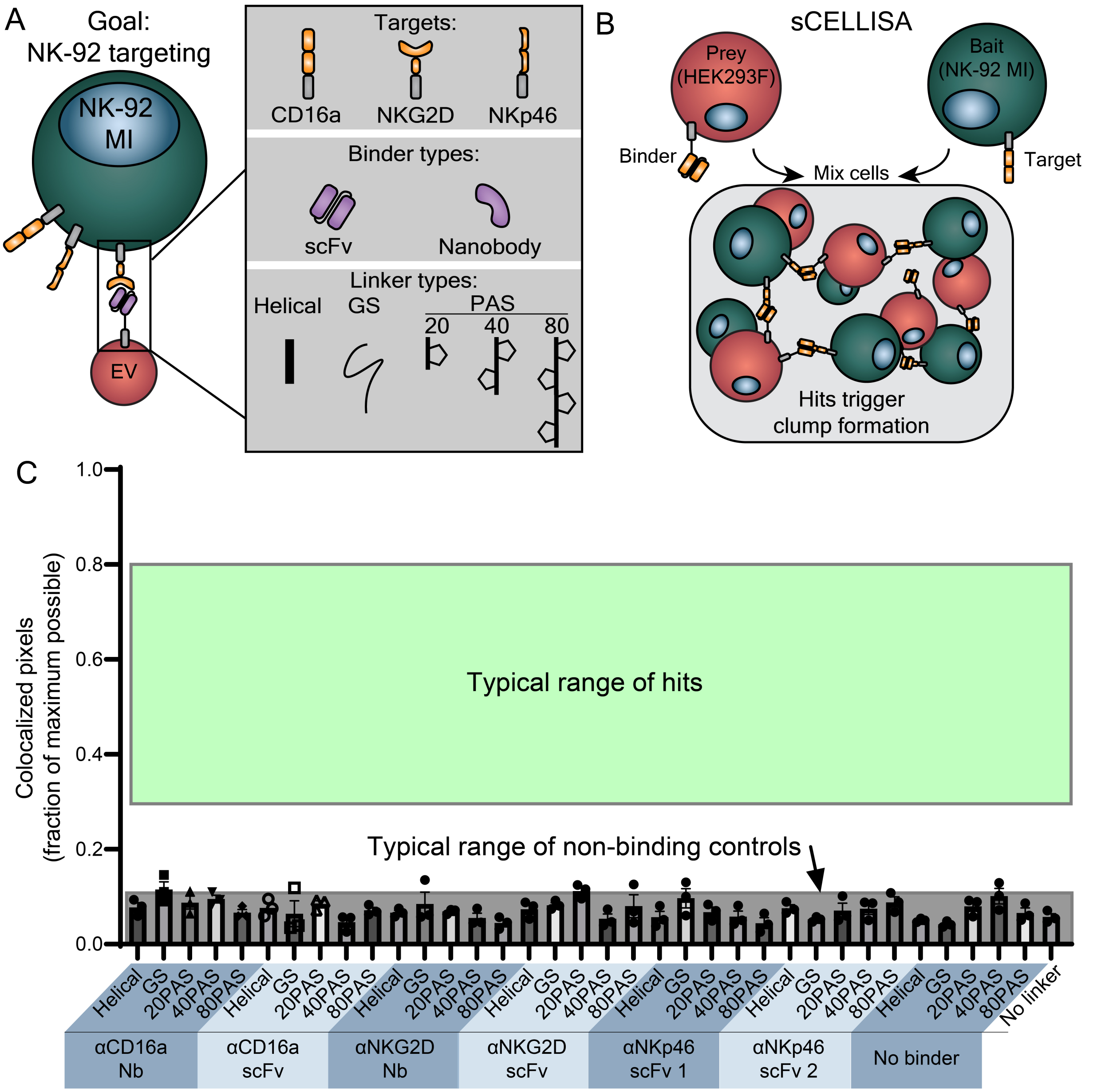


**Figure S13. sCELLISA enabled medium throughput, microscopy-based screening of EV targeting constructs. (A)** Depiction of the design choices evaluated toward building targeting constructs that direct EVs to NK-92 MI cells. In this screen, binding candidates were built that targeted one of three NK surface receptors, via one of two binder classes, linked to a transmembrane anchor using one of five linker types. The anchor was the transmembrane domain region of PDGFR as used in **Figure 2**. **(B)** Illustration of the sCELLISA setup. **(C)** Quantified sCELLISA data. None of the potential designs produced hits in the sCELLISA, suggesting that none of the designs are likely to enable EV targeting to NK-92 MI cells. Shaded regions are visual guides to indicate typical values for non-binding controls or screening hits for Jurkats. Each symbol represents a biological replicate (n = 3), the bar represents the mean, and the error bars represent SEM.

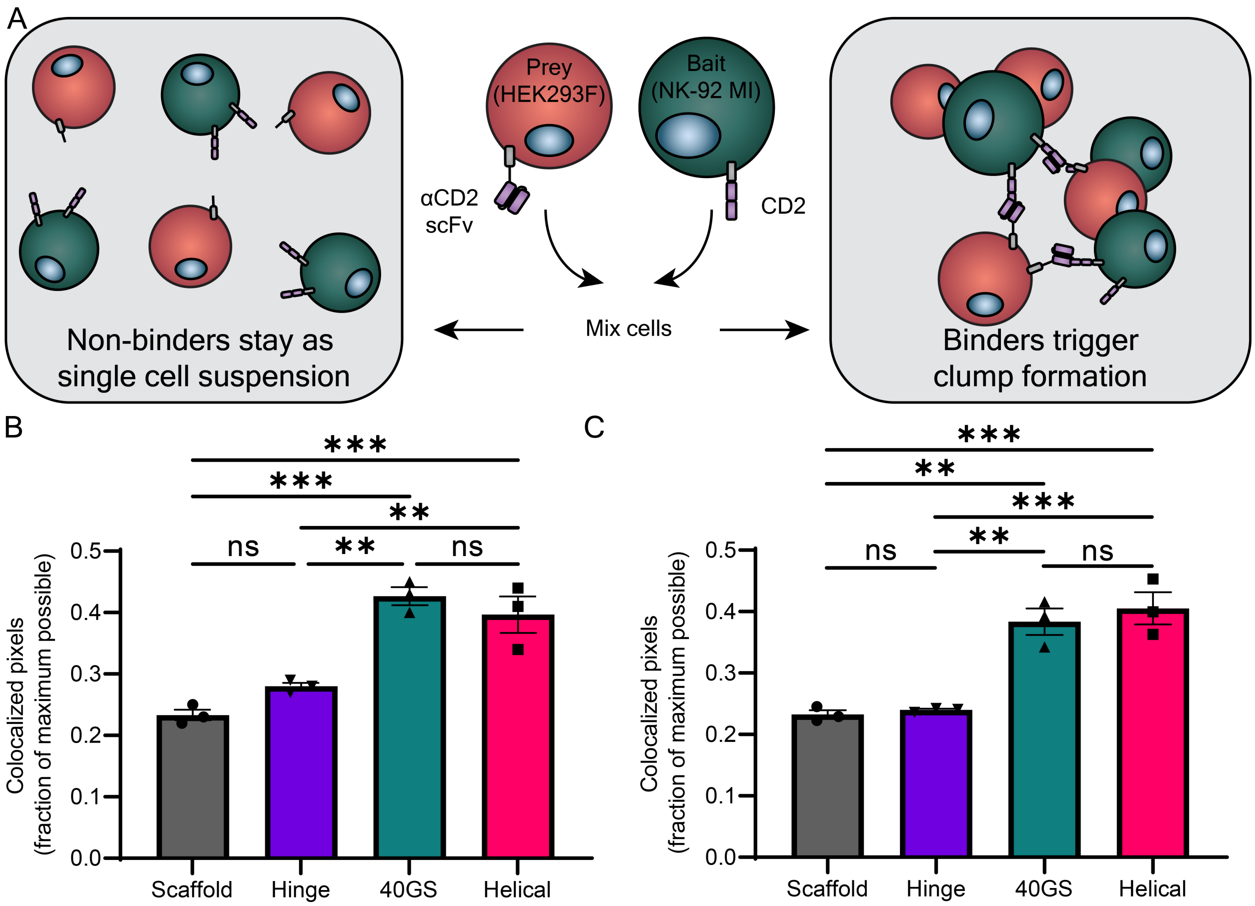


**Figure S14. sCELLISA is compatible with an alternate cell type, NK92 MIs. (A)** Schematic describing the sCELLISA assay setup for this experiment. NK-92MI cells are the bait cells, and HEK293Fs express the αCD2 scFv and linkers from **Figure 2**. NK-92MI cells express CD2 (**Figure S3**). **(B,C)** sCELLISA results for two independent experiments. The fraction of possible pixels that are positive for red and green signals (i.e., that colocalize) indicates interactions between the two cell types. Each symbol represents the average of **(B)** three fields of view or **(C)** two fields of view from a biological replicate (n = 3), the bar represents the mean, and the error bars represent SEM. A one-way ANOVA was performed, and the Tukey's multiple comparisons test are shown (****, $p < 0.0001$; **, $p < 0.01$; ns, $p > 0.05$). Because there is no *a priori* EV targeting data with which to compare these results, these data suggest two of the linker designs, the 40GS and the Helical, are good candidate binders for NK-92 MIs. Importantly, these results demonstrate the sCELLISA assay and analysis is compatible with multiple cell types.

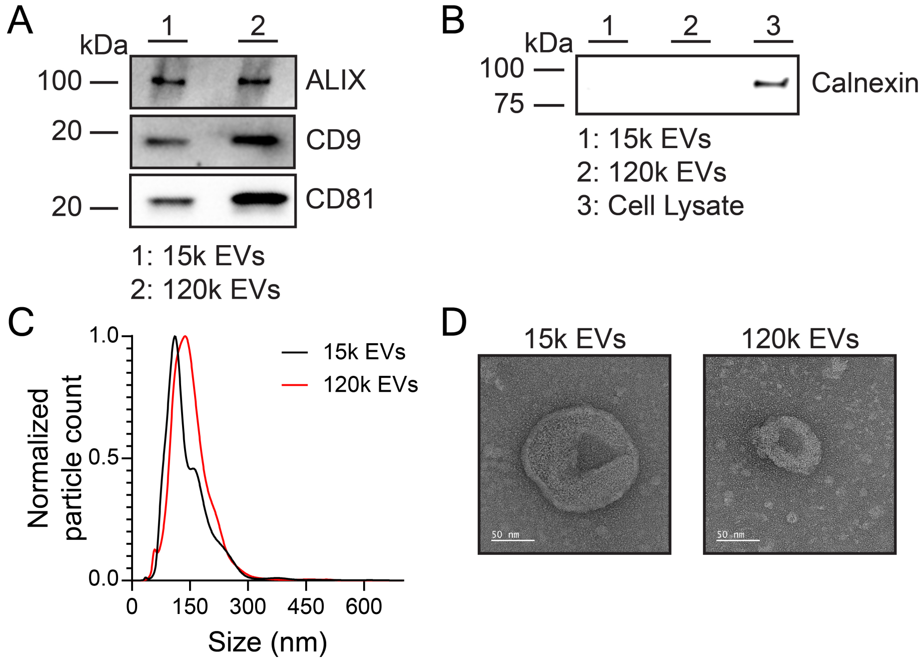


**Figure S15. EV characterization studies. (A,B)** Western blots of EVs and cell lysates. The western blots yielded expected patterns of common EV markers in vesicles versus producer cells. EVs contain the expected markers (ALIX, CD9, and CD81), and calnexin is only present in cell lysate (considered a protein contaminant of EVs). **(C)** Representative histogram of nanoparticle tracking analysis for 15k EVs and 120k EVs derived from Freestyle HEK293F cells. The curves are normalized to the mode in each population. **(D)** EV morphology analysis by transmission electron microscopy. EVs show standard cup-shaped morphology expected for this method. Left: 15k EVs. Right: 120k EVs. Scale bar: 50 nm.

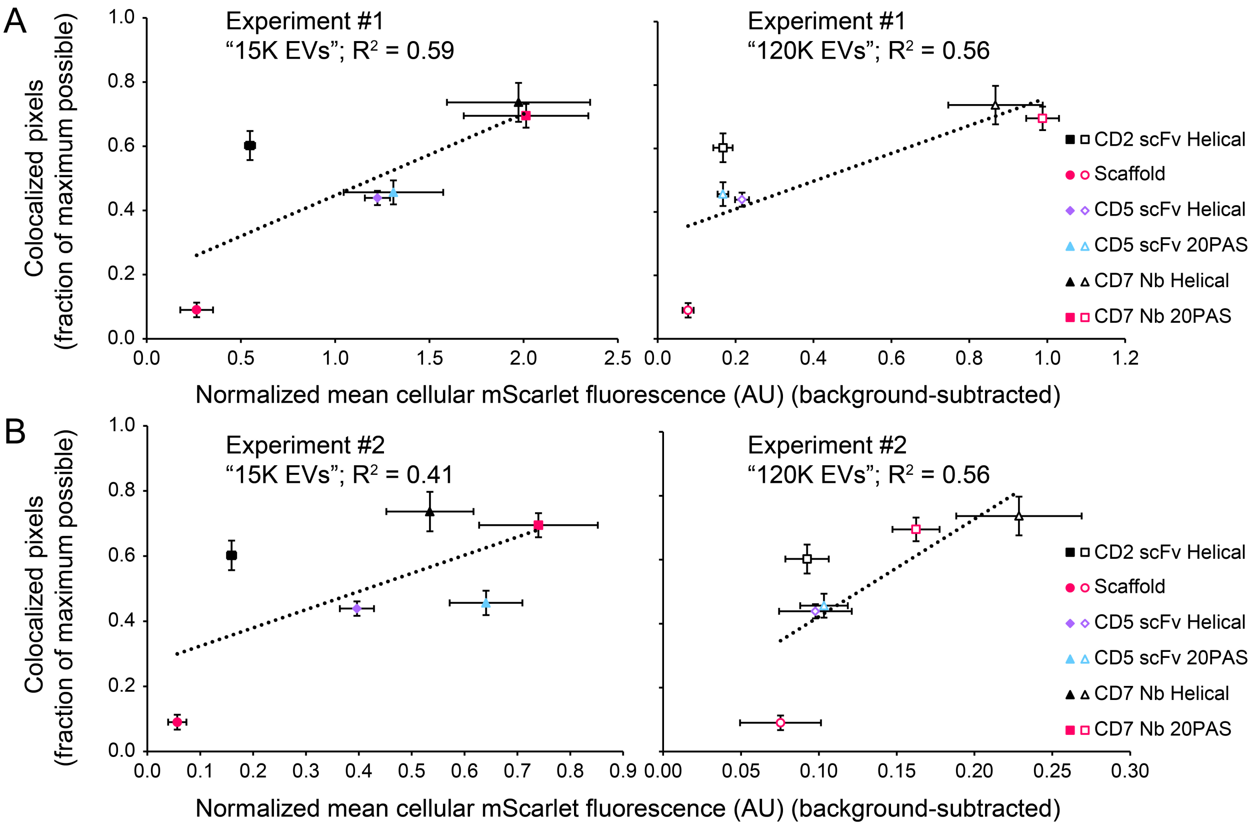


**Figure S16. Correlations between sCELLISA and EV targeting data.** Comparison of sCELLISA data (**Figure 4B**) with EV targeting data (**Figure 4D**) for these constructs for EVs pelleted at 15,000 g (left, "15K EVs") and EVs pelleted at 120,000 g (right, "120k EVs") for EV targeting Experiment 1 **(A)** and Experiment 2 **(B)**. Symbols represent the mean of 3 biological replicates for sCELLISA data (y-axis) and 3 biological replicates for EV targeting data (x-axis); error bars represent the standard error of the mean. The black dotted line represents a linear regression of all six averages. sCELLISA results correlate with EV targeting data ($R^2$ ranges from 0.41 to 0.59 for 15k EVs, $R^2$ is 0.56 for 120k EVs across both experiments).

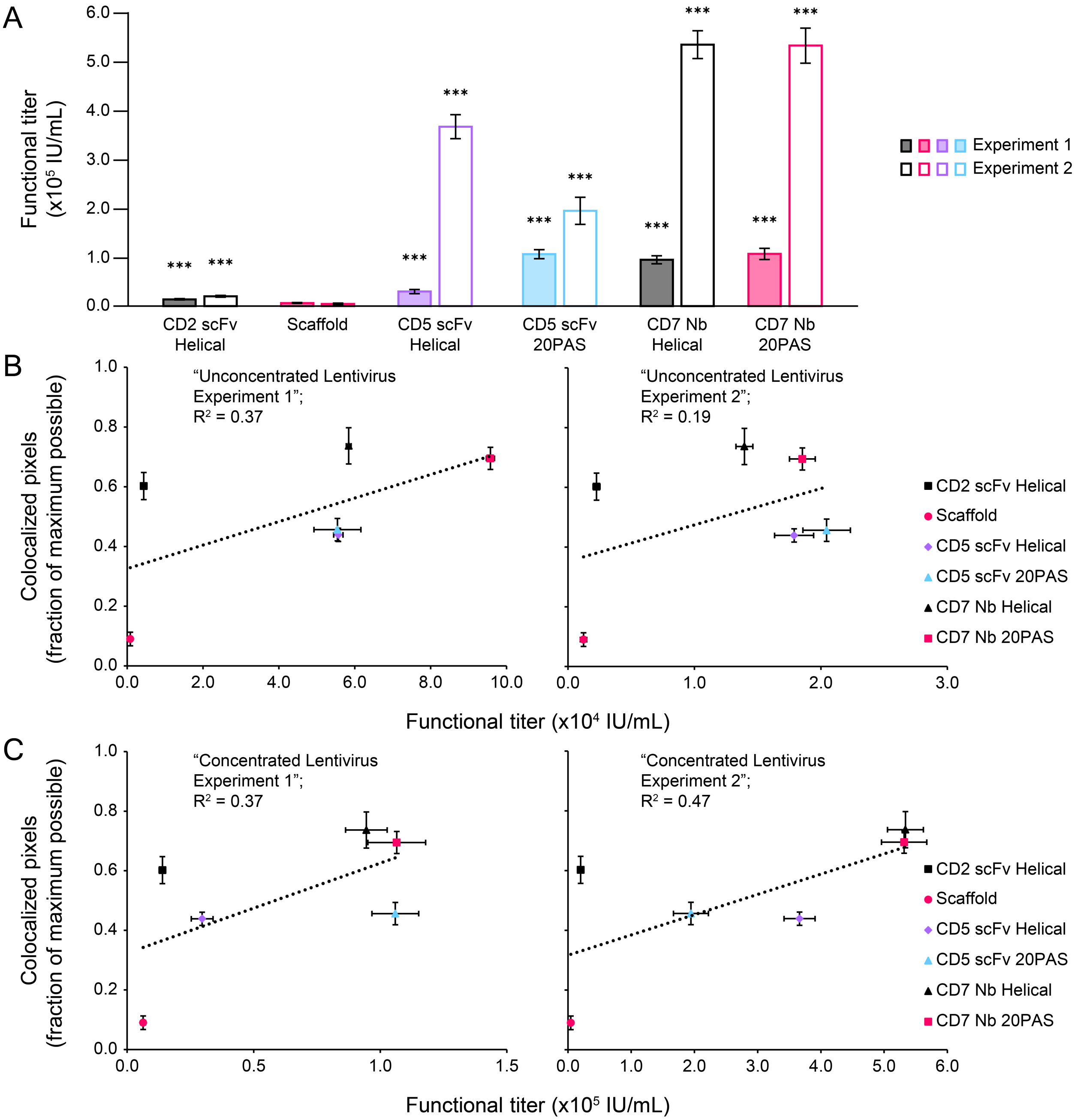


**Figure S17. Concentrated lentivirus functional titers and lentiviral delivery study correlations with sCELLISA data. (A)** Quantification of functional titers of concentrated lentiviruses from cells expressing mutant VSV-G and the binder indicated on the x-axis. Filled and outlined bars each represent an independent experiment. Each bar represents the mean functional titer as determined by linear regression. Error bars represent the standard error of slope of the linear regression. To identify if a binder improved delivery compared to the scaffold negative control, a pairwise comparison of each slope from each linear regression to the slope of the scaffold using a one-sample t-test (***, $p < 0.001$; **, $p < 0.01$; *, $p < 0.05$; ns, $p > 0.05$). **(B** and **C)** Comparison of sCELLISA data (**Figure 4B**) with unconcentrated **(B)** and concentrated **(C)** lentiviral delivery data for these constructs. Symbols represent the mean of 3 biological replicates for sCELLISA data (y-axis), and the functional titer as determined by linear regression analysis of Jurkat transduction data (x-axis); y-error bars represent the standard error of the mean and x-error bars represent the standard error of the regression. The black dotted line represents a linear regression drawn through all six averages. sCELLISA results correlate with unconcentrated lentiviral delivery data ($R^2$ of 0.37 and 0.19 for each independent experiment) and the concentrated lentiviral delivery data ($R^2$ of 0.37 and 0.47 for each independent experiment).

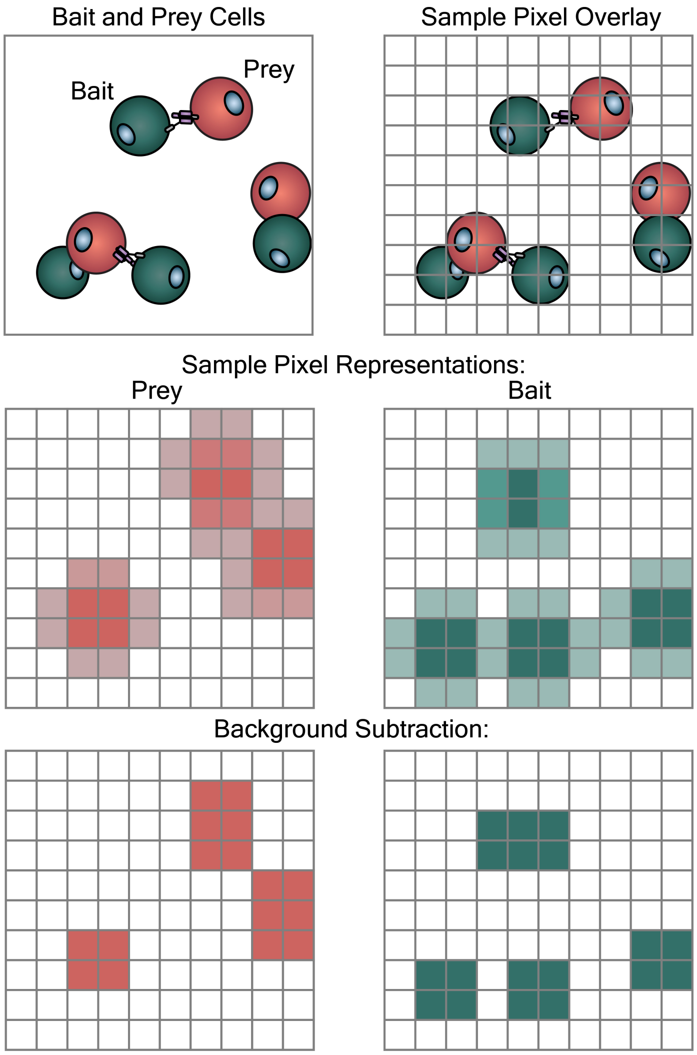


**Figure S18. Illustration of how bait and prey images are represented as pixels before and after background subtraction in the software.**

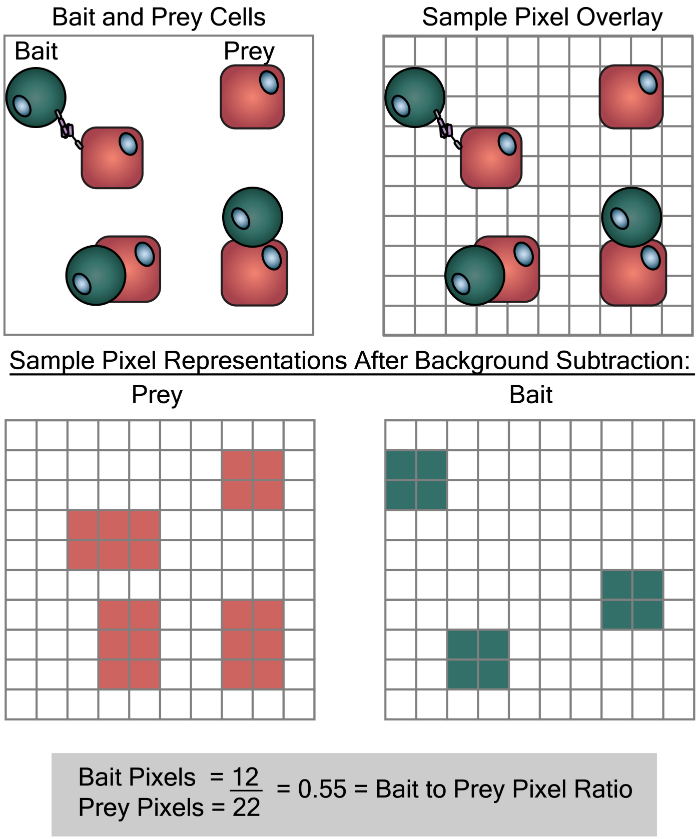


**Figure S19. Illustration of how the software calculates bait to prey pixel ratio for an adherent CELLISA.**

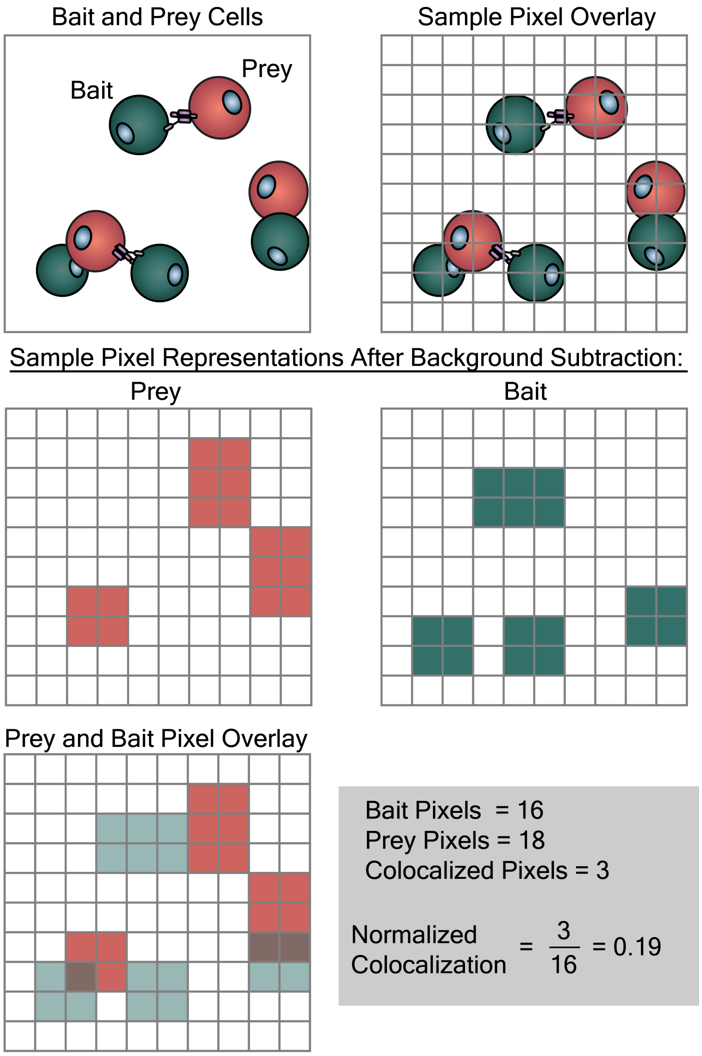


**Figure S20. Illustration of how the software calculates normalized colocalization for a suspension CELLISA.**

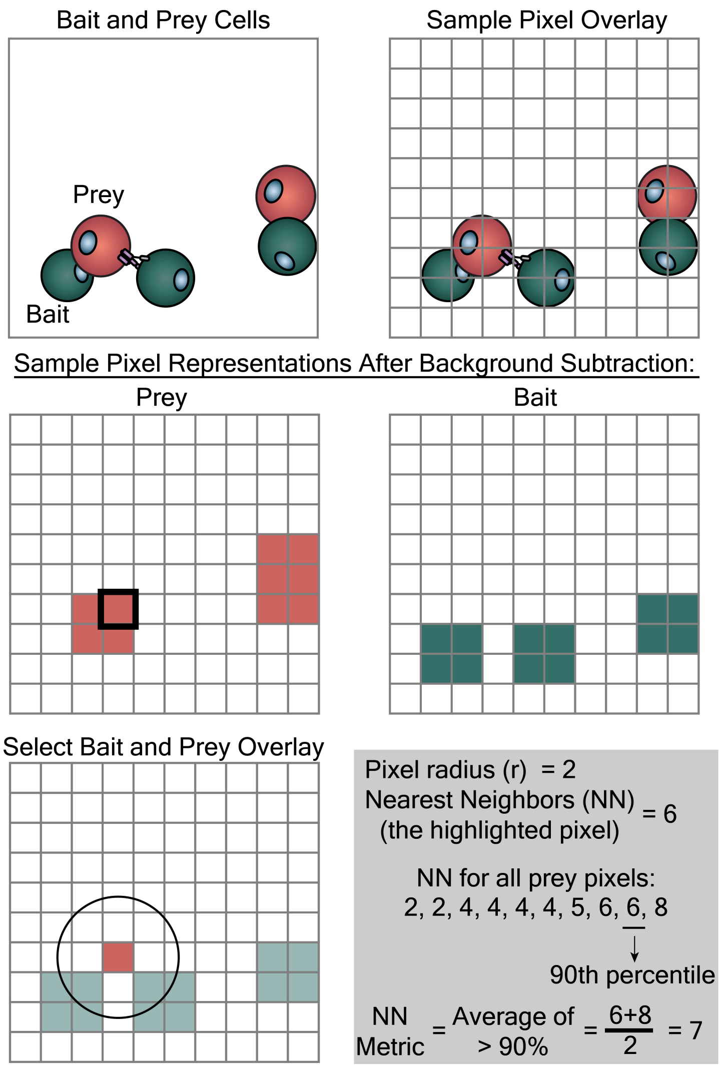


**Figure S21. Illustration of how the software calculates the nearest neighbors metric for a suspension CELLISA.**

### SUPPLEMENTARY TABLES

**Table S1.** Antibodies used for western blot experiments

| Antibody Target | Manufacturer (#) | Denature temperature / time | Antibody dilution | Reducing/Non-reducing Laemmli | Animal of origin |
|---|---|---|---|---|---|
| CD9 | Santa Cruz (sc-13118) | 95°C / 10 min | 1:500 | Reducing | Mouse |
| CD81 | Santa Cruz (sc-23962) | 95°C / 10 min | 1:500 | Non-reducing | Mouse |
| Alix | Abcam (Ab117600) | 95°C / 10 min | 1:500 | Reducing | Mouse |
| Calnexin | Abcam (Ab22595) | 70°C / 10 min | 1:1000 | Reducing | Rabbit |
| Rabbit | Invitrogen (32460) | Not applicable | 1:3000 | Not applicable | Goat |
| Mouse | Cell Signaling Technology (7076) | Not applicable | 1:3000 | Not applicable | Horse |

**Table S2.** Antibodies used for surface staining experiments.

| Antibody Target | Manufacturer | Conjugated fluorophore | Catalog Number | Isotype Control |
|---|---|---|---|---|
| CD2 | Biolegend | Phycoerythrin (PE) | #300207 | Mouse IgG1 kappa |
| CD3 | Biolegend | PE | #300408 | Mouse IgG1 kappa |
| CD5 | Biolegend | PE | #300608 | Mouse IgG1 kappa |
| CD7 | Biolegend | PE | #343106 | Mouse IgG2a kappa |
| NKG2D | Biolegend | PE | #320805 | Mouse IgG1 kappa |
| NKp46 | Biolegend | PE | #331907 | Mouse IgG1 kappa |
| CD16a | Biolegend | PE | #302007 | Mouse IgG1 kappa |
| Isotype control: Mouse IgG1 kappa | Biolegend | PE | #400112 | N/A |
| Isotype control: Mouse IgG2a kappa | Biolegend | PE | #400212 | N/A |
| FLAG | R&D Systems | Alexa Fluor 488 (AF488) | #IC8529G | N/A |

### SUPPLEMENTARY NOTES

#### Supplementary Note 1. CELLISA analysis workflow.

This note is intended as a complement to the software posted to GitHub (see **Appendix A**), with the goal of explaining the algorithmic approach but not superseding the user guide included in that repository.

The general approach is as follows:

1. <u>Upload images</u>: Matched pairs of bait and prey images are loaded into the analysis program.

2. <u>Background subtraction</u>: A region representative of the background fluorescence associated with each image is manually identified by the user (**Figure S18**). The background is selected independently for each of the two images per matched bait and prey pair. The software calculates the background for each image (i.e., in each channel), which is defined as the average intensity of the brightest 20% of pixels within the background region; any pixel brighter than this calculated threshold value was defined as fluorescent. We refer to these as “bait pixels” or “prey pixels.”. This approach was chosen to render the calculated magnitude of background fluorescence less sensitive to the (arbitrary) boundaries of the background region defined by the user; this choice also results in a conservative (high) definition of background fluorescence magnitude, which seemed to make it easier for users to find an appropriate background.

3. <u>Define area of analysis</u>: Prior to image processing, regions considered inappropriate for calculating CELLISA metrics are manually curated. Thus, in some cases in this study, an area exclusion tool was utilized to remove regions such as the edge of a well, where optical effects and/or variable well density made these regions poor representations of the overall image (e.g., cells could apparently co-localize due to high density near the edge of the well, and that is expected to have no physical meaning).

4. <u>Calculate bait-prey interaction metrics:</u>

   **aCELLISA**: For this type of assay, the relevant interaction metric was defined as the ratio (i.e., quotient) of the number of fluorescent bait pixels divided by the number of fluorescent prey pixels within a given region of analysis (**Figure S19)**. This metric reflects the degree to which bait cells in suspension are recruited to and retained on the surface (after washing), normalized by the abundance of (adherent) prey cells (which could vary from well to well or region to region). This ratio was calculated for each matched pair of bait and prey images and repeated for all samples in an experiment.

   **sCELLISA**: For this type of assay, we evaluated several potential metrics to determine which is most appropriate, and a comparison of those approaches is presented in **Figure S10** and the accompanying caption. We considered two general metrics for quantifying bait-prey interactions: (1) *Normalized colocalization* and (2) *Nearest neighbors*.

   To calculate *Normalized colocalization,* within the region of analysis for each pair of images (bait and prey images), the software first calculated the number of pixel locations that were positive for *both* bait and prey fluorescence (i.e., colocalized) (**Figure S20**). To account for differences in the number of cells present in a given field of view, these raw colocalized pixel counts were then normalized to the number of bait or prey pixels (whichever is smaller). This normalization method was chosen such that a "normalized colocalization" value of 1 would represent the theoretical maximum of this metric. To illustrate, in a case where the number of prey pixels exceeds the number of bait pixels, the maximum number of colocalized pixels is by definition the number of bait pixels, and thus the maximum value of the normalized colocalization metric is 1.

   We explored the *Nearest neighbors* metric as an alternative to the normalized colocalization metric because we hypothesized that the former could better capture cases where bait and prey cells were bound to one another but were not overlapping in the x-y plane, as viewed by the microscope. To calculate the nearest neighbors metric, for each matched pair of bait and prey images, the software first calculated the number of bait pixels "nearby" to each prey pixel (**Figure S21)**. Here, nearby pixels are defined as those bait pixels which are located within a circle, with radius "r", centered around each prey pixel (**Figure S10C,** top). The software then calculated the population-level nearest neighbors metric by averaging the "number of bait pixels nearby" for only those prey pixels falling above the $90^{th}$ percentile (i.e., falling within the top 10%) in the distribution of values for "number of bait pixels nearby the prey pixel" (**Figure S10C**, top). This thresholding method focuses our metric on the most highly clustered regions of an image, serving as a filter to minimize the contribution of random colocalization of bait and prey cells (i.e., pixels) in a way that is internally scaled to the features of each image.

As described in the caption and results presented in **Figure S10,** we found that the normalized colocalization metric yielded a high dynamic range (i.e., enabled differentiation of performance measured across binder designs), and so we used this metric going forward. However, since such a scenario may differ across other uses of the CELLISA method, we include nearest neighbors calculations as an option within the software program (**Appendix A**).

5. Export: all values are exported as a CSV file for subsequent analysis.
