## Supplementary figures and images for "CELLISA – a cell-cell binding assay for evaluation of nanovesicle targeting proteins"

### 25.05.11_13.33.56_Calnexin.tif

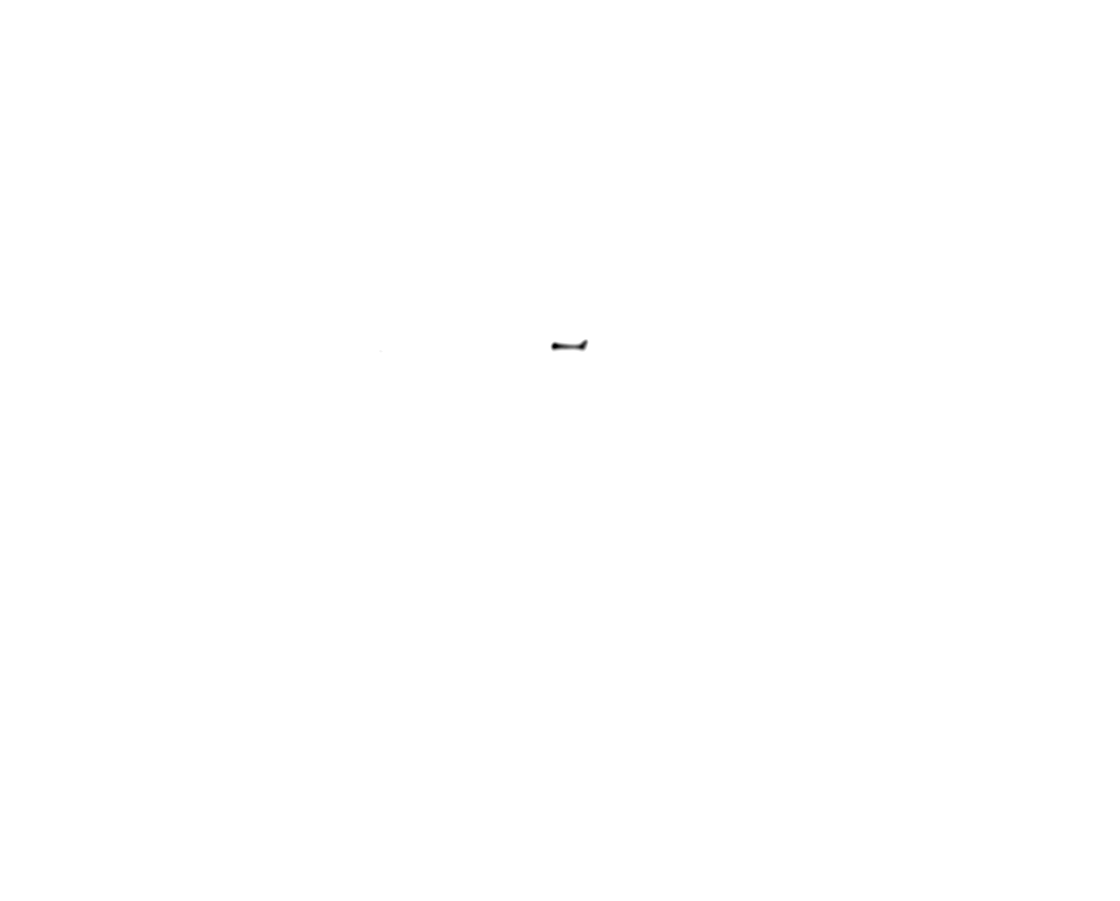

### 25.05.11_13.33.56_marker_PUB_600_Calnexin.tif

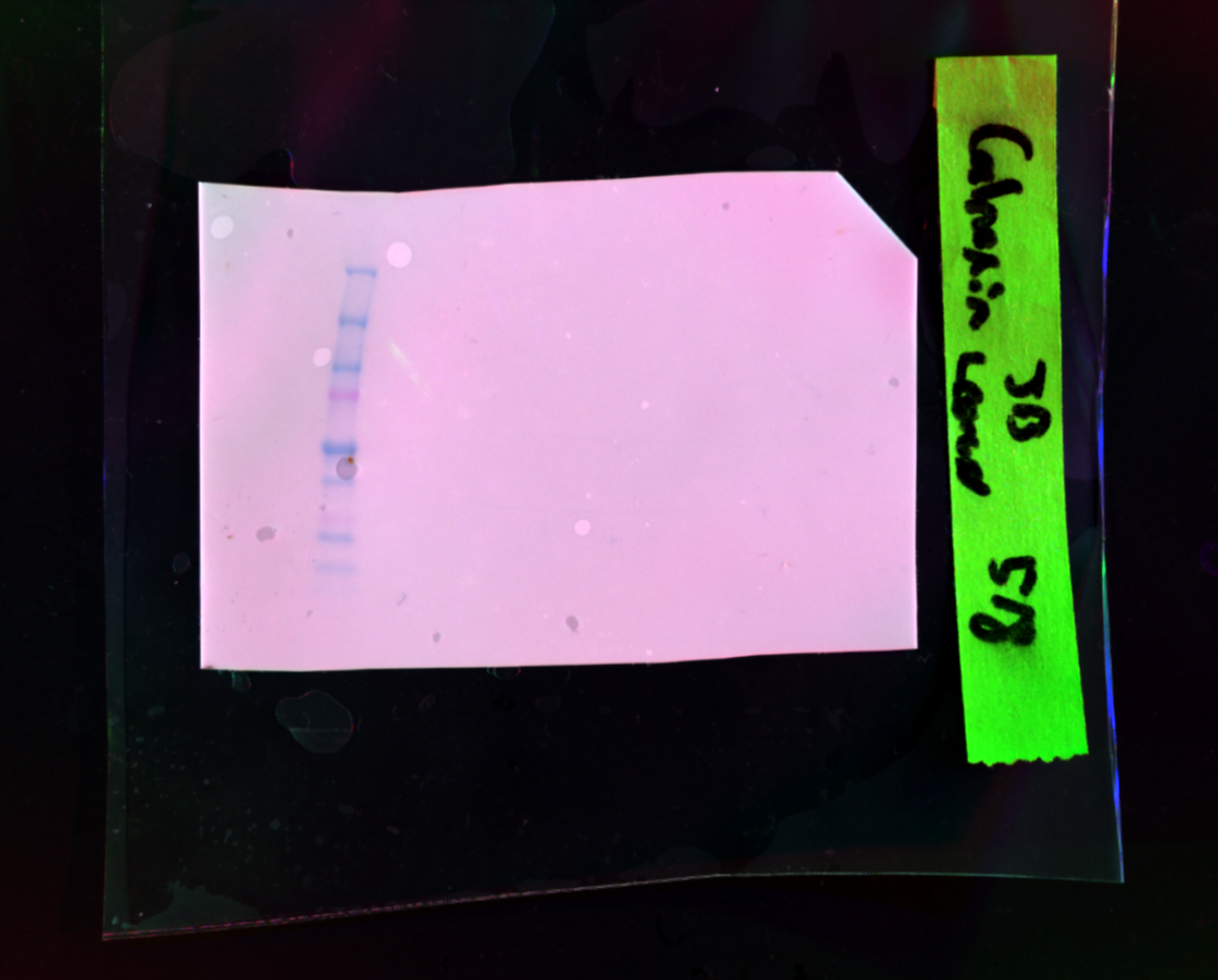

### 25.06.05_22.14.41_marker_PUB_600_ALIX.tif

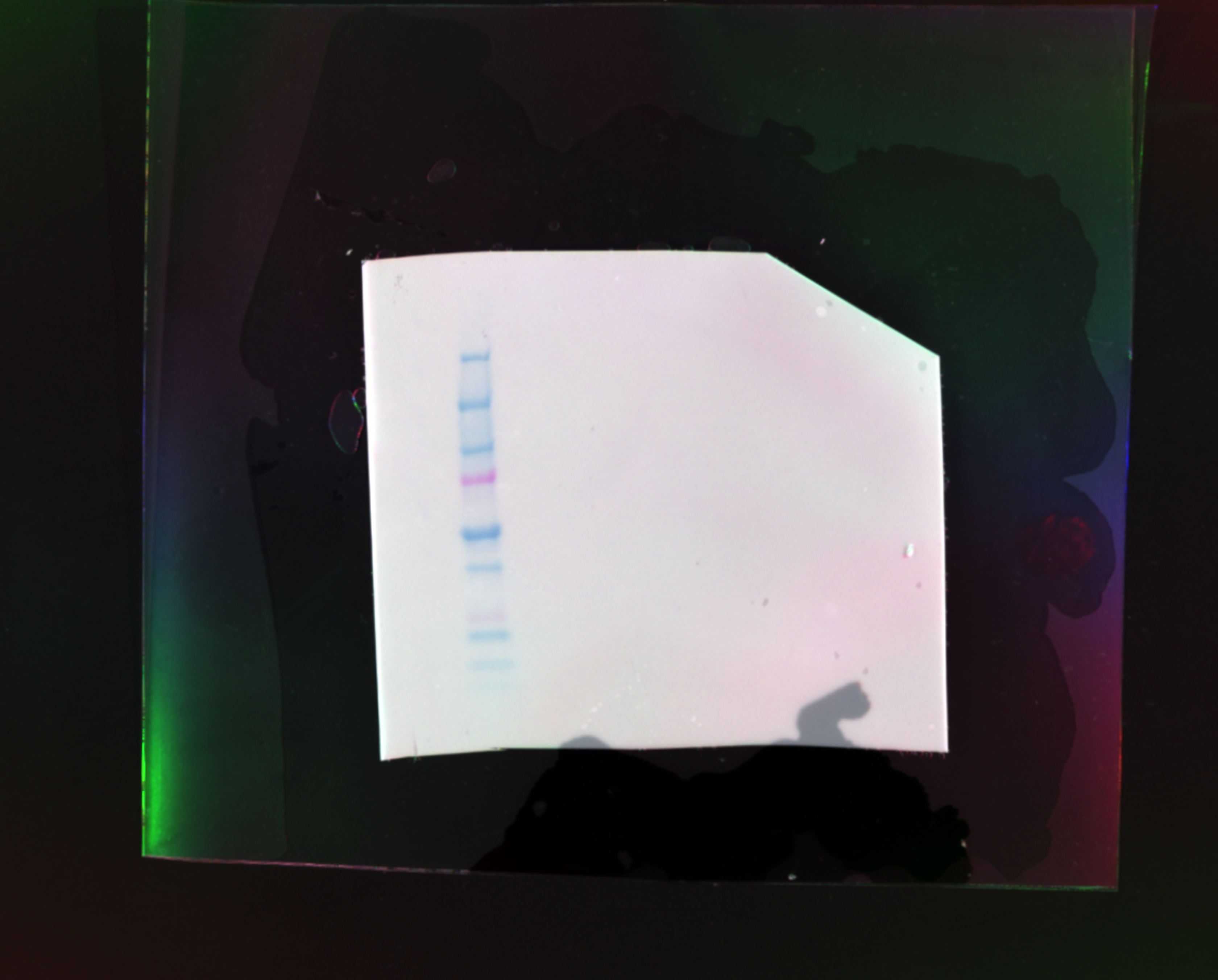

### 25.06.05_22.14.41_PUB_600_ALIX.tif

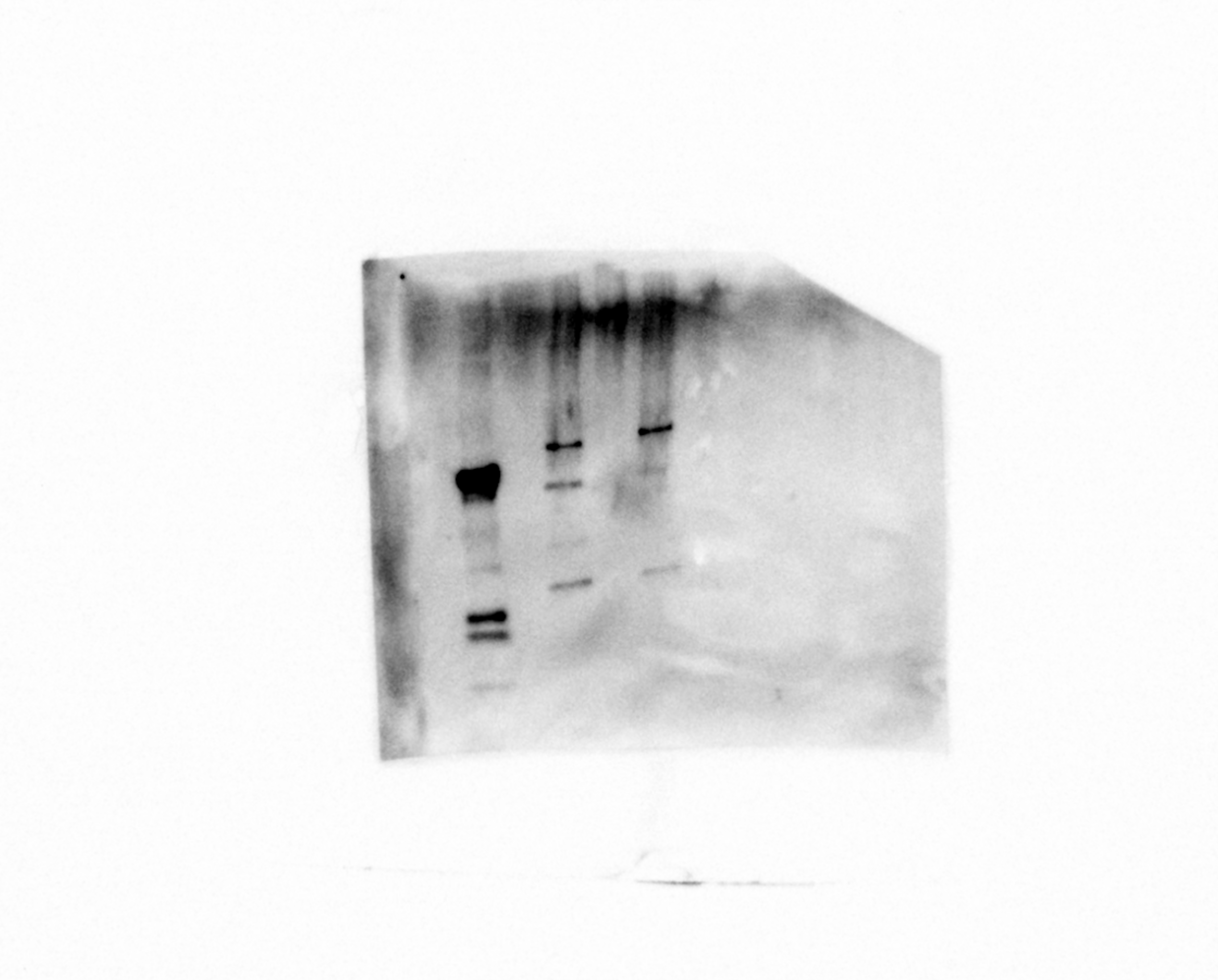

### 25.06.07_20.42.00_S2_F10_PUB_600_CD9.tif

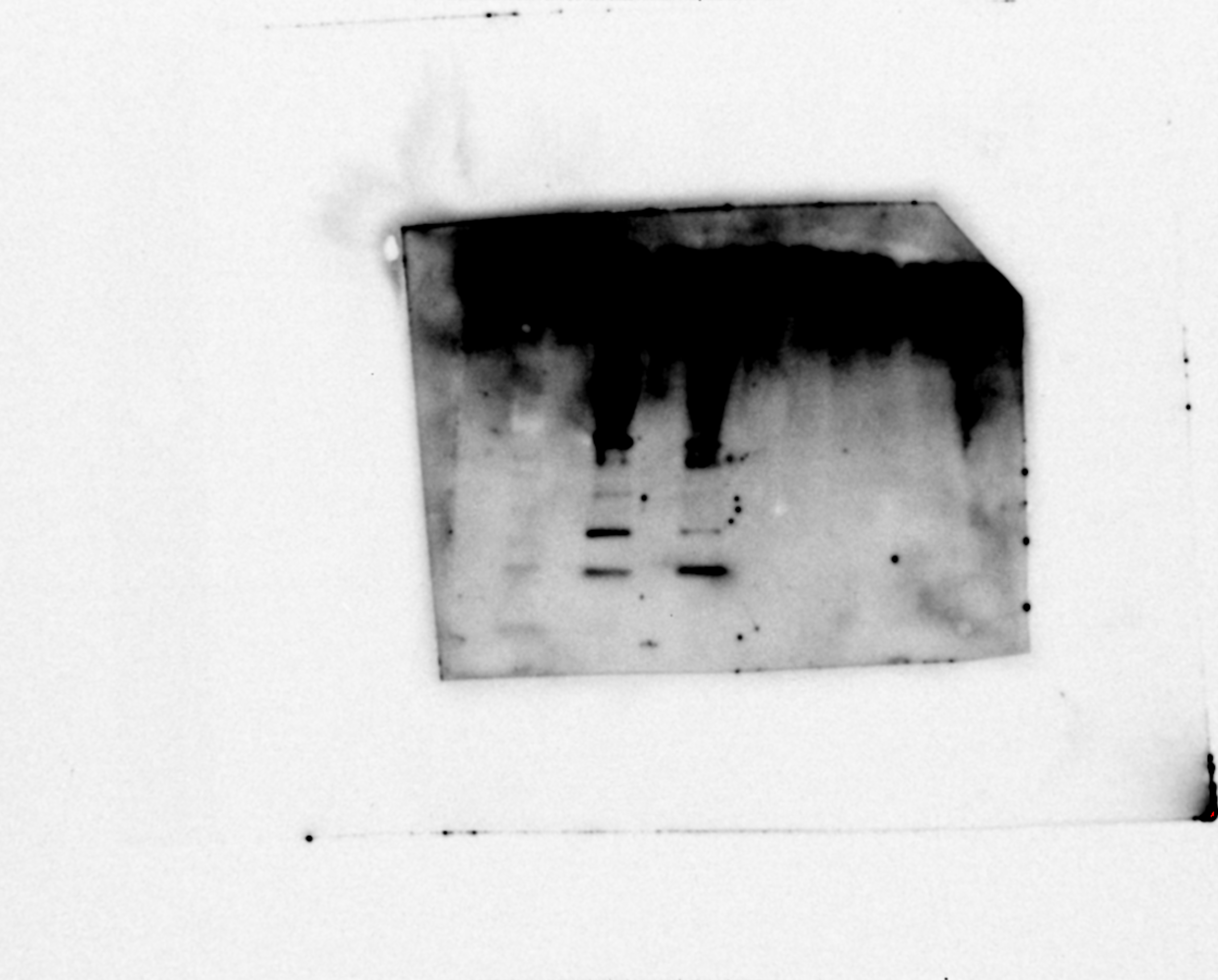

### 25.06.07_20.42.00_S2_marker_PUB_600_CD9.tif

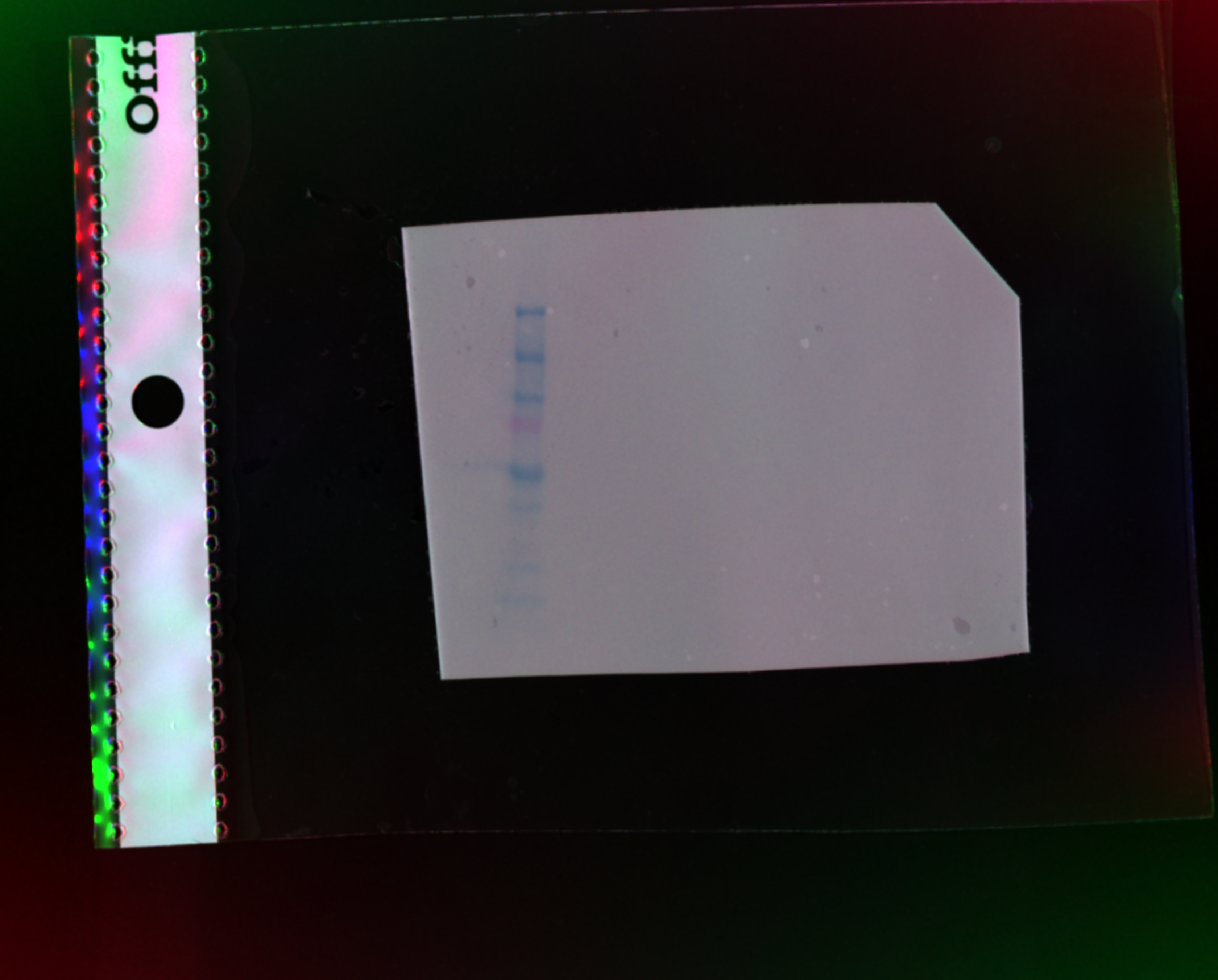

### 25.06.07_21.47.32_marker_PUB_600_CD81.tif

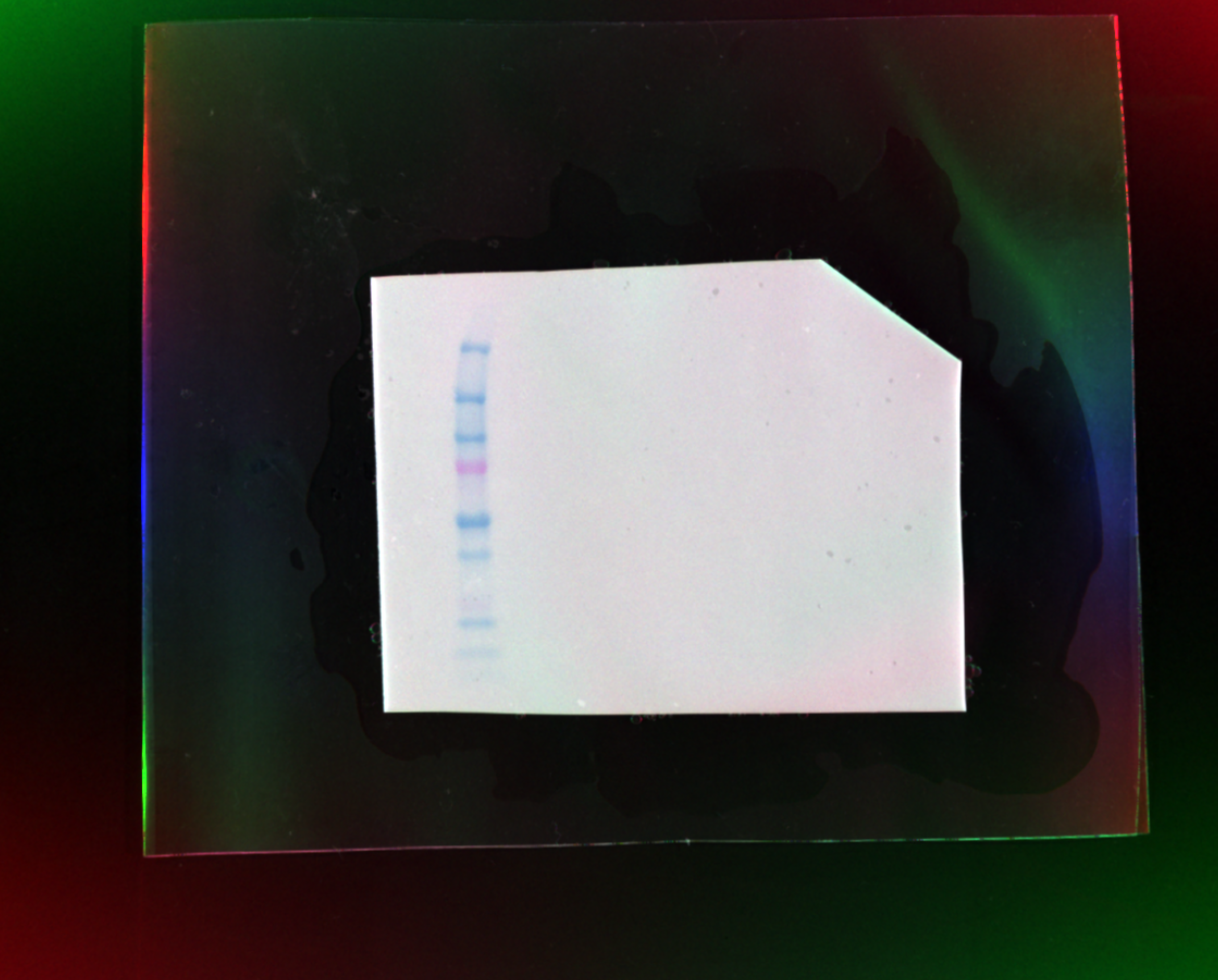

### 25.06.07_21.48.59_S3_F03_PUB_600_CD81.tif

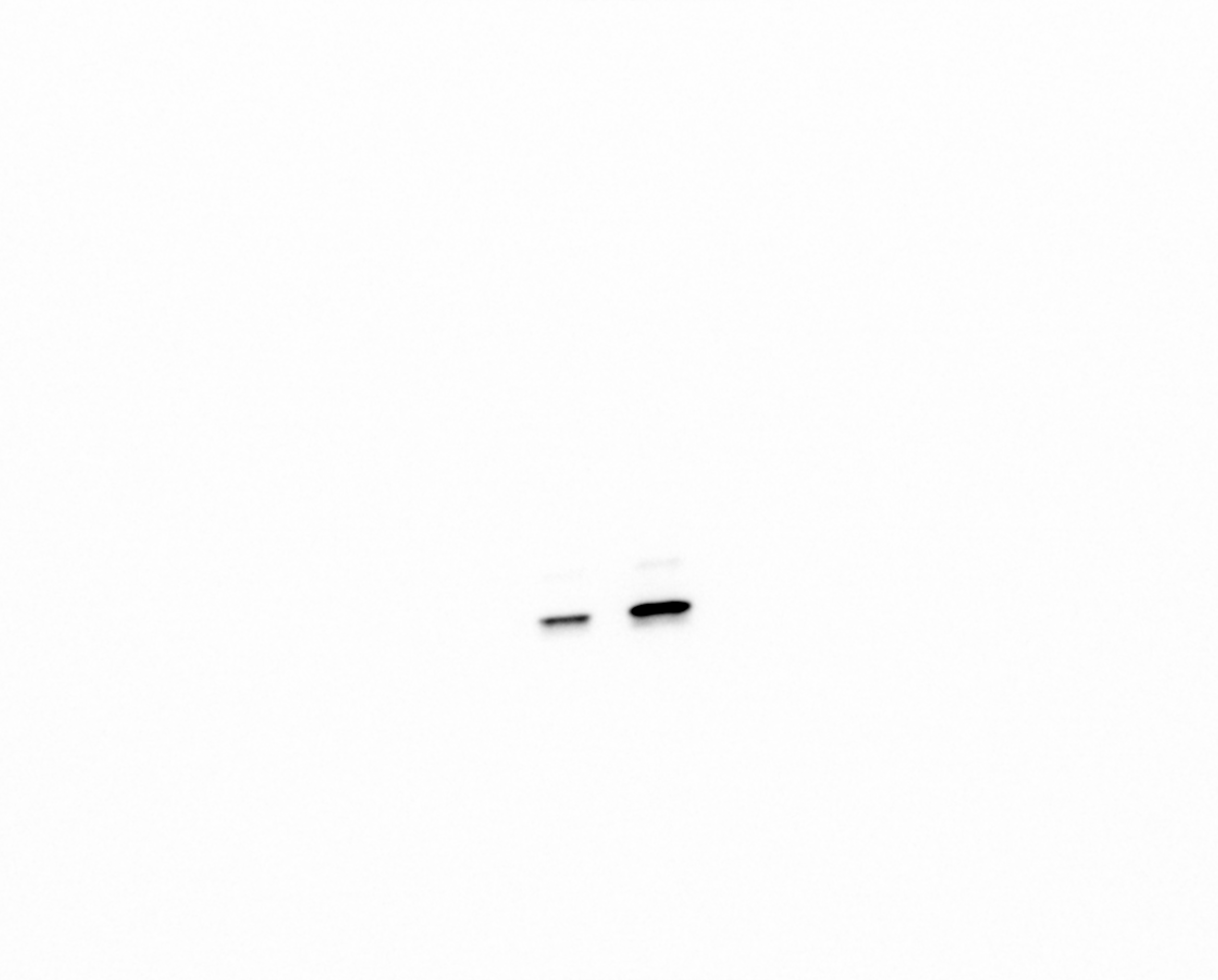

### TEM_CD2 scFv_15k.tif

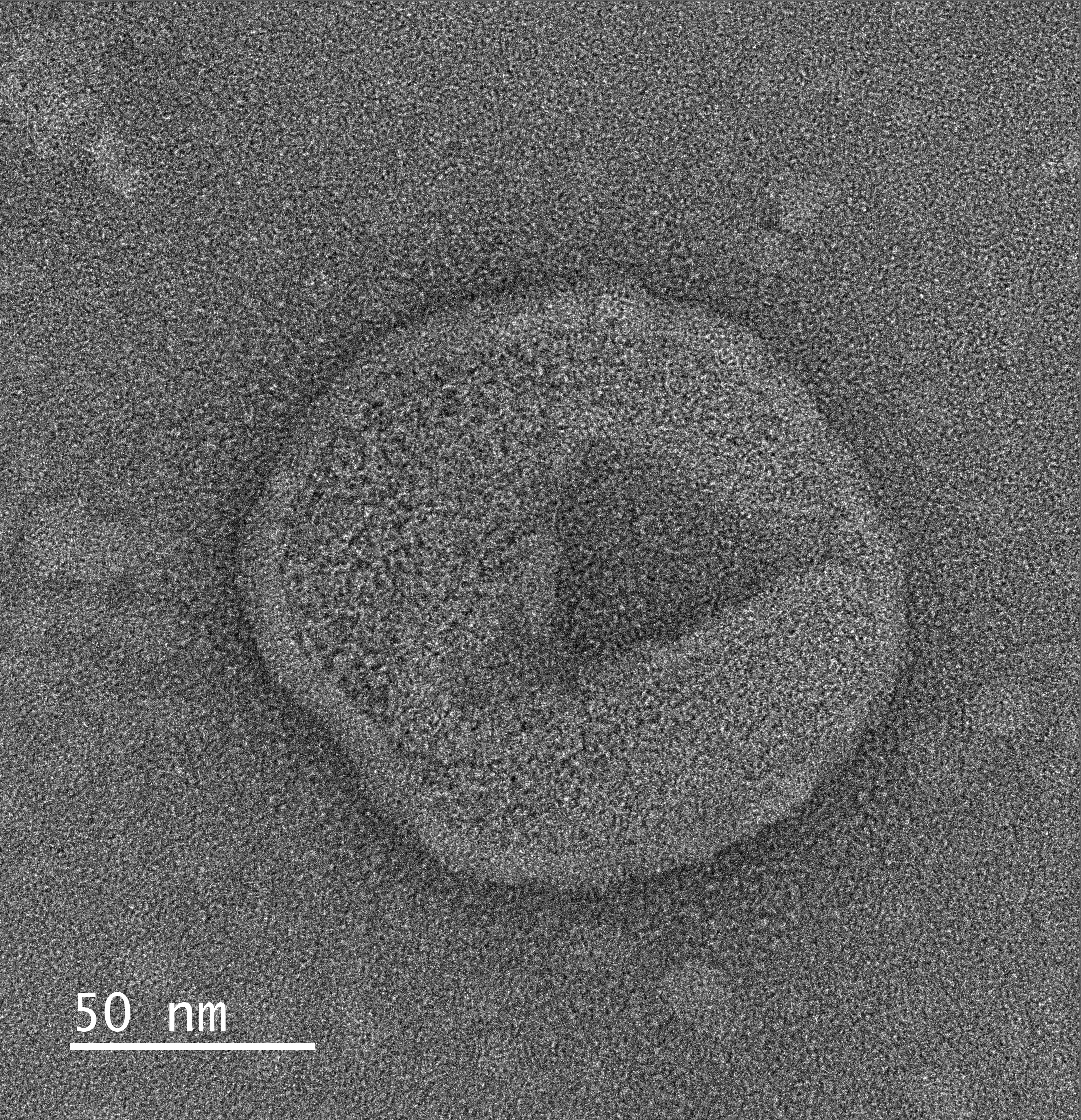

### TEM_CD2 scFv_120k.tif

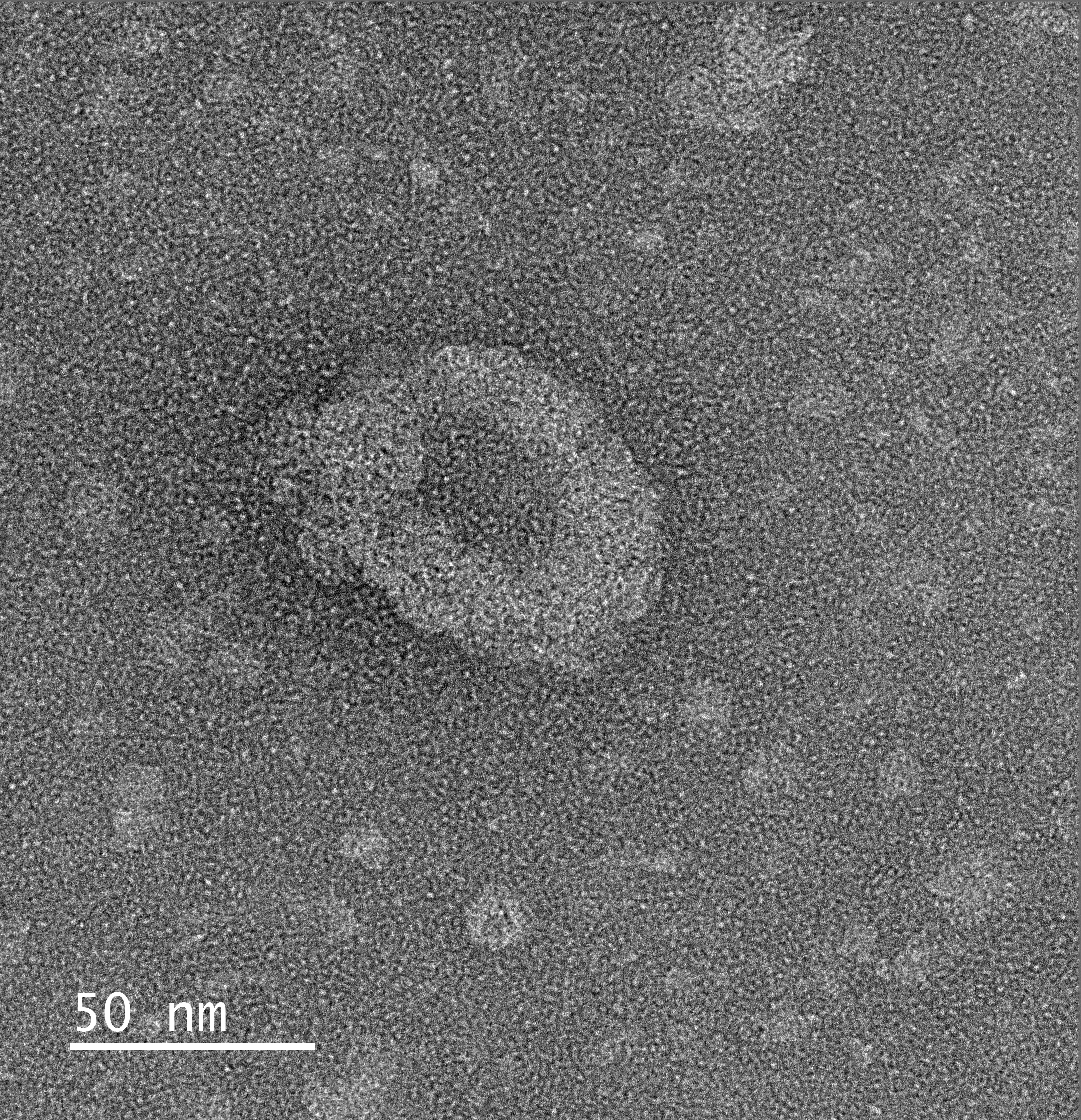
